# Persistently activated virus-specific CD8 T cells indicate antigen-driven immune aberrations in long COVID

**DOI:** 10.64898/2026.10.14.759221

**Authors:** Mark M. Painter, David B. Reeg, Naomi R. Douek, Sophie L. Gray-Gaillard, Sasikanth Manne, Josephine R. Giles, Thomas Dalhuisen, Melissa Buitrago, Emily A. Fehrman, Dylan Ryder, Meghann C. Williams, Ava-Kathleen Rybicki, Gwen Baraniecki-Zwil, Zina Ahn, John P. Reilly, Danielle K. Sandsmark, Michael O. Harhay, Sasha Tabachnikova, Bornali Bhattacharjee, David Putrino, Amy D. Proal, J. Daniel Kelly, Jeffrey N. Martin, Benjamin A. Abramoff, Nuala J. Meyer, Akiko Iwasaki, Timothy J. Henrich, Steven G. Deeks, Michael J. Peluso, E. John Wherry

## Abstract

Long COVID (LC) affects millions of people worldwide, yet its driving mechanisms remain incompletely defined and no treatment has been approved. Persistence of SARS-CoV-2 and reactivation of herpesviruses have been implicated in LC, but direct detection of these viral antigens in blood and tissues has been challenging. T cells are exquisitely sensitive to viral antigens, survey organs throughout the body, and can be readily sampled in peripheral blood. Using high-dimensional spectral flow cytometry and single-cell sequencing approaches with multiplexed peptide/HLA tetramers, we profiled virus-specific T cells from hundreds of people with LC and recovered controls. We identified a subset of people with LC that maintained elevated activation of CD8 T cells targeting SARS-CoV-2 spike and/or lytic antigens from Epstein-Barr virus, but not other viral specificities. Spike-specific cells in LC expressed genes linked to TCR stimulation, terminal effector differentiation, and exhaustion and demonstrated reduced functionality upon cognate peptide stimulation *ex vivo*. Thus, T cells from a subset of people with LC are likely chronically stimulated by persistent SARS-CoV-2 antigens. These findings establish the use of T cells as biosensors of persisting antigen in LC and other disease settings and may help develop new therapeutic strategies for LC.

## Introduction

Long COVID (LC) is an infection associated chronic illness characterized by a broad constellation of symptoms ranging from mild to debilitating that continue for at least three months after SARS-CoV-2 infection(*1*). Symptoms persist for years with no improvement in many individuals, and no effective treatment has been approved(*2, 3*). The biological mechanisms driving disease remain undefined, hindering the development of effective therapies(*4*). Several mechanistic hypotheses have emerged, including persistence of SARS-CoV-2(*5–8*), reactivation of common latent viruses such as Epstein-Barr virus (EBV)(*9–12*), residual tissue damage in sensitive locations (*13, 14*), and new onset autoimmunity(*15*), among others. The heterogeneous nature of LC suggests that multiple distinct mechanisms may drive disease in different subsets of patients, but accessible and sensitive biomarkers are needed to link specific mechanisms to disease in individual patients.

Emerging evidence suggests that some individuals may harbor a persistent tissue reservoir of SARS-CoV-2. Independent studies have detected SARS-CoV-2 RNA or protein in blood(*16–20*), stool(*21, 22*), and in various tissues including the gut(*6, 21, 23–28*), brain(*5, 23*), tonsils and adenoids(*29, 30*), and nasal mucosa(*31*). However, detection alone does not establish a connection to LC, since SARS-CoV-2 RNA or protein has also been found in fully recovered individuals. Viral persistence nonetheless remains a leading mechanistic hypothesis in LC, as persisting SARS-CoV-2 could drive inflammation or viral pathology, explaining chronic symptoms and providing a target for therapeutic intervention(*32*). Direct detection of the tissue-based SARS-CoV-2 reservoir requires sophisticated methods and invasive sampling to observe relatively weak signals in only a fraction of individuals. Thus there remains an urgent need to develop sensitive and accessible approaches to assess viral persistence and to apply them at scale to determine whether SARS-CoV-2 persistence is linked to LC.

Chronic antigen exposure in LC may not derive from SARS-CoV-2 alone. Herpesviruses such as EBV, cytomegalovirus (CMV), and varicella zoster virus (VZV) establish persistent infections that are typically well-controlled by the immune system in healthy individuals, but can reactivate in response to stress, aging, infection, or stimulation of the host cell(*33–35*). Herpesvirus reactivation is frequently observed during acute SARS-CoV-2 infection(*36*), and the symptom profile of LC overlaps substantially with that of EBV infection(*37*). Elevated herpesvirus activity as measured by serological assays has been linked to the development of LC and has been observed most consistently with EBV, but also with CMV and VZV (*9–12, 38*). Reactivation of these viruses may therefore contribute to chronic symptoms, although it remains unknown whether herpesviruses reactivate transiently during acute infection or sustain lytic replication in people with LC.

A large and diverse population of T cells surveys most tissues in the body, with each T cell bearing a unique T cell receptor (TCR) capable of recognizing a specific peptide/HLA antigen. T cells are exquisitely sensitive to viral antigens and can be activated by a single molecule of peptide/HLA(*39, 40*). Upon antigen stimulation, memory T cells differentiate into effector cells, upregulate activation markers, and can re-enter circulation bearing these hallmarks of recent activation(*41, 42*). In the context of chronic infection, antigen-specific T cells can become exhausted, a distinct state of differentiation that shares some features with acute activation(*43*). Circulating virus-specific T cells isolated from peripheral blood can therefore be used as sensitive and specific biosensors of transient or chronic viral antigen expression in tissues throughout the body. The vast majority of circulating T cells, however, do not recognize any individual antigen of interest. Thus, to interrogate whether a particular antigen is driving T cell activation, sensitive methods to isolate and analyze the rare circulating T cells with the desired specificity are needed.

HLA-I and HLA-II tetramers loaded with viral peptides can be used to sensitively identify and/or isolate peptide-specific T cells without stimulation, preserving their *in vivo* activation state (*44, 45*). Small studies have used HLA-I tetramers to profile SARS-CoV-2-specific CD8 T cells in LC, including a recent study of 10 people with LC that reported an increase in cytotoxicity(*46–48*). Cytotoxicity, however, is also a common feature of spike-specific memory CD8 T cells in healthy individuals and is not a clear indicator of recent activation(*49–53*). *In vitro* peptide stimulation has also been used to identify antigen-specific T cells in LC (*48, 54*), but such approaches obscure the *in vivo* activation state of these cells. Thus, larger studies interrogating the activation and differentiation states of virus-specific T cells directly *ex vivo* are needed to determine whether persistent SARS-CoV-2 antigens activate SARS-CoV-2-specific T cells in people with LC. Such efforts have the potential to identify distinct virally-driven endotypes of LC and advance our understanding of the mechanisms regulating disease.

## Results

### Spectral flow cytometry identifies distinct differentiation states for virus-specific T cells

We hypothesized that T cells could be exploited to test whether ongoing immune stimulation by specific viruses is associated with LC. We therefore developed a high-dimensional spectral flow cytometry approach using multiplexed peptide/HLA tetramers to identify and interrogate rare virus-specific T cell populations and test whether these natural biosensors showed evidence of recent stimulation with viral antigens. In total, we analyzed over 500 peripheral blood mononuclear cell samples collected from over 300 individuals across multiple cohorts, comparing T cell responses in over 150 people with LC to those observed in recovered participants without LC, during acute SARS-CoV-2 infection, and less than 30 days after SARS-CoV-2 vaccination (acute vax) (**Fig. 1A** and **Tables 1-2**). We assessed all participants for interim reinfection and vaccination and excluded any sample collected within 45 days of either event. People with LC were recruited at three sites, each applying a case definition of at least one new symptom persisting at least 3 months after SARS-CoV-2 infection. Each site independently recruited recovered controls (**Tables 1-2, Tables S1-S3**, further details in Methods). Participants were screened for expression of HLA-A*02:01 and HLA-A*03:01, identifying 475 unique samples from 288 individuals that were suitable for analysis using HLA-I/peptide tetramers to study CD8 T cells specific for SARS-CoV-2, CMV, influenza A (Flu), VZV, and lytic and latent EBV antigens (**Fig. 1B, Table S4**). A subset of individuals recruited at the University of Pennsylvania were found to express HLA-DPB1*04:01 and/or HLA-DRB1*15:01, and 77 samples from these individuals were analyzed for SARS-CoV-2 spike-specific CD4 T cell responses (**Fig. 1B**). To capture differences in *in vivo* activation of virus-specific T cells, we measured several markers associated with T cell activation (CD38, CD39, PD-1, and Ki67 for both CD4 and CD8 T cells, plus HLA-DR for CD8 T cells and CTLA-4 and ICOS for CD4 T cells), allowing quantification of activated T cells expressing at least 2 activation markers directly *ex vivo* (**Fig. 1B, Fig. S1-S2**).

**Fig. 1:**
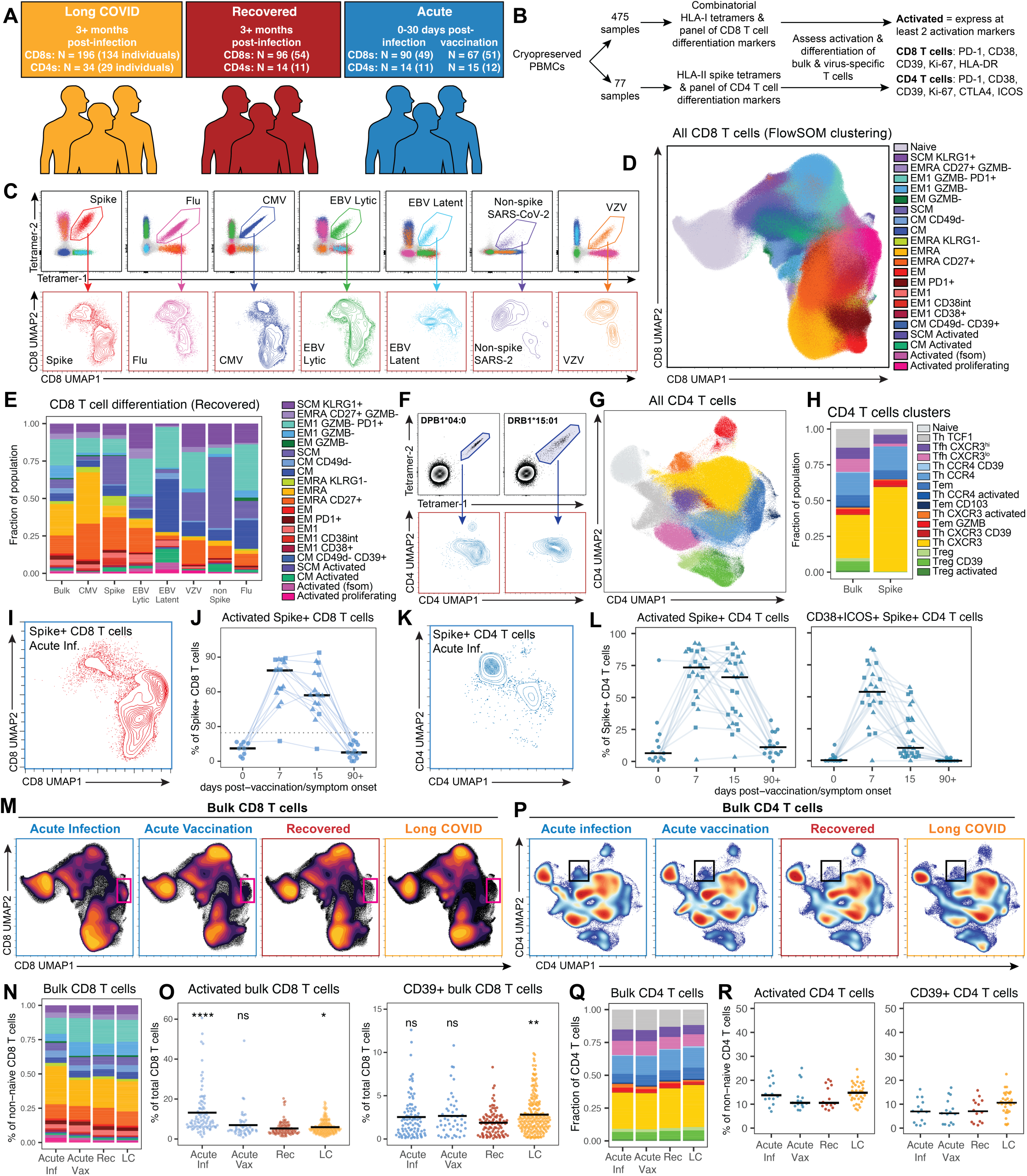
Spectral flow cytometry identifies distinct differentiation states for virus-specific T cells and minimal changes to the bulk T cell landscape in long COVID. (**A**) Cohort overview, see Tables 1-2 and Tables S1-3 for details. (**B**) Summary of flow cytometric workflow and strategy used to identify activated T cells, see Fig. S1-2. (**C**) Representative gating of HLA-I tetramer+ CD8 T cell populations from n=20 concatenated samples (top) and UMAP coordinates of HLA-I tetramer+ populations in recovered controls (n=96, bottom). (**D**) UMAP projection of FlowSOM metaclusters identifying differentiation states of CD8 T cells. (**E**) Summary data of the percent of the indicated tetramer+ CD8 T cell populations found in each FlowSOM metacluster in recovered controls. (**F**) Representative plots of HLA-II tetramer+ CD4 T cells (top) and UMAP coordinates of HLA-II tetramer+ populations in recovered controls (bottom). (**G**) UMAP projection of FlowSOM metaclusters identifying differentiation states of CD4 T cells. (**H**) Summary data of the percent of bulk or spike-specific CD4 T cells found in each FlowSOM metacluster in recovered controls. (**I**) UMAP coordinates of spike-specific CD8 T cells during acute infection. (**J**) Activation (as defined in B) of spike-specific CD8 T cells from longitudinal samples collected before, during, and after acute infection or vaccination. Squares represent acute infection and triangles represent acute vaccination samples. Lines connect individuals sampled longitudinally. (**K**) UMAP coordinates of spike-specific CD4 T cells during acute infection. (**L**) Activation (as defined in B) and percent CD38+ICOS+ of spike-specific CD4 T cells from longitudinal samples collected before, during, and after acute infection or vaccination. Squares represent acute infection and triangles represent acute vaccination samples. Lines connect individuals sampled longitudinally. (**M**) UMAP projection of bulk CD8 T cells concatenated from all samples in the indicated categories. Pink box highlights activated regions in CD8 T cell UMAP space. (**N**) Summary data of the percent of bulk non-naive CD8 T cells found in each FlowSOM metacluster as in (E). (**O**) Summary data of the activation (as defined in B) and percent CD39+ of bulk CD8 T cells from all samples in the indicated categories. (**P**) UMAP projection of bulk CD4 T cells concatenated from all samples in the indicated categories. Black box highlights activated regions in CD4 T cell UMAP space. (**Q**) Summary data of the percent of bulk non-naive CD4 T cells found in each FlowSOM metacluster as in (H). (**R**) Summary data of the activation (as defined in B) and percent CD39+ of bulk CD4 T cells from all samples in the indicated categories. All statistical tests were Wilcoxon tests with Benjamini-Hochberg correction for multiple comparisons. *p < 0.05, **p < 0.01, ***p < 0.001, ****p < 0.0001. In the absence of brackets, all statistical tests are comparisons to recovered controls.

**Table 1:** All samples used for CD8 T cell analyses in Fig. 1-3.

| Characteristic | Study Cohorts |  |  |  |
| --- | --- | --- | --- | --- |
|  | LC<br>N = 196<sup>1</sup> | Recovered<br>N = 96<sup>1</sup> | Acute infection<br>N = 90<sup>1</sup> | Acute vaccination<br>N = 67<sup>1</sup> |
| **age (years)** | 47.6 (22 - 80) | 37.9 (22 - 85) | 47.3 (22 - 75) | 40.1 (22 - 86) |
| **sex** |  |  |  |  |
| Female | 134 (68%) | 46 (48%) | 38 (42%) | 42 (63%) |
| Male | 62 (32%) | 50 (52%) | 52 (58%) | 25 (37%) |
| **ethnicity** |  |  |  |  |
| Hispanic/Latino | 24 (12%) | 7 (7.3%) | 2 (2.2%) | 9 (13%) |
| Not Hispanic/Latino | 172 (88%) | 89 (93%) | 88 (98%) | 58 (87%) |
| **race** |  |  |  |  |
| Asian | 7 (4.0%) | 5 (5.5%) | 3 (3.3%) | 5 (8.3%) |
| Black | 6 (3.5%) | 3 (3.3%) | 31 (34%) | 0 (0%) |
| Native American | 2 (1.2%) | 1 (1.1%) | 0 (0%) | 1 (1.7%) |
| White | 158 (91%) | 82 (90%) | 56 (62%) | 54 (90%) |
| Unknown/Other | 23 | 5 | 0 | 7 |
| **first infection date** |  |  |  |  |
| March 2020 - April 2021 | 153 (79%) | 45 (47%) | 47 (52%) | NA |
| May 2021 - November 2021 | 14 (7.3%) | 8 (8.3%) | 6 (6.7%) | NA |
| >=December 2021 | 26 (13%) | 43 (45%) | 37 (41%) | NA |
| Unknown | 3 | 0 | 0 |  |
| **n vaccine doses** | 1.6 (0 - 6) | 2.7 (0 - 5) | 1.7 (0 - 4) | 2.7 (1 - 5) |
| Unknown | 3 | 0 | 4 | 0 |
| **n SARS-CoV-2 infections** | 1.1 (1 - 3) | 1.3 (1 - 3) | 1.2 (1 - 3) | 0.8 (0 - 2) |
| Unknown | 3 | 0 | 0 | 0 |
| **days since last vaccine/infection** | 182.6 (46 - 627) | 190.8 (45 - 641) | 8.3 (0 - 23) | 13.7 (2 - 29) |
| **hospitalized with COVID-19** | 29 (15%) | 1 (1.0%) | 45 (51%) | NA |
| **days since symptom onset** | 356.8 (92 - 1,196) | 331.2 (88 - 1,021) | 8.3 (0 - 23) | NA |
| Unknown | 3 | 0 | 0 |  |

<sup>1</sup>Mean (Min - Max); n (%)
N = number of samples

**Table 2:** All samples used for CD4 T cell analyses in Fig. 1-2.

| **Characteristic** | **LC**<br>**N = 34<sup>1</sup>** | **Recovered**<br>**N = 14<sup>1</sup>** | **Acute infection**<br>**N = 14<sup>1</sup>** | **Acute vaccination**<br>**N = 15<sup>1</sup>** |
| --- | --- | --- | --- | --- |
| **age (years)** | 50.4 (20 - 80) | 31.9 (25 - 50) | 31.9 (25 - 50) | 31.8 (22 - 52) |
| **sex** |  |  |  |  |
| Female | 25 (74%) | 9 (64%) | 9 (64%) | 10 (67%) |
| Male | 9 (26%) | 5 (36%) | 5 (36%) | 5 (33%) |
| **ethnicity** |  |  |  |  |
| Hispanic/Latino | 2 (5.9%) | 1 (7.1%) | 1 (7.1%) | 1 (6.7%) |
| Not Hispanic/Latino | 31 (91%) | 13 (93%) | 13 (93%) | 14 (93%) |
| Unknown | 1 (2.9%) | 0 (0%) | 0 (0%) | 0 (0%) |
| **race** |  |  |  |  |
| Black | 1 (3.1%) | 1 (7.1%) | 1 (7.1%) | 2 (13%) |
| White | 31 (97%) | 13 (93%) | 13 (93%) | 13 (87%) |
| Unknown | 2 | 0 | 0 | 0 |
| **first infection date** |  |  |  |  |
| March 2020 - April 2021 | 9 (30%) | 0 (0%) | 0 (0%) | NA |
| May 2021 - November 2021 | 0 (0%) | 0 (0%) | 0 (0%) | NA |
| >=December 2021 | 21 (70%) | 14 (100%) | 14 (100%) | NA |
| Unknown | 4 | 0 | 0 |  |
| **n vaccine doses** | 3.3 (0 - 5) | 3.7 (3 - 5) | 3.8 (3 - 4) | 4.3 (4 - 5) |
| Unknown | 4 | 0 | 0 | 0 |
| **n SARS-CoV-2 infections** | 1.4 (1 - 3) | 1.3 (1 - 2) | 1.5 (1 - 3) | 0.9 (0 - 2) |
| Unknown | 4 | 0 | 0 | 0 |
| **days since last vaccine/infection** | 254.1 (79 - 627) | 145.4 (103 - 193) | 9.9 (5 - 16) | 8.3 (6 - 21) |
| **hospitalized with COVID-19** |  |  |  |  |
| No | 33 (100%) | 14 (100%) | 14 (100%) | NA |
| Unknown | 1 | 0 | 0 |  |
| **days since symptom onset** | 528.5 (213 - 1,126) | 262.3 (106 - 708) | 9.9 (5 - 16) | NA |
| Unknown | 4 | 0 | 0 |  |

<sup>1</sup>Mean (Min - Max); n (%)
N = number of samples

We used multiplexed combinatorial HLA-I/peptide tetramer staining to simultaneously detect CD8 T cells specific for SARS-CoV-2 spike and non-spike proteins, Flu, CMV, EBV lytic and latent antigens, and VZV (**Fig. 1C** and **Table S4**)(*45, 51*). For participants expressing HLA-A*02:01, a well-characterized herpesvirus cross-reactive (HVCR) response was also identified(*55*). CD8 T cells targeting the non-spike epitopes from SARS-CoV-2 used in this study were rare in most samples, preventing robust analysis of these responses (**Fig. S3A-B**). Dimensionality reduction and clustering of circulating CD8 T cells based on expression of 17 markers identified naive cells, multiple clusters of granzyme B (GZMB)+ effector memory cells and GZMB-effector and central memory cells, and several clusters of activated cells (**Fig. 1D, Fig. S4A-B**). CD8 T cells from recovered controls occupied divergent differentiation states depending on their viral specificity, with CMV-specific cells enriched for GZMB+ terminal effector states and Flu-specific cells enriched for GZMB-central memory cells, whereas SARS-CoV-2 spike-specific CD8 T cells were divided between these states (**Fig. 1D-E**). A similar approach using two previously described immunodominant epitopes restricted to HLA-DPB1*04:01 and HLA-DRB1*15:01 identified spike-specific CD4 T cells (**Fig. 1F**)(*53, 56–58*). Several clusters associated with known CD4 T cell differentiation states were identified (**Fig. 1G**, **Fig. S5A-B**), and spike-specific CD4 T cells from recovered individuals predominantly displayed a TH1-like CXCR3+ memory phenotype (**Fig. 1F-H**).

To first test whether these assays could detect *in vivo* T cell biosensor activity during a known antigen challenge, we studied longitudinal sample sets collected before, during and after SARS-CoV-2 mRNA booster vaccination or breakthrough infection. As we previously reported(*51*), memory spike-specific CD8 T cells had low baseline activation before infection or vaccination. However, these cells were robustly activated by day 7 post-vaccination or post-symptom onset, remained activated at day 15, and returned to baseline activation status beyond day 90 (**Fig. 1I-J**). Robust activation of spike-specific CD4 T cells was also detected after vaccination or infection, followed similar kinetics, and likewise returned to baseline by 90 days post-exposure (**Fig. 1K-L**). Thus, flow cytometric assays incorporating multiplexed HLA tetramers are sufficiently sensitive to detect transient *in vivo* activation of virus-specific CD4 and CD8 T cells upon antigen encounter.

### The overall T cell landscape is not substantially altered in long COVID

We next sought to use these assays to address whether the activation and differentiation of bulk CD8 and CD4 T cells were altered in LC independent of antigen specificity. The overall landscape of circulating CD8 T cell populations was largely indistinguishable comparing acute infection, vaccination, fully recovered, and LC samples, with only minor enrichment in activated clusters during acute infection (**Fig. 1M-N**). LC was not associated with significant changes in the proportion of any major CD8 T cell population in circulation (**Fig. 1N**). Bulk T cell activation was increased during acute SARS-CoV-2 infection and very modestly but significantly increased in LC compared to recovered controls, primarily driven by a small but significant increase in CD39 expression (**Fig. 1O**). A similar pattern was observed for bulk CD4 T cells, with minimal changes to the overall T cell landscape (**Fig. 1P-Q, Fig. S5C**) and a trend towards mildly elevated activation in acute SARS-CoV-2 infection and LC, with activation in LC driven by a small increase in CD39 expression (**Fig. 1R**). Thus, in agreement with several prior studies(*12, 48, 54, 59–62*), we did not observe broad alterations in bulk circulating T cell populations in people with LC. These observations suggest a lack of bystander immune activation in LC and are consistent with the notion that, even if viral antigens were present in LC, only a small fraction of circulating T cells would be specific for those particular antigens.

### A subset of people with LC have persistently elevated activation of spike-specific CD8 T cells

We therefore analyzed SARS-CoV-2-specific CD8 T cells for evidence of recent stimulation by SARS-CoV-2 spike protein in people with LC (**Fig. 2A**). Activation of spike-specific CD8 T cells was generally low in individuals who fully recovered after SARS-CoV-2 infection compared to samples taken during acute infection or vaccination, consistent with antigen clearance in most recovered individuals (**Fig. 2A-B, Fig. S6A**). In contrast, people with LC had markedly increased activation of spike-specific CD8 T cells directly *ex vivo* compared to controls, and this remained statistically significant after adjusting for age, sex, race, and SARS-CoV-2 exposure history (**Fig. 2A-B, Fig. S6A-B**). Notably, spike-specific CD8 T cell activation was driven by a subset of people with LC, and this pattern was observed across all three sites where samples were collected (**Fig. S6C**). Using a cutoff of one standard deviation above the mean activation observed in recovered controls, 40% of LC samples had elevated activation of spike-specific CD8 T cells (termed Spike^Act^LC subset hereafter), compared to only 16% of recovered controls (**Fig. 2B**, OR = 3.63, 95% CI = 1.90-7.30, p = 0.000016, Fisher’s exact test). The Spike^Act^ LC subset had significant increases in the proportion of spike-specific cells found in several activated/effector clusters, including clusters defined as activated proliferating, activated, activated CM, CD39+ CM, CD38+ EM1, and EM1, and significant decreases in resting memory clusters, including CM, EM1 GZMB-, and SCM KLRG1+ (**Fig. 2C**). SPICE analysis revealed that the activation signal in Spike^Act^ LC was driven by multiple activation markers, including CD39, CD38, HLA-DR, and to a lesser extent Ki67 (**Fig. 2D, Fig. S6D**). CD8 T cells targeting non-spike epitopes from SARS-CoV-2 were near the limit of detection in most samples (**Fig. S3A-B**), but there was no corresponding increase in activation of these cells in LC (**Fig. S6E**). CD8 T cells specific for another respiratory virus, influenza A (Flu), did not show elevated activation in people with LC (**Fig. 2E**, OR = 1.71, 95% CI = 0.78-4.1, p = 0.17, Fisher’s exact test). Low-level bystander activation of Flu-specific CD8 T cells was observed during acute SARS-CoV-2 infection but not in LC, suggesting that activation of spike-specific CD8 T cells in LC was not due to bystander activation (**Fig. 2E**).

**Fig. 2:**
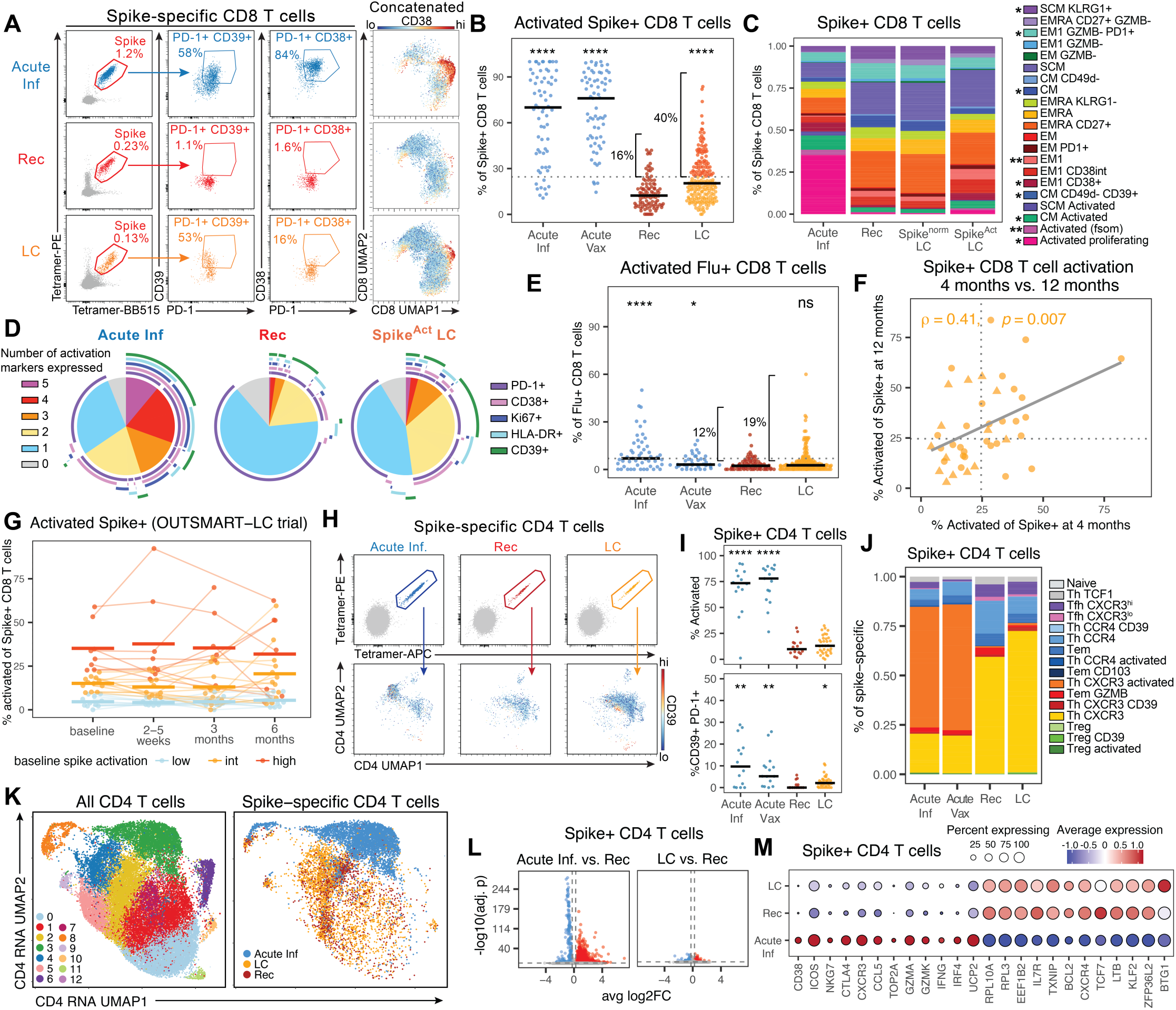
A subset of people with LC have persistently elevated activation of spike-specific CD8 T cells. (**A**) Representative flow cytometry plots from individual samples displaying the gating of spike-specific CD8 T cells and the expression of activation markers by these cells (left), and the positions in UMAP space of all spike-specific CD8 T cells, concatenated by category and colored by the intensity of CD38 expression (right). (**B**) Summary data of the activation (defined in Fig. 1B and Fig. S1) of spike-specific CD8 T cells by group. Horizontal dotted line represents the mean plus one standard deviation of recovered controls, and the percent of samples falling above this threshold is indicated. (**C**) Summary data of the percent of the indicated spike-specific CD8 T cell populations found in each FlowSOM metacluster. LC samples with activation above or below the threshold in B are referred to as Spike^Act^ LC and Spike^norm^ LC, respectively. Asterisks in the legend indicate statistically significant differences in cluster frequency comparing Spike^Act^ LC to recovered controls. (**D**) SPICE analysis of the co-expression of five activation markers by spike-specific CD8 T cells from the indicated groups. (**E**) Summary data of the activation of Flu-specific CD8 T cells by group. Horizontal dotted line represents the mean plus one standard deviation of recovered controls, and the percent of samples falling above this threshold is indicated. (**F**) Correlation (calculated using nonparametric Spearman rank correlation) of the percent activation of spike-specific CD8 T cells from paired longitudinal LC samples from the LIINC cohort collected at 4 and 12 months post-infection. (**G**) Longitudinal analysis of spike-specific CD8 T cell activation of samples from the OUTSMART-LC clinical trial, displaying the mean activation from all samples for each individual at the indicated timepoints. Samples were divided into low (x < 9.5%), intermediate (9.5% < x < 22%), and high (x > 22%) average activation at baseline timepoints. Lines connect individuals sampled longitudinally. (**H**) Representative plots of HLA class II tetramer staining of CD4 T cells (top) and the positions in UMAP space of all spike-specific CD4 T cells, concatenated by category and colored by the intensity of CD39 expression (bottom). (**I**) Dot plots displaying the activation status (defined in Fig. 1B and Fig. S2) as well as the coexpression of CD39 and PD-1 on spike-specific CD4 T cells. (**J**) Summary data of the percent of the indicated spike-specific CD4 T cell populations found in each FlowSOM metacluster. (**K**) UMAP projections of single-cell RNA sequencing data for all bulk and spike-specific CD4 T cells colored by cluster (left) and for spike-specific CD4 T cells only colored by group (right). (**L**) Volcano plots displaying upregulated (red) and downregulated (blue) DEGs in spike-specific CD4 T cells from acute infection (left) or LC (right) compared to recovered controls. Spike-specific CD4 T cells from each sample were downsampled to a maximum of 250 cells to avoid bias associated with spike-specific CD4 T cell recovery. Significant DEGs were identified via Seurat’s FindMarkers function by applying the model-based analysis of single-cell transcriptomics (MAST) framework and considering sex as a latent variable. (**M**) Dot plot displaying the expression of several genes associated with activated or resting/memory T cells in spike-specific CD4 T cells, using downsampled populations as in (L). Statistical significance was determined using Wilcoxon tests with Benjamini-Hochberg correction for multiple comparisons. *p < 0.05, **p < 0.01, ***p < 0.001, ****p < 0.0001. In the absence of brackets, all statistical tests are comparisons to recovered controls.

LC is a chronic condition, and in some cases symptoms persist for years. To test whether elevated activation of spike-specific CD8 T cells was maintained over time, we leveraged longitudinal sampling from the LIINC cohort to measure activation at 4 months and 12 months post-acute infection, excluding participants with any interim SARS-CoV-2 infection or vaccination. Activation of spike-specific CD8 T cells was significantly correlated at 4 and 12 months, with individuals showing either consistently low or consistently elevated activation (**Fig. 2F**). To extend this analysis to later timepoints and more frequent sampling, we analyzed longitudinal samples from the placebo-controlled OUTSMART-LC clinical trial (NCT05877508), which tested a monoclonal antibody targeting SARS-CoV-2 spike (AER002) in people with LC(*63*). Although AER002 did not improve LC symptoms relative to placebo, the trial provided densely sampled specimens from people with established LC. In this cohort, elevated activation of spike-specific CD8 T cells was again present in a subset of participants and remained stable across follow-up timepoints over the subsequent 6 months (**Fig. 2G, Fig. S7**). Participants stratified by high, intermediate, or low baseline activation of spike-specific CD8 T cells maintained their activation status over time (**Fig. 2G, Fig. S7A-B**). Baseline activation of spike-specific CD8 T cells was strongly correlated with activation 2-5 weeks post-treatment and remained so at 3 and 6 months (**Fig. S7C**). Participants with low activation at baseline maintained low activation throughout 6 months of sampling. Taken together, these data suggest that a subset of people with LC can be defined by maintaining elevated activation of spike-specific CD8 T cells for months after the initial infection, consistent with a possible persistent reservoir of SARS-CoV-2.

### Activation of spike-specific CD4 T cells is less prominent than CD8 T cells in LC

We next asked whether spike-specific CD4 T cells were similarly activated in LC. Spike-specific CD4 T cells were robustly activated during acute SARS-CoV-2 infection, with a return to low activation in individuals who fully recovered (**Fig. 2H-I, Fig. S8A-B**). In the smaller cohort used for CD4 T cell analyses, there was a modest trend towards elevated activation of spike-specific CD4 T cells in people with LC driven by a small but statistically significant increase in CD39 expression (**Fig. 2H-I**). Spike-specific CD4 T cells from people with LC were found in similar differentiation states to those from recovered controls (**Fig. 2J, Fig. S8C**). In a subset of samples where HLA types allowed both CD4 and CD8 T cells to be analyzed, a modest but statistically significant correlation between CD39 expression on spike-specific CD4 T cells and activation of spike-specific CD8 T cells was observed (**Fig. S8D**). Expression of CD39 on CD4 T cells did not reach levels higher than 8%, however, even in samples where over 30% of spike-specific CD8 T cells were activated. Thus, in contrast to CD8 T cells, there was only minimal evidence that spike-specific CD4 T cells were activated in a subset of people with LC.

To investigate whether spike-specific CD4 T cells may have transcriptional signals of activation that could not be captured by flow cytometry, we performed 10x single-cell sequencing on sorted spike-specific and bulk CD4 T cells from 8 people with LC, 4 recovered controls, and 4 acute SARS-CoV-2 infection samples (**Fig. S8E-F**). In agreement with the flow cytometry data for these samples, spike-specific CD4 T cells from LC clustered with those from recovered controls, whereas spike-specific CD4 T cells from acute infection occupied distinct clusters (**Fig. 2K, Fig. S9A-D**). DEG analysis revealed robust changes in gene expression for spike-specific CD4 T cells during acute infection compared to recovered control samples, but very few DEGs for LC compared to recovered controls (**Fig. 2L**). The few DEGs identified in LC overlapped partially with DEGs from acute samples (**Fig. S9E**). Nevertheless, spike-specific CD4 T cells from LC more closely resembled cells from recovered controls across a large suite of genes spanning activation, proliferation, translation, and memory biology (**Fig. 2M, Fig. S9F**). Thus, activation of spike-specific CD4 T cells was less prominent compared to the robust activation of spike-specific CD8 T cells in a subset of people with LC.

### EBV-specific CD8 T cells are persistently activated in a subset of people with LC

Reactivation of common herpesviruses, especially EBV, can occur during acute SARS-CoV-2 infection and is associated with the development of LC(*9–12, 36*). It is unclear, however, whether EBV returns to a latent state in people with LC or continues to express lytic antigens. CD8 T cells targeting lytic antigens from EBV had significantly increased activation in samples from both acute SARS-CoV-2 infection and LC compared to recovered controls (**Fig. 3A-B**, OR = 3.04, 95% CI = 1.4-7.2, p = 0.0026, Fisher’s exact test for recovered vs. LC). This observation is consistent with EBV reactivation during acute infection and sustained lytic antigen expression in a subset of people with LC. Activation of CD8 T cells recognizing the herpesvirus cross-reactive (HVCR) epitope, which is also a lytic phase EBV antigen, followed a similar pattern (**Fig. S10A**) and was strongly correlated with the activation of CD8 T cells recognizing EBV lytic epitopes in the same samples (**Fig. S10B**). EBV latent-, CMV- and VZV-specific CD8 T cells did not have significantly elevated activation in LC, although these populations had slightly elevated activation during acute infection (**Fig. 3C-D**, **Fig. S10C**), Fisher’s exact test: not significant). Similar to what was previously observed for spike, activation of EBV lytic-specific CD8 T cells was strongly correlated at 4 and 12 months post-acute infection in people with LC (**Fig. S10D**). Thus, elevated activation of EBV lytic antigen-specific CD8 T cells, but not CMV, VZV, or EBV latent antigen-specific cells, was sustained in a subset of people with LC.

**Fig. 3:**
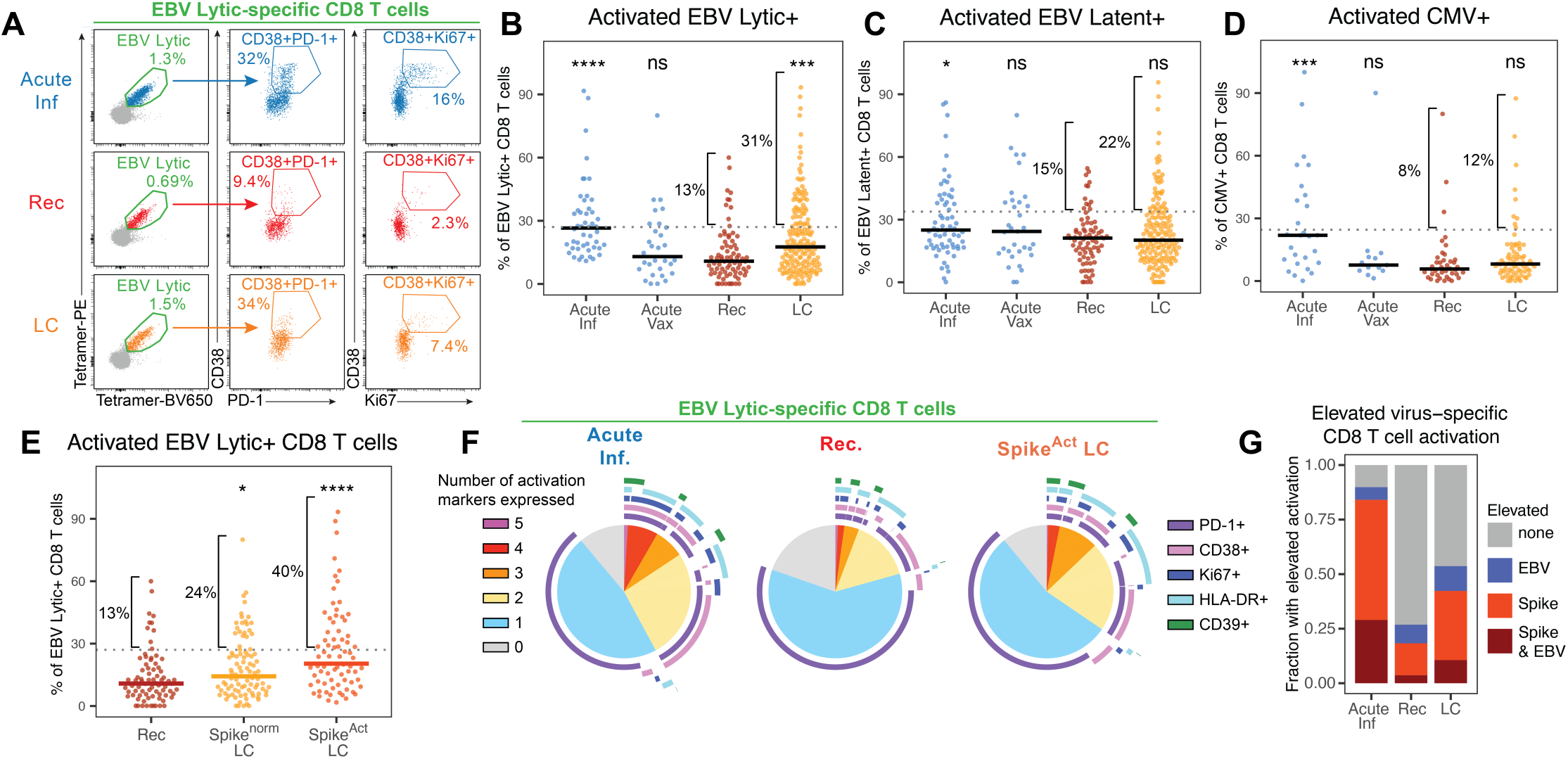
EBV-specific CD8 T cells are persistently activated in a subset of people with LC. (**A**) Representative flow cytometry plots from individual samples displaying the gating of EBV lytic antigen-specific CD8 T cells and the expression of activation markers by these cells. (B-D) Summary data of the activation (defined in Fig. 1B and Fig. S1) of (**B**) EBV lytic-specific, (**C**) EBV latent-specific, and (**D**) CMV-specific CD8 T cells by group. Horizontal dotted lines represent the mean plus one standard deviation of recovered controls, and the percent of samples falling above this threshold is indicated. (**E**) Summary data of the activation of EBV lytic-specific CD8 T cells with the LC group divided based on spike-specific CD8 T cell activation as in Fig. 2B. (**F**) SPICE analysis of the co-expression of five activation markers by EBV lytic-specific CD8 T cells from the indicated groups. (**G**) Summary data indicating the percent of samples in each group where elevated activation of spike- and/or EBV lytic-specific CD8 T cells was observed. Elevated activation was determined using the mean plus one standard deviation observed in recovered controls indicated by dotted lines in Fig. 2B and Fig. 3B. Statistical significance was determined using Wilcoxon tests with Benjamini-Hochberg correction for multiple comparisons. *p < 0.05, **p < 0.01, ***p < 0.001, ****p < 0.0001. In the absence of brackets, all statistical tests are comparisons to recovered controls.

We next asked whether people with LC who had elevated activation of spike- and EBV-specific CD8 T cells represented distinct biological endotypes of LC. Elevated EBV lytic-specific CD8 T cell activation was largely restricted to the Spike^Act^ LC subset (**Fig. 3E**), and activation of EBV lytic-specific CD8 T cells was correlated with activation of spike-specific CD8 T cells in LC (**Fig. S10E**). Activation of EBV lytic-specific CD8 T cells in the Spike^Act^ LC subset involved co-expression of multiple activation markers, including CD38, HLA-DR, and Ki67 (**Fig. 3F**). CMV-specific CD8 T cells had elevated expression of HLA-DR only, and Flu-, EBV latent-, and VZV-specific CD8 T cells showed no signs of increased activation in LC (**Fig. S11**). Altogether, over half of people with LC had elevated activation of spike- and/or EBV lytic-specific CD8 T cells (**Fig. 3G**). These data collectively indicate that LC was not associated with broad bystander activation, but rather was uniquely linked to activation of spike-specific, and to a lesser extent EBV lytic-specific, CD8 T cells.

### Single-cell RNA sequencing with barcoded HLA-I/peptide tetramers identifies shared gene expression profiles for spike-specific CD8 T cells from LC and acute SARS-CoV-2 infection

To more fully investigate the molecular basis of spike- and EBV-specific CD8 T cell activation in LC, we selected samples from recovered controls, acute SARS-CoV-2 infection, and people with LC who had elevated activation of spike- and EBV-specific CD8 T cells for single-cell RNA, TCR, and antigen specificity barcode sequencing (**Fig. 4A**). Samples were stained with fluorophore and oligonucleotide-barcoded HLA-I tetramers loaded with 27 distinct viral peptides using the 10x Genomics Barcode Enabled Antigen Mapping (BEAM) platform, with a unique combination of two oligonucleotide barcodes used for each specificity (**Fig. S12A-B, Table S4**). Each sample was labeled with a unique hashtag oligonucleotide (**Fig. S12C**), and antigen-specific and bulk CD8 T cells were sorted and pooled for sequencing. Dimensionality reduction and clustering identified 18 clusters corresponding to known circulating CD8 T cell states (**Fig. 4B-C, Fig. S13**). Clusters 2 and 18 represent naive cells with and without an interferon stimulated gene signature. Clusters 1, 3, 9, 13, 15, and 16 have GZMB+ effector features (GZMB, CX3CR1, FGFBP2) and lack memory features (IL7R, SELL, TCF7), whereas clusters 0, 4, 10, 12, and 17 have memory features and lack effector features. Of these, clusters 0 and 11 (which lacks memory features), are GZMK high, whereas the others have intermediate expression of GZMK. Cluster 7 represents activated proliferating cells, cluster 6 contains less proliferative activated cells, and cluster 8 has features of activation and T cell exhaustion. Cluster 5 is high for IKZF2 and TYROBP, clusters 9 (effector) and 17 (naive/memory) have IFN signatures, and clusters 12-18 are minor populations. Cluster 14 is a participant-specific cluster that was excluded from further analysis (**Fig. S13**). Overall, these populations reflected those observed in the flow cytometric analysis above (**Fig. 1D**) but with greater depth and improved resolution of distinct memory T cell states.

**Fig. 4:**
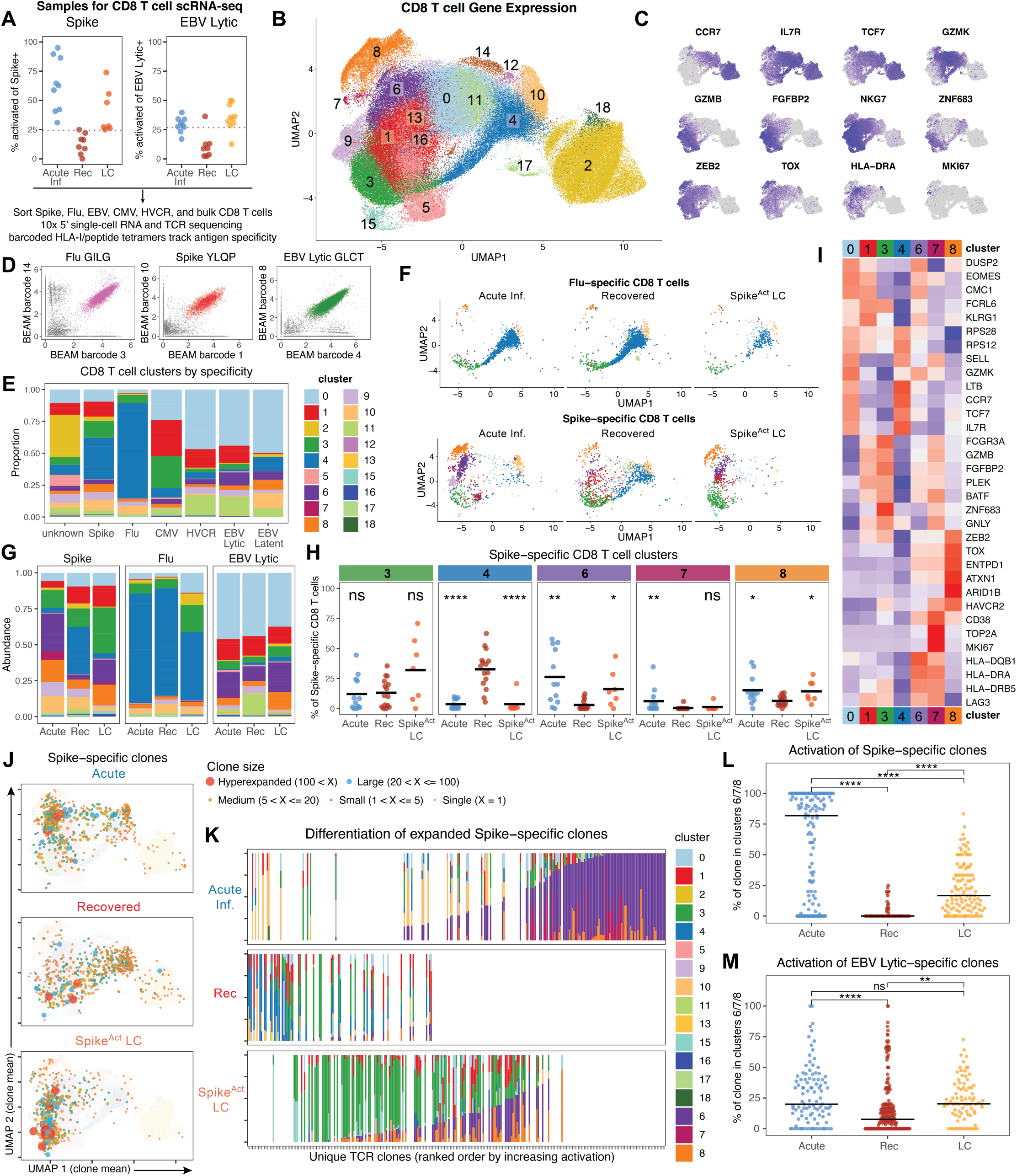
Single-cell sequencing with barcoded HLA-I/peptide tetramers identifies shared gene expression profiles for Spike-specific CD8 T cell clones from LC and acute SARS-CoV-2 infection. (**A**) Summary flow cytometric data describing the activation of spike- and EBV lytic-specific CD8 T cells in the samples selected for single-cell RNA, TCR, and tetramer barcode sequencing. (**B**) UMAP representation of single-cell RNA sequencing data of all bulk and antigen-specific CD8 T cells, colored by cluster. (**C**) Feature plots displaying the expression of selected genes by CD8 T cells in UMAP space. (**D**) Representative plots showing examples of tetramer barcode sequencing reads used to identify antigen-specific CD8 T cells in single-cell sequencing data. (**E**) Summary data of the percent of the indicated tetramer+ CD8 T cell populations found in each scRNA-seq cluster in recovered control samples. (**F**) UMAP coordinates of Flu-(top) and spike-specific (bottom) CD8 T cells isolated from each group. Antigen-specific populations were downsampled to a maximum of 150 cells from each sample to avoid biasing these representations towards high magnitude responses. (**G**) Summary data of the percent of the indicated tetramer+ CD8 T cell populations found in each scRNA-seq cluster, split by group. (**H**) Quantification of the frequency of spike-specific CD8 T cells found in a selection of the most prominent and variable clusters for each sample, split by group. (**I**) Heatmap displaying average expression of selected genes across all CD8 T cells found in the clusters analyzed in (H). (**J**) Mean UMAP coordinates for all cells within each spike-specific CD8 T cell clonotype. Clones were defined using the CTaa parameter from the scRepertoire package, and the size and color of the points represent the number of cells recovered within each clonotype. (**K**) Ranked ordering by percent of cells falling in activated clusters 6, 7, or 8 for all spike-specific CD8 T cell clones with at least 5 cells. Each column represents a unique clone, ranked along the x-axis and colored by the proportion of cells assigned to each cluster. (**L-M**) Summary data for the percent of each spike-(L) and EBV lytic-specific (**M**) CD8 T cell clone in activated clusters 6, 7, or 8. Each point represents a unique clone, and only clones with at least 5 cells were included. Statistical significance was determined using Wilcoxon tests with Benjamini-Hochberg correction for multiple comparisons. *p < 0.05, **p < 0.01, ***p < 0.001, ****p < 0.0001. In the absence of brackets, all statistical tests are comparisons to recovered controls.

Sequencing of BEAM barcodes allowed highly sensitive and specific detection of antigen-specific CD8 T cells using a dual-barcode approach, including over 16,000 spike-specific, 20,000 EBV-specific, 9,000 CMV-specific, and 9,000 Flu-specific CD8 T cells identified together with over 115,000 bulk CD8 T cells (**Fig. 4D, Fig. S12B, Fig. S14A**). As observed in the cytometric analysis, mapping of antigen-specific populations to gene expression clusters in recovered controls revealed distinct homeostatic differentiation states of memory T cells associated with different antigen specificities (**Fig. 4E, Fig. S14B**). CD8 T cells specific for Flu antigens, and to a lesser extent spike, were mostly found in memory cluster 4, whereas CMV-specific CD8 T cells were predominantly found in GZMB+ effector clusters 1 and 3 and EBV-specific CD8 T cells were dominated by GZMK+ clusters 0 and 11. Bulk cells with unknown antigen specificity had contributions from most clusters and were enriched for naive cells compared to known antigen-specific populations (**Fig. 4E**).

We then asked whether LC was associated with changes in gene expression compared to recovered controls for any of the virus-specific CD8 T cell populations. In contrast to Flu-, CMV-, and EBV latent-specific CD8 T cells, which occupied a similar gene expression space across acute infection, recovered controls, and LC samples, spike-specific CD8 T cells were much more dynamic (**Fig. 4F-G, Fig. S14C**). Exposure to SARS-CoV-2 spike antigens during acute infection drove spike-specific CD8 T cells into distinct regions of gene expression space that were not commonly occupied in recovered controls, with a disappearance of cells in memory-associated cluster 4 and the emergence of cells in activated clusters 6, 7, and 8 (**Fig. 4F-H**). Spike-specific CD8 T cells from LC were similarly absent from cluster 4 and instead enriched in activation-associated clusters 6 and 8 and in effector-like cluster 3 in some individuals (**Fig. 4F-H**). CD8 T cells specific for EBV lytic antigens showed a similar but more modest effect, with a significantly higher frequency of cells in clusters 6 and 8 in LC compared to recovered controls (**Fig. S14C-D**).

To better understand the biology underlying these clusters, we explored the differentially expressed genes that defined each of these clusters (**Fig. 4I**) and performed gene set enrichment analysis (GSEA) for gene expression profiles (GEPs) from T-CellAnnoTator (TCAT)(*64*) (**Fig. S15A**). Cluster 3, which was overrepresented in some people with LC, likely represents terminal effector cells, with high expression of FCGR3A, GZMB, FGFBP2, GNLY, and ZNF683 (and enriched for genes associated with cytotoxic and cytoskeleton GEPs) and low expression of SELL, TCF7, and CCR7 (and depleted for genes associated with naive/resting and protein translation GEPs) (**Fig. 4I, Fig. S15A**). Cells in cluster 3 may also be moderately activated, as these cells expressed HAVCR2, CD38, LAG3, and HLA-DRB5 and had enrichment for the activation-associated TIMD4/TIM3 and HLA GEPs. Cluster 4, which was common in spike-specific CD8 T cells in recovered controls but depleted in acute and LC samples, enriched for memory and lacked cytotoxic and activation-associated genes. Cluster 6, which was prevalent in spike-specific cells from acute infection and LC, is a highly activated cluster, expressing numerous HLA-II genes, LAG3, CD38, and ENTPD1 (and enriched for genes in activation-associated GEPs including HLA, ICOS/CD38, TIMD4/TIM3, CTLA4/CD38, and various cell cycle GEPs). Cluster 7, increased only in acute infection, is an activated and proliferative cluster with high expression of TOP2A and MKI67.

Cluster 8 expressed a suite of genes associated with T cell exhaustion, including TOX, ENTPD1, ARID1B, HAVCR2, CD38, and ZEB2. Interestingly, TCAT analysis revealed that cluster 8 was associated with the Poor-Quality GEP (**Fig. S15A**). Despite passing initial filtering and QC cutoffs (see Methods), cluster 8 had relatively low RNA counts and a higher percent of mitochondrial reads compared to other clusters (**Fig. S15B**). However, cluster 8 was significantly enriched in spike-specific CD8 T cells during acute infection, where ongoing activation can produce a large population of short-lived effector cells that are predisposed to die over the ensuing days. Across all specificities and all individuals, the frequency of CD8 T cells in cluster 8 was strongly correlated with the frequency of cells in the other activated clusters 6 and 7 (**Fig. S15C**). Thus, although these cells are often excluded in conventional scRNA-seq workflows, the additional resolution provided by barcoded tetramers made it possible to capture their contribution to the activated CD8 T cell response. Nevertheless, cells in cluster 8 were excluded from later DEG and GSEA analyses due to their low RNA count. Taken together, these analyses indicate that spike-specific CD8 T cells from people with LC occupy activated and effector gene expression clusters, resembling spike-specific cells from acute infection.

### Spike-specific CD8 T cell clones are broadly activated in people with LC

Memory T cell responses are composed of different TCR clonotypes ranging from small expansions to highly expanded clones. To assess whether LC was associated with changes in CD8 T cell clonality or distinct differentiation states for individual clones, we identified clones from paired single-cell TCR sequencing of virus-specific T cells. With the exception of cells in cluster 8, we recovered TCR sequences from the vast majority of cells in the dataset (**Fig. S16A**). Naive clusters 2 and 18 were dominated almost exclusively by unexpanded singletons, whereas all non-naive clusters contained a substantial fraction of expanded clones (**Fig. S16A**). Due to the abundance of naive T cells that have not clonally expanded, bulk CD8 T cells had a high inverted Simpson index, indicating contributions from a large number of small clones to the overall repertoire (**Fig. S16B**). By contrast, all virus-specific CD8 T cell populations were clonally expanded. No substantial differences in clonality were observed comparing recovered, acute, and LC T cells targeting the same antigens across subjects (**Fig. S16B-E**).

To explore how gene expression patterns may differ across individual CD8 T cell clones, we mapped the mean UMAP coordinates of all cells from each individual clone as a single point, with points reflecting the size of each clone (**Fig. 4J**). In recovered controls, clone averages mapped to similar regions where individual cells were found for each specificity (**Fig. S16F**). During acute SARS-CoV-2 infection, the majority of non-singlet spike-specific CD8 T cell clones shifted towards activated regions of the UMAP, suggesting cells from almost every clone were stimulated by spike epitopes and experienced activation-associated changes in gene expression. Clones from people with LC were similarly shifted towards activated regions, with more clones located towards the terminal effector cluster 3 region of the UMAP compared to recovered controls or acute infection (**Fig. 4J**). These clones from LC were also nearly absent from the memory-associated cluster 4 (**Fig. 4J**).

To assess clonal activation in a quantitative manner, each expanded spike-specific clone with at least 5 cells was ranked based on the proportion of cells in the clone that were in the activated clusters 6, 7, and 8 (**Fig. 4K**). The least activated clones in the dataset were from recovered controls, whereas the most activated clones all derived from acute infection samples, consistent with the presence of abundant viral antigens during acute infection. Clones from people with LC, however, displayed evidence of activation above what was typical for recovered controls but below that observed for most acute infection samples, consistent with the persistence of spike antigen in LC (**Fig. 4K-L, Fig. S17A**). Activation of EBV lytic antigen-specific T cell clones followed a similar pattern, although the differences were less pronounced compared to spike-specific clones (**Fig. 4M, Fig. S17B-C**). Clones recognizing EBV latent or Flu antigens showed no increase in activation in acute infection or LC compared to recovered controls, and clones from CMV were only modestly more activated in LC (**Fig. S17D-F**). Thus, these data demonstrate that the Spike^Act^ LC subset has spike-specific CD8 T cell populations comprised of many activated clones, consistent with ongoing antigen stimulation.

### Spike-specific CD8 T cells in LC express genes associated with TCR stimulation, effector biology, and T cell exhaustion

To further explore the genes and molecular programs underlying the differential clustering of spike-specific CD8 T cells in LC, we performed DEG analysis of antigen-specific populations from acute infection or LC versus recovered controls. Spike-specific CD8 T cells from acute SARS-CoV-2 infection significantly upregulated over 2000 genes compared to spike-specific CD8 T cells from recovered controls, consistent with antigen-driven activation during acute infection (**Fig. 5A, Fig. S18A**). Spike-specific CD8 T cells from LC samples also had substantial changes in gene expression compared to spike-specific CD8 T cells from recovered controls, with 549 genes significantly upregulated and 138 genes significantly downregulated (**Fig. 5B, Fig. S18A**). No other antigen-specific population had as many DEGs when comparing LC and recovered controls (**Fig. S18A**), suggesting that spike-specific CD8 T cells are substantially altered in LC in a way that is distinct from other CD8 T cell specificities. Smaller changes in gene expression were observed for all other specificities in acute infection compared to recovered controls, consistent with low-level bystander activation (**Fig. S18A-C**). There were relatively few DEGs comparing EBV lytic-specific CD8 T cells during acute SARS-CoV-2 infection to those from recovered controls (**Fig. S18C**). Although less prominent than the changes to spike-specific CD8 T cells in LC, EBV lytic- and latent-specific, and to a lesser extent, CMV-specific CD8 T cells also had a number of DEGs compared to recovered controls in LC, whereas Flu-specific CD8 T cells showed few changes in gene expression in LC compared to recovered controls (**Fig. S18A-C**).

**Fig. 5:**
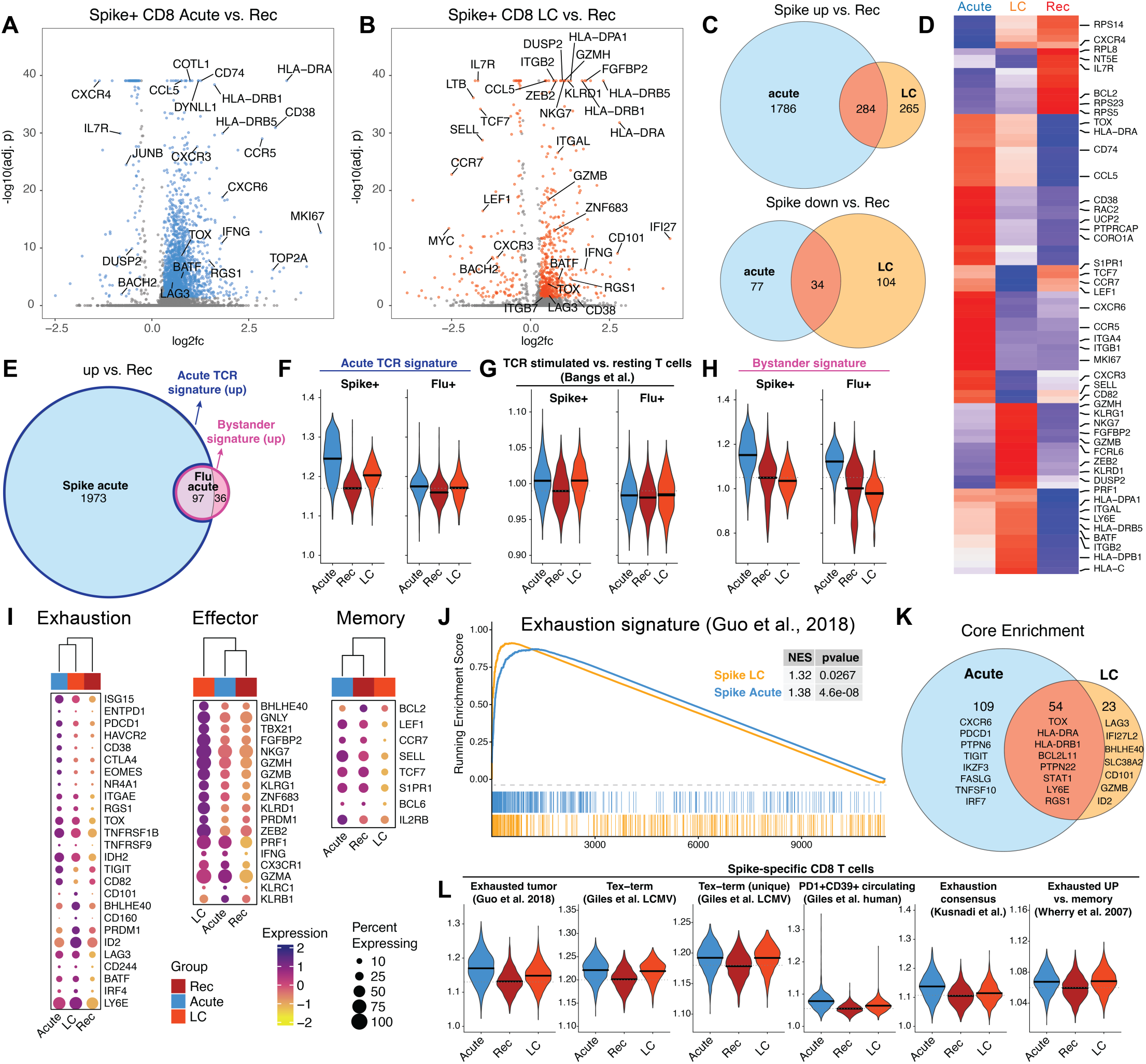
Spike-specific CD8 T cells in LC express genes associated with TCR stimulation, effector biology, and T cell exhaustion. (A-B) Volcano plots displaying significantly upregulated and downregulated DEGs in spike-specific CD8 T cells from (**A**) acute infection or (**B**) LC compared to recovered controls. Significant DEGs were identified via the Seurat FindMarkers function. (**C**) Venn diagrams displaying the shared and distinct significantly upregulated (top) and downregulated (bottom) genes from spike-specific CD8 T cells from acute infection and LC versus recovered controls. (**D**) Clustered heatmap of average expression of selected genes by spike-specific CD8 T cells from each of the six distinct regions in (C). (**E**) Venn diagram displaying the shared and distinct significantly upregulated genes for spike- and Flu-specific CD8 T cells from acute infection compared to recovered controls. Along with downregulated genes (Fig. S20E), these genes were used to derive signatures of acute TCR stimulation and bystander stimulation as indicated. (**F-H**) Summary data depicting module scores for spike-or Flu-specific CD8 T cells from the indicated groups. Module scores were calculated using the UCell package. For (F) and (H), upregulated and downregulated modules were derived separately and combined by subtracting the downregulated score from the upregulated score after normalizing by the number of genes in each module. For (G), genes from a published signature that were upregulated after TCR stimulation were used to derive the score(*66*). (**I**) Clustered dot plots displaying the expression of selected genes associated with CD8 T cell memory differentiation, effector differentiation, or exhaustion. (**J**) Gene set enrichment analysis of spike-specific CD8 T cells from acute infection or LC compared to recovered controls, using a published gene set of T cell exhaustion(*68*). (**K**) Venn diagram depicting the overlap in genes driving the core enrichment for the gene set in (J) for spike-specific CD8 T cells from acute infection or LC compared to recovered controls. Selected genes are labelled within each category. (**L**) Summary data depicting module scores for spike-specific CD8 T cells from the indicated groups. Module scores were calculated using the UCell package using six distinct published gene sets of T cell exhaustion, including genes upregulated in exhausted CD8 T cells within non-small cell lung cancer tumors(*68*), terminally exhausted CD8 T cells from LCMV clone 13 infection compared to all other cells(*70*), terminally exhausted CD8 T cells from LCMV clone 13 infection and not upregulated in effector CD8 T cells(*70*), circulating PD-1+ CD39+ CD8 T cells in humans(*69*), a consensus signature of T cell exhaustion(*71*), and a signature of exhausted T cells compared to memory T cells in LCMV(*72*). All analyses in Fig. 5 were performed after downsampling each antigen-specific population to a maximum of 150 cells per sample to avoid bias associated with highly expanded responses. Significant DEGs were defined as those with an adjusted p value below 0.05 and log2fc above 0.3 or below -0.3.

To investigate whether there were any shared molecular programs for antigen-specific CD8 T cell populations in acute SARS-CoV-2 infection and LC, we compiled all of the upregulated or downregulated DEGs from spike-, Flu-, EBV lytic-, and CMV-specific cells from acute or LC samples compared to recovered controls. Of the 549 genes upregulated in spike-specific CD8 T cells from LC, 284 were also significantly upregulated by spike-specific cells during acute infection, and 34 of the 138 downregulated genes were also downregulated during acute infection (**Fig. 5C**). This overlap was far greater than that observed for DEGs for Flu-specific (**Fig. S18D**) or EBV lytic-specific CD8 T cells (**Fig. S18E**) from acute infection versus LC. Moreover, across all DEGs from different antigen-specific populations, the most prominent overlapping upregulated and downregulated patterns were genes shared only by acute spike-specific and LC spike-specific CD8 T cells (**Fig. S19A-B**). Thus, across the entire dataset, the two populations that were most similar in their deviation from recovered controls were spike-specific CD8 T cells responding to acute SARS-CoV-2 infection and spike-specific CD8 T cells from LC.

We next investigated the molecular programs that were shared or distinct between spike-specific CD8 T cells from acute infection and LC. The overlapping upregulated genes included many genes associated with T cell activation or exhaustion, including HLA-II genes (HLA-DRA, HLA-DRB5, HLA-DPA1, etc.), CD74, CCL5, LY6E, and BATF (**Fig. 5D**). Many of the shared downregulated genes are known to have lower expression after acute TCR stimulation, including NT5E (CD73), IL7R, and BCL2, as well as genes associated with translation (RPL8, RPS20, RPS5, etc.). Genes associated with proliferation, including MKI67, were only upregulated by spike-specific CD8 T cells from acute infection, suggesting that cells from LC may be less proliferative despite engaging other activation programs. Several genes associated with T cell trafficking were also differentially regulated, with CXCR3, CXCR6, CCR5, ITGA4, and ITGB1 more highly expressed in acute infection, and ITGB7, CD69, ZNF683, ITGB2, and ITGAL more highly expressed in LC, suggesting that these cells may have been trafficking from different anatomical locations when they were activated *in vivo* (**Fig. 5D, Fig. S20A**). Genes that were uniquely upregulated in spike-specific cells from LC but not acute infection included many associated with cytotoxicity and terminal effector biology (GZMB, GZMH, FGFBP2, FCRL6, SEB2, DUSP2), and uniquely downregulated genes in LC reflected the loss of a stem-like memory T cell program (TCF7, CCR7, LEF1, S1PR1), which was retained during acute infection despite antigen-driven activation (**Fig. 5D**).

To further interrogate these gene expression profiles, we once again performed gene set enrichment analysis of GEPs from T-CellAnnoTator (TCAT) on ranked gene sets comparing spike-specific CD8 T cells from acute infection or LC versus recovered controls. Spike-specific CD8 T cells from acute infection were highly enriched for several activation and cell cycle-associated GEPs and depleted of GEPs associated with translation and naive/resting memory T cells compared to cells from recovered controls (**Fig. S20B**). Spike-specific CD8 T cells from LC were depleted for the same naive/resting memory GEPs and were strongly enriched for the cytotoxic GEP and somewhat enriched for some activation and cell cycle GEPs (**Fig. S20B**). In contrast, no GEPs from TCAT were significantly enriched in the comparisons of Flu-or EBV lytic-specific CD8 T cells in either acute infection or LC compared to recovered controls (**Fig. S20C-D**). When comparing spike-specific CD8 T cells from acute infection to those from LC, fewer GEPs were enriched than when either population was compared to cells from recovered controls, suggesting that spike-specific CD8 T cells from LC were more similar to spike-specific cells from acute infection than either were to cells from recovered controls. Spike-specific CD8 T cells from LC were enriched for cytotoxicity compared to cells from acute infection, which were alternatively enriched for central memory genes (**Fig. S20B**). GEPs associated with activation and proliferation were only modestly enriched in spike-specific CD8 T cells from acute infection compared to those from LC (**Fig. S20B**). In summary, spike-specific CD8 T cells from LC upregulated genes associated with activation and TCR stimulation, engaged a cytotoxic effector program, and lost the transcriptional programs associated with proliferation and stem-like memory.

T cells are most potently activated by stimulation via the TCR after binding to peptide/HLA but can also undergo bystander activation in response to cytokines and other inflammatory signals(*65*). We reasoned that we could derive bystander and TCR-driven activation signatures from this dataset, with Flu- and spike-specific T cells both responding to the overall inflammatory environment during acute infection, but only spike-specific T cells receiving TCR stimulation from spike peptides. Indeed, 97 of the 133 genes significantly upregulated by Flu-specific CD8 T cells during acute SARS-CoV-2 infection were also upregulated by spike-specific CD8 T cells (**Fig. 5E**). However, spike-specific CD8 T cells upregulated many additional genes that were not shared with Flu-specific CD8 T cells, which could likely be attributed to TCR-driven stimulation. Using all significantly upregulated (**Fig. 5E**) and downregulated (**Fig. S20E**) genes by Flu-specific cells during acute SARS-CoV-2 infection, we derived a bystander T cell activation signature. Similarly, genes uniquely upregulated or downregulated by spike-specific CD8 T cells were used to derive a signature of acute TCR stimulation (**Fig. 5E, Fig S20E**). Spike-specific CD8 T cells from LC strongly enriched for the acute TCR signature compared to recovered controls (**Fig. 5F**). A similar enrichment was not observed for Flu-specific CD8 T cells from LC or acute infection. The acute TCR signature was only minimally elevated during acute infection for EBV-or CMV-specific CD8 T cells, with no increase in LC compared to recovered controls (**Fig. S21A**). Two published signatures of TCR stimulation were similarly upregulated most prominently by spike-specific CD8 T cells during acute infection and LC (**Fig. 5G, Fig. S21B-C**)(*66, 67*). In contrast, the bystander signature was upregulated by spike-, Flu-, and all other CD8 T cell specificities during acute SARS-CoV-2 infection (**Fig. 5H, Fig. S21D**). None of the CD8 T cell specificities examined were enriched for this signature in LC, however, indicating that T cell activation in LC is distinct from the bystander activation signatures found during acute viral infection. Taken together, these data support the conclusion that spike-specific CD8 T cells in LC were persistently activated via TCR stimulation rather than bystander inflammation.

Chronic TCR stimulation is a driver of CD8 T cell exhaustion. Given that persistent TCR-driven activation appeared to be a feature of spike-specific CD8 T cells in LC, we explored the expression patterns of known effector, memory, and exhaustion transcriptional programs. As previously noted, spike-specific CD8 T cells from LC downregulated many genes associated with stem-like/central memory T cells, including LEF1, CCR7, SELL, TCF7, S1PR1, and BCL6 (**Fig. 5I**). Despite being potently activated, spike-specific CD8 T cells from acute SARS-CoV-2 infection retained expression of these genes. Together with loss of memory T cell genes, spike-specific CD8 T cells from LC had higher expression of many genes associated with cytotoxic and terminal effector CD8 T cells, including BHLHE40, TBX21, FGFBP2, GZMB, ZNF683, and ZEB2 (**Fig. 5I**). Both exhausted and acutely stimulated T cells often have high expression of exhaustion-associated genes, as a main driver of T cell exhaustion is chronic T cell activation. Indeed, many exhaustion-related genes had elevated expression in spike-specific CD8 T cells from acute infection and LC compared to recovered controls (**Fig. 5I**). Spike-specific CD8 T cells from acute infection had the highest expression of ISG15, HAVCR2, CD38, CTLA4, ITGAE, RGS1, and TOX, although these genes were also upregulated in LC. In contrast, CD160, BHLHE40, PRDM1, ID2, LAG3, and CD244 (2B4) were preferentially upregulated in spike-specific CD8 T cells from LC (**Fig. 5I**). Overall, these gene expression profiles were consistent with CD8 T cell exhaustion but could not definitively distinguish exhaustion from other activated cell states.

To more quantitatively assess whether exhaustion transcriptional programs were enriched in spike-specific cells in LC, we performed GSEA analysis using a T cell exhaustion gene set from non-small-cell lung cancer (**Fig. 5J**)(*68*). Compared to recovered controls, spike-specific CD8 T cells from both acute infection and LC were strongly enriched for this exhaustion signature. Core enrichment analysis revealed 54 shared genes that drove enrichment in spike-specific CD8 T cells from both acute infection and LC, including the key exhaustion transcription factor TOX (**Fig. 5K**). This shared enrichment likely reflects the overlap of T cell exhaustion with T cell activation. In addition to 109 genes that were only in the leading edge in the acute condition, 23 genes were unique to the leading edge in LC, suggesting that spike-specific CD8 T cells from LC upregulate some aspects of the exhaustion program beyond those associated with acute activation. These genes included LAG3, BHLHE40, CD101, and ID2. Similar analysis with a second exhaustion gene set from circulating human PD-1+CD39+ CD8 T cells(*69*) again revealed strong enrichment by spike-specific CD8 T cells in both acute infection and LC compared to recovered controls (**Fig. S22A-C**). We then derived module scores from these and 4 additional exhaustion gene sets and asked which virus-specific CD8 T cell populations enriched for exhaustion signatures(*70–72*). Spike-specific CD8 T cells from LC were enriched for all exhaustion signatures tested compared to recovered controls, whereas no other specificities consistently enriched for exhaustion signatures in LC (**Fig. 5L, Fig. S22D-H**). Taken together, these data indicate that spike-specific CD8 T cells from people with LC express genes associated with TCR-driven activation, terminal effector differentiation, and T cell exhaustion, consistent with potential chronic antigen stimulation.

### Spike-specific CD8 T cells that are persistently activated in LC express TOX and 2B4 and have reduced cytokine production, survival, and proliferative potential

To more directly test whether spike-specific CD8 T cells form LC had features of T cell exhaustion, we performed three complementary assays to measure functionality and T cell exhaustion in 46 samples from the LIINC cohort (**Fig. S23A**). First, we analyzed PBMCs using an additional spectral flow cytometry panel including HLA-I/peptide tetramers with additional markers of activation and exhaustion. Samples from people with LC were divided into two groups based on the level of spike-specific CD8 T cell activation observed in prior experiments (Spike^Act^ LC and Spike^norm^ LC). The Spike^Act^ LC samples had significant increases in activated (CD45RA-CD38/HLA-DR+, HLA-DR+2B4+) and exhausted-like (PD-1+TOX+, PD-1+2B4+, TOX+2B4+) cells compared to recovered controls (**Fig. 6A-C**). These cells also had higher expression of HLA-DR, CD38, and TOX and trended towards lower expression of CD45RA and higher expression of 2B4, all of which are associated with T cell activation and/or exhaustion (**Fig. S23B**). Consistent with the sequencing data, spike-specific CD8 T cells from Spike^Act^ LC trended towards lower expression of CD127, in contrast to highly activated cells from acute infection that maintained higher CD127 expression (**Fig. 6A, Fig. S23B**). In contrast, Spike^norm^ LC samples resembled recovered controls. Thus, a subset of people with LC had higher expression of markers associated with T cell activation and exhaustion.

**Fig. 6:**
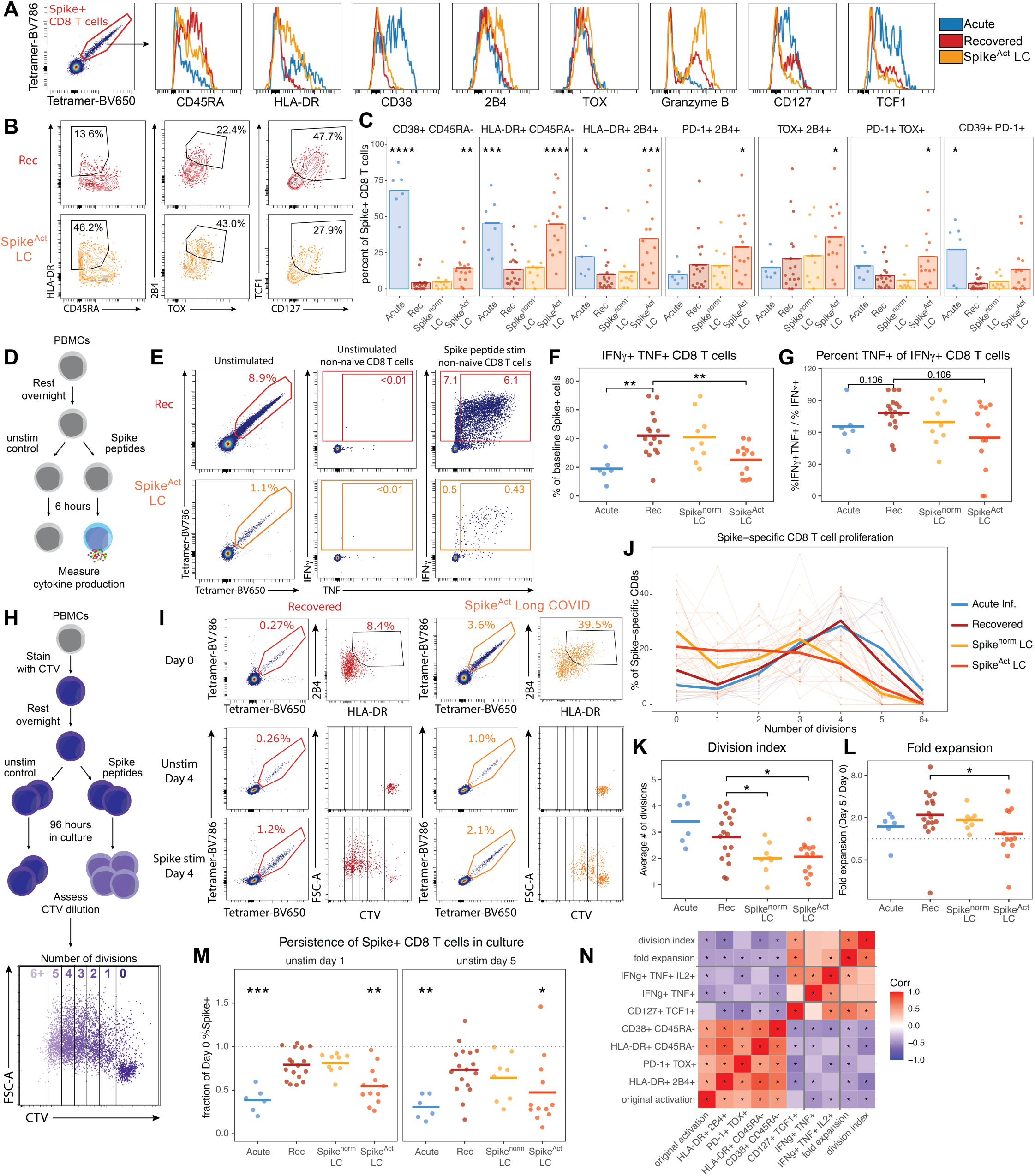
Spike-specific CD8 T cells that are persistently activated in LC express TOX and 2B4 and have reduced cytokine production, survival, and proliferative potential. (**A**) Representative gating of spike-specific CD8 T cells (left) and overlaid histograms of concatenated flow cytometry data from spike-specific CD8 T cells from all samples in the indicated groups. (**B**) Concatenated flow cytometry plots from spike-specific CD8 T cells from all samples in the indicated groups. For visualizations in (A) and (B), each sample was downsampled to a maximum of 150 spike-specific CD8 T cells to avoid bias associated with highly expanded responses. (**C**) Summary data depicting the percent of spike-specific CD8 T cells in the indicated gated populations. (**D**) Schematic representation of 6-hour stimulation assays to measure the cytokine production capacity of spike-specific CD8 T cells. (**E**) Representative gating of spike-specific CD8 T cells at baseline (left) and cytokine producing cells after 6h stimulation with or without spike peptides (right). Cytokine production was measured on total non-naive CD8 T cells, subtracting non-specific background staining from paired unstimulated controls and normalizing to the tetramer+ frequency at baseline. (**F-G**) Summary data of the frequency of spike-specific CD8 T cells producing IFNγ and TNF (F) and the fraction of IFNγ+ cells also producing TNF (G). (**H**) Schematic representation of CTV staining and 96-hour stimulation assays to measure the proliferative potential of spike-specific CD8 T cells. (**I**) Representative gating of spike-specific CD8 T cells at baseline (top) and after 96-hour stimulation with or without spike peptides (bottom). The phenotype of baseline spike-specific CD8 T cells is depicted with biaxial gating of HLA-DR+ 2B4+ cells, and the proliferation of spike-specific CD8 T cells is captured by the progressive dilution of CTV with each cell division. (**J**) Summary data showing the percent of spike-specific CD8 T cells that had completed the indicated number of cell divisions based on dilution of CTV staining. Thin lines connect individual samples, thick lines are the average across participants within a group. (**K-L**) Summary data showing (K) the division index (average number of cell divisions) or (**L**) the fold expansion (spike-specific CD8 T cell frequency after 96h stimulation divided by frequency at baseline) of spike-specific CD8 T cells from each sample. (**M**) Summary data showing the persistence of spike-specific CD8 T cells in unstimulated culture wells, calculated as the frequency of tetramer+ cells observed after culture for 1 or 5 days, divided by the frequency observed in baseline samples. (**N**) Matrix of Spearman rank correlations among baseline spike-specific CD8 T cell differentiation states (as a percentage of the spike-specific CD8 T cell population) and functional assay measurements. All recovered and long COVID samples were included in correlation analyses. All statistical tests were Wilcoxon tests with Benjamini-Hochberg correction for multiple comparisons. *p < 0.05, **p < 0.01, ***p < 0.001, ****p < 0.0001. In the absence of brackets, all statistical tests are comparisons to recovered controls.

We next sought to directly investigate the functionality of spike-specific cells from these people with LC. Exhausted CD8 T cells have a reduced capacity to produce cytokines and proliferate in response to antigen stimulation, as well as poor survival in the absence of antigen compared to memory CD8 T cells(*43*). To first test whether spike-specific CD8 T cells from people with LC had deficiencies in cytokine production, cells were stimulated with the same spike peptides used in HLA-I/peptide tetramers or left unstimulated for 6 hours, followed by intracellular cytokine staining (**Fig. 6D**). The background-subtracted frequency of cells producing cytokines after spike peptide stimulation was compared to the frequency of spike tetramer+ CD8 T cells in the paired samples stained directly *ex vivo*. Spike-specific CD8 T cells from Spike^Act^ LC samples had a significant decrease in the ability to produce both TNF and IFNγ, as did spike-specific cells from acute infection (**Fig. 6E-F**), with a trend towards decreased poly-functionality (the fraction of IFNγ-producing cells that also produced TNF, **Fig. 6G**).

From the same samples, we also assessed the proliferative capacity of spike-specific CD8 T cells by staining with CellTrace Violet (CTV) and stimulating these cells with spike peptides for 4 days (**Fig. 6H**). T cells from unstimulated wells showed no dilution of CTV, confirming that proliferation in these experiments is dependent on peptide stimulation (**Fig. 6I**). In contrast, spike tetramer+ cells stimulated with spike peptides diluted CTV and increased in size. There was minimal background proliferation in tetramer-populations. Spike-specific CD8 T cells from recovered subjects proliferated extensively over 4 days, with very few cells remaining undivided and most cells having undergone 3-5 divisions (**Fig. 6I-J, Fig. S23C**). In contrast, although some spike-specific CD8 T cells from people with LC proliferated, there were more cells that remained undivided during 4 days of stimulation compared to recovered controls (**Fig. S23C**). Moreover, the cells that did divide did not undergo as many divisions as cells from recovered controls or acute infection samples (**Fig. 6I-J**). As a result, spike-specific CD8 T cells from LC samples had a significantly reduced division index compared to cells from acute or recovered samples (**Fig. 6I-K**). In contrast, spike-specific CD8 T cells from acute infection proliferated rapidly, with comparable or greater division compared to recovered controls. These data suggest that spike-specific cells from people with LC may be dysfunctional beyond what is observed during acute activation, a pattern characteristic of T cell exhaustion in the setting of chronic antigen stimulation.

Moreover, spike-specific CD8 T cells from Spike^Act^ people with LC had less numerical accumulation after 4 days of stimulation compared to cells from recovered controls (**Fig. 6L**). The decrease in expansion was multifactorial, as spike-specific CD8 T cells from Spike^Act^ LC had both fewer cell divisions and poorer survival in unstimulated wells compared to cells from recovered controls (**Fig. 6M**). The survival defect was driven by a decrease in cell recovery after the first day of culture and was also observed for spike-specific CD8 T cells from acute infection. This poor survival may be attributed to the death of short-lived effector-like CD8 T cells. Finally, to interrogate the relationships between CD8 T cell differentiation states and functionality, we asked whether the percent of spike-specific CD8 T cells that had activated or exhausted phenotypes directly *ex vivo* was correlated with cytokine production and proliferation independent of LC status. Excluding samples from acute infection, protein expression patterns consistent with activation and/or exhaustion were negatively correlated with cytokine production and proliferation, whereas the abundance of CD127+ TCF1+ memory T cells was positively correlated with these functional readouts (**Fig. 6N, Fig. S23D**). Taken together, these data demonstrate that a subset of people with LC have spike-specific CD8 T cells that express markers associated with exhaustion and produce fewer cytokines, have lower proliferative potential, and have decreased survival compared to memory CD8 T cells present in recovered individuals. These features are consistent with the presence of exhausted or partially exhausted spike-specific CD8 T cells in a subset of patients with LC and are indicative of chronic stimulation by a persistent reservoir of SARS-CoV-2.

## Discussion

This study identified a subset of people with LC who maintained persistent activation of SARS-CoV-2 spike-specific CD8 T cells for months to years after the initial SARS-CoV-2 infection. Spike-specific CD8 T cells from these patients had an increase in cytotoxic effector biology, a decrease in memory T cell transcriptional programs, and expressed genes associated with T cell activation and exhaustion. Moreover, these cells expressed multiple activation markers and were poorly functional, producing fewer cytokines and proliferating less than spike-specific CD8 T cells from recovered controls. Thus, analysis of spike-specific CD8 T cells suggests that chronic stimulation by SARS-CoV-2 antigens occurs in a subset of people with LC and alters T cell differentiation. This finding supports the existence of a persistent tissue reservoir of SARS-CoV-2 that can be accessed by a fraction of the circulating CD8 T cells at any given time. It remains unclear, however, where this reservoir may reside and why the activation signal observed in LC is less pronounced than during acute infection. The gut may be a SARS-CoV-2 reservoir(*24*), and spike-specific CD8 T cells from LC samples expressed higher levels of LFA-1 and ITGB7 compared to those from recovered and acute samples, consistent with possible trafficking of these cells to and/or from the inflamed intestinal mucosa. Additional studies will be necessary to link T cell activation to direct measures of viral persistence.

Activation and exhaustion of spike-specific CD8 T cells were pronounced in a subset of people with LC, but most participants had low activation comparable to recovered controls. This observation is consistent with a model in which multiple mechanisms contribute to LC, and SARS-CoV-2 persistence drives disease in only a fraction of patients. Alternatively, it is possible that many people with LC harbor a persistent SARS-CoV-2 reservoir that does not stimulate CD8 T cells, either because the virus resides in anatomic sites that are inaccessible to CD8 T cells or because CD8 T cells exerted sufficient pressure to drive selection for escape mutants that no longer stimulate these CD8 T cells. We did not observe consistent activation of spike-specific CD4 T cells in LC, but this requires further validation, as these analyses were performed with fewer epitopes and a smaller cohort compared to the CD8 T cell analyses. However, it is possible that persistent SARS-CoV-2 antigens may be presented more efficiently by HLA-I than by HLA-II, or that activated CD4 T cells are retained in tissues and are difficult to capture in blood, resulting in minimal CD4 T cell activation despite robust stimulation of spike-specific CD8 T cells in LC.

The cutoff used in this study identified 40% of people with LC and 16% of recovered controls with elevated spike-specific CD8 T cell activation. These frequencies are aligned with several reports indicating that SARS-CoV-2 persistence is enriched in LC, but is neither universal among people with LC nor absent in those who report full recovery(*6, 7, 16, 21, 23, 24*). No discrete cutoff in spike-specific CD8 T cell activation uniquely identifies people with LC. Rather, persistence of SARS-CoV-2 and subsequent stimulation of spike-specific CD8 T cells may exist on a spectrum, where some degree of viral persistence and corresponding CD8 T cell stimulation may occur without driving prolonged symptoms.

CD8 T cells specific for EBV lytic antigens were also activated in LC, largely in the same participants who had elevated spike-specific CD8 T cell activation. These data nonetheless support an association between EBV activity and the development of LC. CMV-specific CD8 T cells did not show substantially elevated activation in LC, but this should be explored further in larger studies of CMV seropositive individuals. Notably, herpesvirus reactivation could mechanistically link LC to other post-acute infectious sequelae(*73*), such as ME/CFS(*74–76*) and post-treatment Lyme Disease(*77*), as herpesvirus infections are ubiquitous and reactivation can be triggered by many stimuli, including inflammation during infection (*34, 35, 78*). Moreover, EBV is associated with the development of autoimmunity, and poor control of EBV may underly observations that autoantibodies can be involved in LC pathogenesis(*15, 79*). Together, these data implicate EBV reactivation more strongly than reactivation of CMV or VZV. More comprehensive efforts examining T cells specific for other members of the human virome(*36*) will be needed to fully resolve how the activity of common viruses shapes LC and other infection-associated chronic conditions.

Our findings also point toward therapeutic strategies that could be tested in clinical trials. Due to a lack of proven biomarkers, LC trials currently enroll and assess response without any objective measure of the mechanism being targeted. A useful biomarker would be measurable in blood, tied to a proposed disease mechanism, and stable within an individual over time. Activation of virus-specific CD8 T cells meets each of these criteria and could therefore be used to stratify participants in downstream analyses, identify those most likely to benefit from interventions targeting SARS-CoV-2 persistence or EBV reactivation, or as a readout of intervention efficacy through changes in activation or exhaustion. It remains unknown whether activated T cells contribute directly to LC pathogenesis or are simply biosensors of persistent viral antigens that drive disease by other mechanisms. Because the chronic T cell activation we observed in LC appears to be driven by specific viral antigens, the most direct therapeutic approach is to remove the antigen. Although antiviral trials in LC have not yet shown efficacy(*63, 80–82*), stratifying participants by high versus low activation may reveal better responses for those with ongoing antigen exposure. Similarly, efforts to reduce EBV antigens, either with antivirals or by promoting host control of EBV, could be pursued. Existing therapies could also be used to modulate activated T cell responses in multiple ways. If T cell stimulation is a pathogenic driver of LC, interventions aimed at dampening T cell activation or reducing effector function might be effective in some people. Alternatively, despite having reduced functionality, exhausted spike-specific CD8 T cells may partially suppress SARS-CoV-2 replication and limit disease progression. In this case, immune checkpoint blockade (e.g., anti-PD-1) or other attempts to reinvigorate exhausted CD8 T cells could facilitate more efficient viral control or reservoir clearance. Because the relative contributions of these mechanisms are unknown, combination approaches trialed via factorial designs may offer the best path to disentangling the biology and achieving meaningful clinical outcomes (*4*). Beyond understanding and treating LC, these findings support expanding the concept of using T cells as antigen biosensors to drive therapeutic advances in other settings.

In summary, we describe a subset of long COVID patients with evidence of virus-specific immune stimulation that is quantifiable and associated with T cell dysfunction. This provides key mechanistic insights into the disease processes that may be driving LC and identifies a readily sampled biomarker for stratifying patients in clinical trials, assessing response to treatment, and potentially guiding therapies to modulate T cell responses as a treatment for LC.

## Acknowledgments

We would like to thank the study participants, especially members of the long COVID patient community, for their dedication to research and for providing the specimens that made this study possible. We also thank Florin Tuluc and the Flow Cytometry Core at the Children’s Hospital of Philadelphia and the Penn Cytomics and Cell Sorting Resource Laboratory for access to instruments, the NIH Tetramer Core Facility for providing HLA-II tetramers, and members of the Wherry lab for helpful discussions and feedback.

## Author contributions

MMP and EJW conceived the study. MMP, DBR, and NRD carried out experiments. MMP and DBR analyzed data and drafted the initial manuscript. All authors provided input and edits to the manuscript. MMP, DBR, and EJW prepared the final manuscript. MMP, TD, MB, EAF, DR, MCW, A-KR,GB-Z, ZA, JPR, DKS, MOH, ST, and BB performed clinical recruitment, managed sample processing and storage, and managed sample metadata. MMP, DBR, NRD, SGG, SM, and JRG contributed to the methodology. JRG, SM, and JNM provided input on statistical analyses. TD, MB, EAF, DR, MCW, A-KR,GB-Z, ZA, JPR, DKS, MOH, ST, BB, DP, JDK, JNM, BAA, NJM, AI, TJH, MJP, and SGD provided data and materials. MMP, DP, JDK, JNM, BAA, NJM, AI, TJH, MJP, SGD, and EJW supervised cohorts. EJW supervised the study.

## Data, code, and materials availability

All source data, data files, and code are available from the authors upon request.

