## Supplementary Materials for "Persistently activated virus-specific CD8 T cells indicate antigen-driven immune aberrations in long COVID"

**The PDF file includes:**

Materials and Methods

Figs. S1 to S23

Tables S1 to S6

References #83-89

### **Supplementary Materials**

#### **Materials and Methods:**

Details on all key resources can be found in **Table S6**.

#### **Human subject recruitment and sampling**

Human subjects from six cohorts were recruited for this study: five (COVEND, MESSI, AALC, LIINC, MY-LC) were included in the primary analysis, and one (OUTSMART-LC), associated with a clinical trial, was used to generate a parallel dataset. Detailed demographics for each cohort are provided in Tables S1-S3 and Table S5, and cohort descriptions are as follows.

**COVEND:** Participants (N = 48) were recruited at the University of Pennsylvania to study the development of immunity to SARS-CoV-2 in longitudinal samples. At enrollment, a detailed history of all known SARS-CoV-2 infections and vaccine doses was obtained, and each participant was followed closely to document additional vaccine doses or reinfections. Participants were encouraged to perform at-home rapid tests for SARS-CoV-2 whenever they had symptoms of a respiratory infection. All participants were otherwise healthy, with no self-reported history of chronic health conditions, and none were hospitalized during SARS-CoV-2 infection. Peripheral blood samples (30–100 ml) and clinical questionnaire data, including questions aimed at capturing any potential LC symptoms, were collected at each study visit. PBMCs were isolated as previously described(51). Study visits occurred every 3 months (serial monitoring) and at approximately days 7, 15, and 45 after each new SARS-CoV-2 infection or vaccination. Samples from these individuals were used for acute vaccination (N = 22), non-hospitalized acute infection (N = 20), and recovered control (N = 23) groups and paired with samples from the MESSI and AALC cohorts to form the University of Pennsylvania cohort used to study both CD4 and CD8 T cells in this study. This study was approved by the University of Pennsylvania Institutional Review Board (IRB, protocol number 851465).

**Molecular Epidemiology of Sepsis in the ICU (MESSI):** Participants (N = 27 for this study) hospitalized for acute SARS-CoV-2 infection or a presumed COVID-19-related complication between June 2020 and August 2021 were enrolled in a molecular cohort study within 3 days of hospital admission and a positive SARS-CoV-2 test (PCR or rapid antigen test). Informed consent was obtained from participants or their legally authorized representatives. Blood was collected and processed for peripheral blood mononuclear cells (PBMCs) as previously described(83, 84). For these analyses, a total of 47 samples, collected from the day of admission up to 27 days thereafter, were assigned to the acute infection group for CD8 T cell analyses. The study was approved by the University of Pennsylvania IRB (protocol number 808542).

**Autoantibodies in Patients with Long-COVID (AALC):** Participants (N = 44 for CD8 T cell analyses, N = 29 for CD4 T cell analyses) were recruited from an outpatient Post-COVID Assessment and Recovery Clinic at a large, urban academic medical center, the Penn Medicine Post-COVID Assessment and Recovery Clinic. The study was approved by the University of Pennsylvania IRB (protocol number 852269). Inclusion criteria were age over 18, confirmed SARS-CoV-2 infection greater than or equal to 6 months ago, and self-reported fatigue and cognitive symptoms. Exclusion criteria included pregnancy, prior hospitalization for acute COVID-

19, or inability to provide informed consent. A full history of known SARS-CoV-2 infections and vaccine doses was collected to ensure that all LC samples were collected at least 45 days after the most recent SARS-CoV-2 antigen encounter (minimum observed = 79 days). Blood samples were collected during an in-person visit by a study nurse or trained study staff. PBMCs from a single visit per participant were isolated from heparin tubes using the same protocol as the COVEND cohort.

##### **Long-term Impact of Infection with Novel Coronavirus (LIINC, NCT04362150):**

LIINC is a prospective observational cohort at the University of California, San Francisco (UCSF) that follows individuals after SARS-CoV-2 infection confirmed by nucleic acid amplification testing(85). Participants were referred by clinicians or self-referred and enrolled 14-90 days after symptom onset. Visits took place monthly during the first 4 months post-symptom onset and every 4 months thereafter. At each visit, a structured clinical interview recorded demographics, COVID-19 status, comorbidities and medications, as well as ongoing symptoms and quality of life. In addition, biospecimens were collected and stored(85). Isolation of PBMCs was performed as described previously(85). The study was approved by the UCSF IRB (protocol number 20-30479), and all participants provided written informed consent.

For this study, we included LIINC participants with PBMC samples available at both 4 and 12 months after symptom onset, comprising patients with LC (N = 59) and recovered controls (N = 20). LC status was blinded until data collection was complete.

##### **OUTSMART-LC (NCT05877507):**

OUTSMART-LC was a phase 2a, randomized, double-blind, placebo-controlled trial of the SARS-CoV-2-specific monoclonal antibody AER002 in adults with LC(63). Participants were recruited from the LIINC cohort and assessed for eligibility at a separate screening visit. LC was defined as at least two moderate or severe symptoms that had newly appeared or worsened after a test-confirmed SARS-CoV-2 infection occurring before 15 August 2022, had persisted for at least 60 days, and could not be explained by another cause. Further details of recruitment and clinical assessment are listed in the initial trial report(63). Participants were randomized 2:1 to a single intravenous infusion of AER002 or placebo and followed up approximately 14, 30, 90, 180, and 360 days after infusion. Blood samples were collected before treatment at screening, on the day of infusion, and at follow-up visits. PBMCs were isolated as previously described(85).

Since the intervention showed no benefit over placebo, we pooled both treatment arms and analyzed them as a single longitudinal LC cohort (N = 31). Analysis separated by treatment and symptom improvement can be found in Fig. S7. The treatment group and symptom response data for all OUTSMART-LC samples were blinded until all data were collected.

**MY-LC:** MY-LC is a cross-sectional study at Mount Sinai and Yale that enrolled five groups with different SARS-CoV-2 exposure histories and LC status(12). All participants were $\geq 18$ years old and had a history of confirmed or probable SARS-CoV-2 infection. Further inclusion criteria were symptoms persisting for more than 6 weeks after onset of COVID-19 for participants with LC, and the absence of ongoing symptoms attributed to a previous SARS-CoV-2 infection for recovered controls. Exclusion criteria were current pregnancy, immunosuppression at or above the equivalent of 5 mg prednisone daily, active malignancy or chemotherapy, monogenic disorders, and inability to consent or to undergo blood sampling. During study visits, participants provided blood samples and completed symptom surveys. Further details of recruitment, clinical assessment, and sample processing (e.g., isolation of PBMCs) have been described previously(12). The study was approved by the Mount Sinai Program for the Protection of Human Subjects (IRB 20-01758) and the Yale Institutional Review Board (IRB 2000029451), and all participants provided informed consent. For this study, we analyzed a single sample from each of 64 MY-LC participants in total, including people with LC (N = 50) and recovered controls (N = 14).

#### **HLA tetramers for the identification of antigen-specific T cells**

**HLA-I tetramers:** HLA-I tetramers were generated for HLA-A\*02:01 and HLA-A\*03:01 using the biotinylated Flex-T™ Monomer UVX platform (BioLegend), according to the manufacturer's protocol with modifications detailed below. Peptides were custom-synthesized by GenScript with minimum 90% purity, dissolved in DMSO, and stored at -20°C. For peptide exchange, all reagents were equilibrated to 0°C on ice. Peptide stocks were diluted to 400 mg/mL in cold PBS, and 20 µL of diluted peptide was mixed with 20 µL of Flex-T™ Monomer UVX (200 µg/mL) by pipetting in a 96-well V-bottom plate. The plate was sealed, centrifuged at 800xg for 2 minutes (min) at 4°C, unsealed, and illuminated with 366 nm UV light at 4°C for 30 min at a distance of 2–5 cm from the UV lamp. The plate was then covered and incubated for 30 min at 37°C in the dark.

For flow cytometric analyses, tetramers were generated by centrifuging the plate at 800xg for 2 min at 4°C and transferring 30 µL from the top of each well into PCR tube strips. Fluorophore-conjugated streptavidin (0.66 µg) was added, and tetramers were incubated on ice in the dark for 30 min, then stored at 4°C in the dark for up to 2 months. Barcoded tetramers for 10x single-cell sequencing experiments were prepared as described above, with a ratio of 4.5 µL of peptide-exchanged monomer to 1 µL of BEAM PE/oligonucleotide barcode-conjugated streptavidin. Fluorophore and barcode pairs for each viral peptide are listed in Table S4.

**HLA-II tetramers:** DPB1\*04:01 TFEYVSQPFLMDLE (Spike<sub>167-180</sub>) and DRB1\*15:01 NLLLQYGSFCTQLNRAL (Spike<sub>751-767</sub>) tetramers for flow cytometric analyses and 10x single-cell sequencing were produced by the NIH tetramer core facility at Emory University. The respective peptides were custom synthesized by GenScript with minimum 90% purity.

#### **Tetramer and flow cytometry staining**

**CD8 T cell analysis:** PBMC samples were thawed in 10 mL RPMI supplemented with 10% FBS, 2mM L-Glutamine, 100 U/mL Penicillin, and 100 µg/mL Streptomycin (R10). Following a washing step with R10, cells were counted and plated at a concentration of approximately $3 \times 10^6$ cells per well in a 96-well round bottom plate.

For tetramer staining, staining buffer (SB) consisting of brilliant stain buffer (BD) and FACS buffer (PBS containing 2% FBS) at a 1:1 ratio was supplemented with 0.2% milk and 5 µM free D-biotin (Avidity), and all tetramers were added immediately prior to staining at a 1:250 dilution. Tetramers were prepared and maintained as individual stocks and were briefly centrifuged to remove aggregates prior to addition to the tetramer master mix. Cells were resuspended in 50 µL tetramer staining solution and incubated for 20 min at RT. Samples expressing both HLA-A\*02:01 and

HLA-A\*03:01 were divided and split into two wells, and the two responses were analyzed separately. Following a washing step with FACS buffer, Fc receptor blocking was performed using BioLegend Human TruStain FcX (1:1000 in PBS) for 10 min at RT. Cells were then washed and incubated in 50 μL PBS containing Ghost Viability Dye Violet 510 (Tonbo, 1:1000) for 10 min at RT. After a further washing step, surface staining was performed for 30 min at RT using 50 μL of a staining solution containing surface marker antibodies diluted in SB. Surface staining included CD14, CD16, CD19, and CD41a conjugated to V500, used as a dump channel.

Following removal of the staining solution and washing, cells were fixed and permeabilized for 30 min at 4°C using 100 μL fixation/permeabilization solution (eBioscience Foxp3 Transcription Factor Staining Buffer Kit) per well. Cells were washed in 1X Perm buffer, and intracellular staining was performed overnight at 4°C by adding 50 μL of an intracellular staining solution containing antibodies diluted in 1X Perm buffer. Finally, the samples were washed in 1X Perm buffer, resuspended in FACS buffer, and acquired on a Cytex Aurora flow cytometer. Tetramer+ CD8 T cells were first gated on CD8 T cells that were negative for all other tetramer channels before gating for cells positive in the two channels assigned to each specificity.

**CD4 T cell analysis:** The CD4 T cell protocol largely followed the CD8 T cell protocol described above, with several modifications for the use of MHC-II tetramers: prior to tetramer staining, cells were incubated with dasatinib (Cell Signaling Technology, 50 nM in FACS buffer) for 30 min at 37°C to enhance staining efficiency(86). For the subsequent tetramer and chemokine receptor stain, a staining solution consisting of brilliant stain buffer (BD) and FACS buffer at a 1:1 ratio was supplemented with 50 nM dasatinib and antibodies against CXCR3, CXCR5, CCR6, and CCR4. Tetramers stocks generated by the NIH tetramer core were briefly centrifuged prior to use and added in a 1:200 (DRB1\*15:01 S<sub>751</sub>) or 1:250 (DPB1\*04:01 S<sub>167</sub>) dilution. Cells were resuspended in 100 μL staining solution and incubated for 60 min at 37°C. Surface and intracellular staining was then performed as described above for the CD8 T cell analysis.

**Flow cytometry:** Samples were analyzed on a Cytex Aurora spectral analyzer. Cytex SpectroFlow QC beads were used to ensure consistent acquisition conditions over time. Single stained reference controls for spectral unmixing were prepared with ThermoFisher Ultracomp eBeads Plus. Data analysis was performed in OMIQ and R. Activation was determined by expression of at least two activation markers, as described in Fig. S1 and Fig. S2. For CD8 T cells, tetramer+ responses were determined to be below the limit of detection if fewer than 5 cells were detected in the tetramer gate. A healthy donor reference sample with detectable tetramer+ responses to all antigen categories was analyzed in each experimental batch to ensure consistency in activation measurements across batches.

#### **Single-cell RNA sequencing: sample preparation**

**CD8 T cells:** The sample preparation protocol used for single cell RNA sequencing largely corresponds to the tetramer and surface staining workflow described above. To minimize cell stress, experimental steps were performed at 4°C or on ice whenever possible. Specifically, PBMC samples were thawed in 10 mL chilled RPMI supplemented with 10% FBS, 2 mM L-Glutamine, 100 U/mL Penicillin, and 100 μg/mL Streptomycin (cR10). Following a washing step with cR10, cells were counted, washed with MACS buffer (PBS supplemented with 0.5% BSA and

2 mM EDTA), and incubated for 20 min on ice with a tetramer staining solution. The staining solution was prepared by combining MACS buffer and brilliant stain buffer (BD) at a 1:1 ratio supplemented with 0.2% milk and 5 μM free D-biotin (Avidity). All fluorophore-conjugated and BEAM-conjugated tetramers were added immediately prior to staining at a 1:250 dilution. Tetramer stocks were prepared 24 hours before staining and were briefly centrifuged prior to use to remove aggregates. For each specificity, four distinct tetramer reagents were made as described above and added to the master mix: two PE/BEAM barcoded tetramers with distinct BEAM barcodes, a PE fluorophore-conjugated tetramer (which was found to improve the PE enrichment efficiency compared to using PE/BEAM tetramers alone), and an additional fluorophore-conjugated tetramer used for sorting.

After a washing step with MACS buffer, magnetic-bead based enrichment was performed as described previously<sup>(87-89)</sup> to increase the frequency of antigen-specific T cells as a means of reducing sort times and hashtag antibody volumes. Briefly, cell pellets were resuspended in MACS buffer and incubated for 30 min at 4°C with anti-PE magnetic beads (Miltenyi). Subsequently, cells were washed, resuspended in 1 mL MACS buffer and processed through LS columns (Miltenyi) according to the manufacturer's instructions. Enriched antigen-specific CD8 T cells and depleted bulk CD8 T cells were captured separately in 10 mL conical tubes. Approximately $1.5 \times 10^6$ depleted cells were added back into the enriched fraction for each sample to normalize the number of cells in each staining reaction and allow sorting of bulk tetramer-negative CD8 T cells.

Following the enrichment, Fc receptor blocking was performed by incubating bulk and antigen-specific cells with BioLegend Human TruStain FcX (1:1000 in PBS) for 10 min on ice. Cells were washed and incubated in 50 μL PBS containing Ghost Viability Dye Violet 510 (Tonbo, 1:1000) for 10 min on ice. After a further washing step, surface staining was performed for 30 min on ice using 100 μL of a staining solution containing antibodies against CD3, CD4, CD8, CD27 and CD45RA in SB. Donor identity was encoded by adding different hashtag antibodies (BioLegend TotalSeq-C) to stain spike-specific and bulk T cells from each sample at a volume of 3 μL (1.5 μg) per well. Finally, cells were washed twice and resuspended in FACS buffer for sorting.

Sorting of CD8 T cells was performed using a BD FACSAria II. Bulk tetramer-negative CD8 T cells were sorted regardless of naive/non-naive phenotype. Antigen-specific CD8 T cell populations were sorted and pooled with bulk CD8 T cells as indicated in Fig. S12A prior to loading in a 10x Chromium controller according to the 10x Chromium Next GEM Single Cell 5' HT Kit v2 protocol.

**CD4 T cells:** The CD4 T cell protocol largely adhered to the CD8 T cell protocol, with several modifications for the use of MHC-II tetramers. Prior to tetramer staining, cells were washed with MACS buffer and incubated for 30 minutes at RT with dasatinib (50 nM in MACS buffer). MACS buffer containing tetramers (1:200 dilution for DPB1\*04:01 S<sub>167</sub>) as well as 50 nM dasatinib was then added, and cells were stained for 60 min at RT. Tetramer stocks were briefly centrifuged prior to use to remove aggregates.

Following magnetic enrichment as described above, antigen-specificity as well as donor identity were encoded by labeling spike-specific and bulk CD4 T cells with different hashtag antibodies

(BioLegend TotalSeq-C). Antibody stocks were centrifuged prior to use to remove aggregates. Finally, cells were washed and resuspended in FACS buffer for fluorescent activated cell sorting.

Sorting of CD4 T cells was performed using a BD FACSAria II. Naive and non-naive bulk CD4 T cells were sorted based on their expression of CD27 and CD45RA, counted, and recombined at a 1:4 ratio. Subsequently, spike-specific CD4 T cells were sorted, combined with bulk CD4 T cells for a total of roughly 75,000 cells, and loaded in two reactions following the 10x Chromium Next GEM Single Cell 5' HT Kit v2 protocol.

#### **Single-cell RNA sequencing: library preparation and sequencing**

All downstream library preparation steps for CD4 and CD8 T cells were performed adhering to the 10x Chromium Next GEM Single Cell 5' HT Kit v2 protocol. Quality control regarding size and concentration of amplified cDNA was performed using an Agilent Tapestation and a ThermoFisher Qubit. Prior to sequencing, libraries were quantified using a KAPA Library Quantification Kit (KK4824) and pooled thereafter. Next Generation Sequencing was performed by Azenta (Genewiz) on an Illumina NovaSeq.

#### **Single-cell RNA sequencing: processing, quality control, and analysis**

scRNA-seq data were generated using the 10x Cell Ranger pipeline (9.0.0) and GRCh38 genome. Specifically, fastq files were created using cellranger mkfastq, reads were quantified using cellranger multi, and samples were combined using cellranger aggr. Analysis of processed sequencing data was performed in R (version 4.6.1) using the Seurat v5 package using default parameters unless otherwise noted. Cells from different reactions on the same 10x chip were combined in Seurat using the merge function, and batches (for CD8 T cells) were joined using JoinLayers.

**CD8 T cell dataset:** For the CD8 T cell dataset, cells with less than 1,200 or more than 10,000 reads (nCount\_RNA), cells with more than 3500 detectable genes (nFeature\_RNA), and cells with more than 7.5% of reads mapping to mitochondrial genes (percent\_mt) were excluded. TCR genes were excluded from the gene expression data to avoid clonotype-driven clustering. Standard Seurat data processing and normalization steps were performed: SCTransform, RunPCA, RunUMAP, FindNeighbors and FindCluster. The final clustering resolution was 0.6. Analysis of spike-specific CD8 T cells in LC was restricted to 7 samples from the Spike<sup>Act</sup> LC subset, as indicated in Fig. 4A. To minimize the impact of highly expanded responses in some samples, all antigen-specific populations were downsampled to 150 cells per sample prior to differential gene expression and gene set enrichment analyses. DEGs were calculated using the functions FindAllMarkers or FindMarkers for pairwise comparisons using a log2 fold-change threshold of 0.3 and an adjusted P value of less than 0.05. Venn diagrams and upset plots were generated using the eulerr and UpSetR packages. Module scores were calculated for published and the newly generated acute and bystander gene sets using the AddModuleScore\_UCell function (UCell). Gene set enrichment analysis was performed using the GSEA function from clusterProfiler and plotted with gseaplot2 (enrichplot). Average gene expression was calculated using the AggregateExpression function (Seurat). Heat maps were generated using complexheatmap (version 2.20.0). Gene sets for gene expression programs (GEPs) from T-Cell AnnoTator (TCAT) were derived using all genes from the published gene sets, excluding CITE-

seq markers(64). Analysis of T cell clones was performed using ScRepertoire package, with clones defined using the CTaa parameter.

**CD4 T cell dataset:** For the CD4 T cell dataset, data were analyzed in R (version 4.4.3) using Seurat (version 5.4.0). Cells with less than 1,200 or more than 20,000 reads (nCount\_RNA), cells with more than 3800 detected genes (nFeature\_RNA), and cells with more than 4% of reads mapping to mitochondrial genes (percent\_mt) were excluded. TCR genes as well as HLA-G and AC004448.2 were excluded to avoid clonotype- or donor-driven clustering. In addition, a cluster expressing myeloid markers (e.g., CD14, LYZ, FCN1) and a low-quality cluster with fewer detected genes and higher mitochondrial transcript content than the remaining clusters were removed. Standard Seurat data processing and normalization steps were performed on all CD4 T cells: SCTransform, RunPCA, FindNeighbors, and RunUMAP. A set of immediate early genes (IEGs) and sex-chromosomal genes was excluded from variable features to minimize confounding effects caused by sample handling, sample origin from different cohorts, and unequal sex distribution. Clustering was performed using Seurat's FindClusters function, the final resolution used for downstream analyses and visualizations was 0.4.

DEGs were calculated on a subsampled object containing a maximum of 250 spike-specific CD4 T cells per sample using the FindMarkers function combined with the model-based analysis of single-cell transcriptomics (MAST) framework. Sex was included as a latent variable. Genes with an absolute log2FC > 0.25 and an adjusted p-value < 0.05 were considered statistically significant. Heat maps were generated using complexheatmap (version 2.20.0), and Venn diagrams were generated using the eulerr package (version 7.0.4). Analysis of CD4 T cell clonality was performed with scRepertoire (version 2.0.7). Clones were defined using the CTaa parameter.

#### **Stimulation assay for cytokine production by spike-specific CD8 T cells**

PBMC samples were thawed by warming frozen cryovials in a 37°C water bath and resuspending cells in 10 mL R10. Cells were washed once in R10, counted using a Countess automated cell counter (ThermoFisher), and resuspended in fresh R10 to a density of $10 \times 10^6$ cells/mL. For each condition, $2 \times 10^6$ cells were plated in 200 μL in 96-well round-bottom plates and rested overnight in a humidified incubator at 37°C, 5% CO<sub>2</sub>. After 16 hours, cells were stimulated for 6 hours with costimulation (anti-human CD28/CD49d, BD Biosciences) and custom peptide pools containing all spike peptides used in HLA-I tetramers (see **Table S4**, "Used in Fig. 6" column), with each peptide at a final concentration of 1 μg/mL. Matched unstimulated samples for each donor at each timepoint were treated with costimulation alone. After 2 hours, monensin (GolgiStop, BD Biosciences) was added to all cultures for the final 4 hours, after which cells were washed in PBS supplemented with 2% FBS (FACS buffer), stained for 10 min at RT with Ghost Viability Dye Violet 510 (Tonbo, 1:1000 in PBS) and Fc receptor blocking solution (Human TruStain FcX™, BioLegend, 1:1000 in PBS), and washed once in FACS buffer. Surface staining for 30 min at RT was then performed with a cocktail of surface staining antibodies in a 1:1 solution of FACS and brilliant stain buffer (BD). Cells were washed once in FACS buffer and fixed and permeabilized for 30 min at RT (eBioscience Foxp3/Transcription Factor Staining Kit). Cells were washed once in 1X Permeabilization Buffer prior to staining with antibodies directed against TNF, IFN-γ, IL-2, and other intracellular targets. Following incubation with antibodies overnight at 4°C, cells were washed once in 1X Permeabilization buffer and resuspended in FACS buffer prior to data acquisition.

#### **CellTrace Violet T cell proliferation assay**

Cryopreserved PBMCs were thawed as for the cytokine production assay described above, counted, and stained with CellTrace Violet (CTV) according to the manufacturer's protocol. Briefly, $3 \times 10^6$ PBMCs from each sample were resuspended in 1 mL of warm PBS containing 0.5 μM CTV and incubated in a 37°C water bath for 20 min. The staining reaction was quenched by adding 10 mL of R10 medium and incubating for additional 15 min in a 37°C water bath. Cells were then pelleted at 800xg and resuspended in 2 mL of RPMI supplemented with 10% human serum, 2 mM L-Glutamine, 100 U/mL Penicillin, and 100 μg/mL Streptomycin (Human Serum R10). These cells were split between two wells of a 24-well plate that each contained $1.5 \times 10^6$ CTV-labelled PBMCs in 1 mL of Human Serum R10. Cells were rested overnight in a humidified incubator at 37°C, 5% CO<sub>2</sub>. After 16 hours, cells were stimulated for 96 hours with costimulation (anti-human CD28/CD49d, BD Biosciences) and the same custom spike peptide pool used for the cytokine production assays at a final concentration of 1 μg/mL for each peptide. Paired unstimulated samples for each donor at each timepoint were treated with costimulation alone. No additional cytokines (e.g. IL-2) were added to the cultures. After 96 hours, cells were harvested, stained for 10 min at RT with Zombie Near-Infrared Viability Dye and Fc receptor blocking solution (Human TruStain FcX™, BioLegend) and washed once in FACS buffer. Surface staining for 30 minutes at RT was then performed with a cocktail of antibodies targeting surface antigens in a 1:1 solution of FACS and brilliant stain buffer (BD). After surface staining, cells were washed and fixed in 1X Stabilizing Fixative (BD Biosciences) overnight at 4°C.

#### **Code availability**

All analyses were done with custom R scripts using standard R packages and are available upon request. No new algorithms were developed for this study.

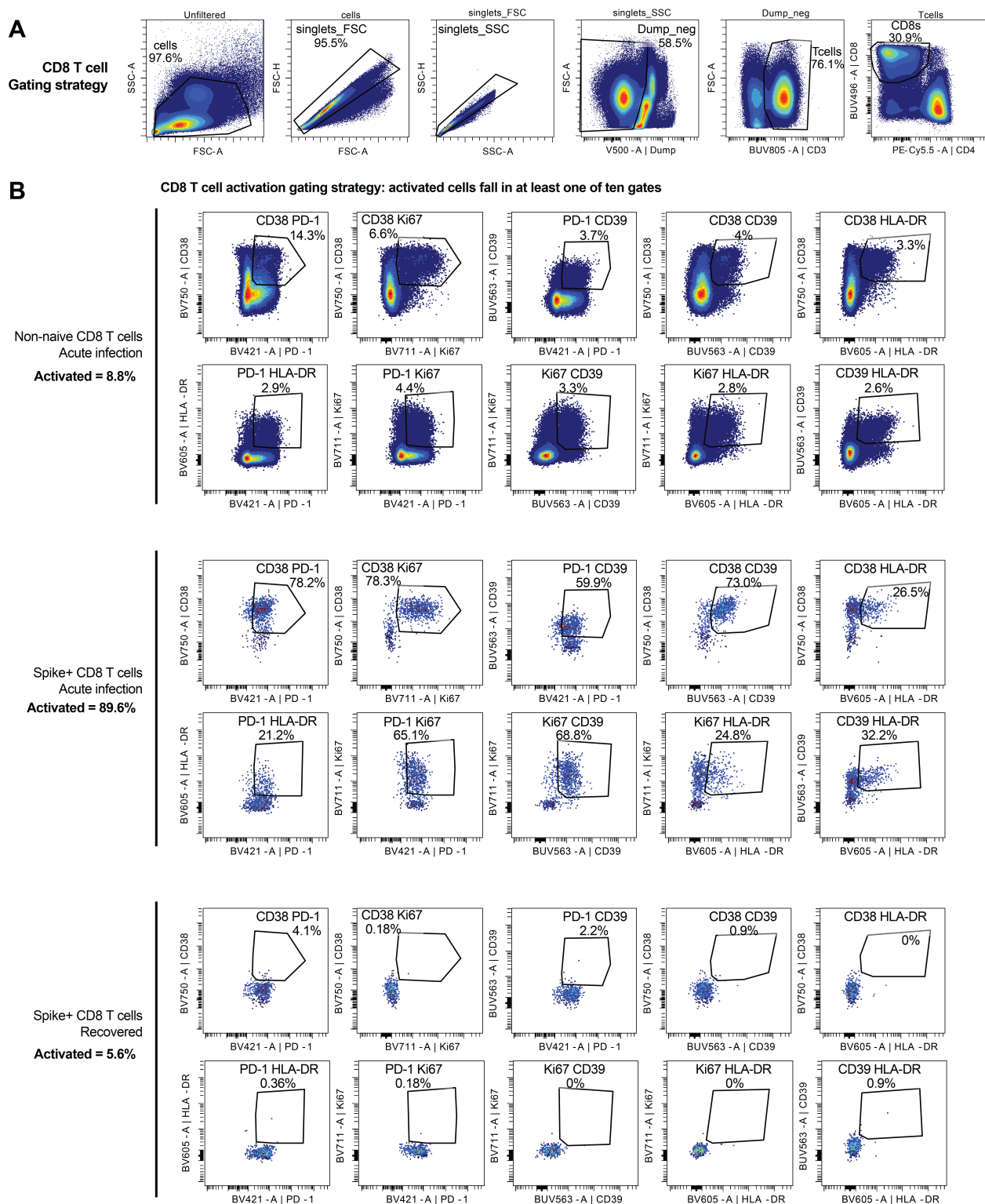

**Fig. S1: CD8 T cell gating strategy**

(A) Representative flow cytometry plots depicting the gating strategy used to identify CD8 T cells. (B) Representative flow cytometry plots depicting the gating strategy used to identify activated CD8 T cells. Cells were assigned as activated if they were found in at least one of the 10 gates, indicating co-expression of at least 2 activation markers.

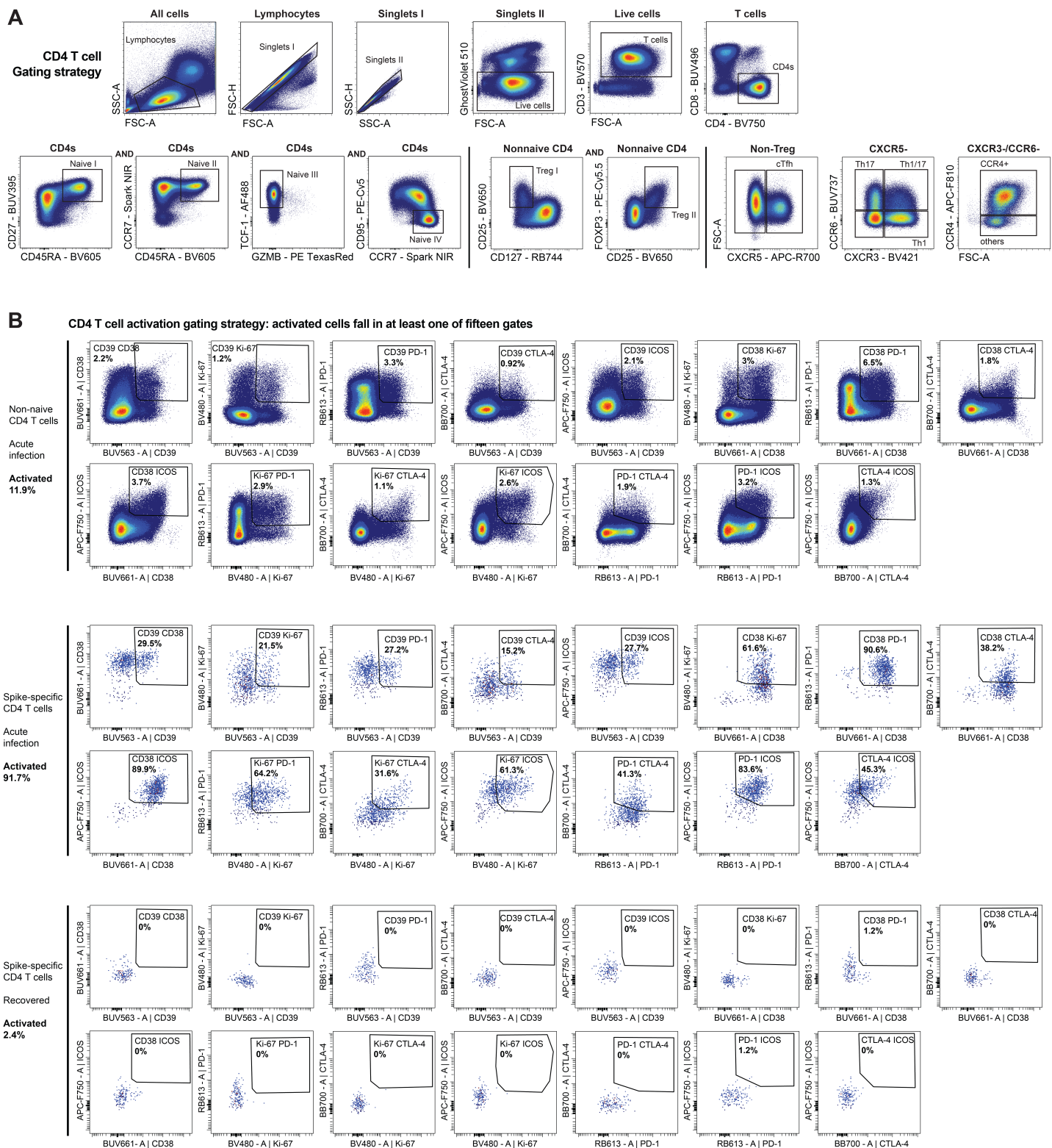

**Fig. S2: CD4 T cell gating strategy**

(A) Representative flow cytometry plots depicting the gating strategy used to identify nonnaive CD4 T cells and to define distinct T helper polarization phenotypes in flow cytometric samples stained with HLA class II tetramers. (B) Representative flow cytometry plots depicting the gating strategy used to identify activated CD4 T cells. Cells were assigned as activated if they were found in at least one of the 15 gates, indicating co-expression of at least 2 activation markers.

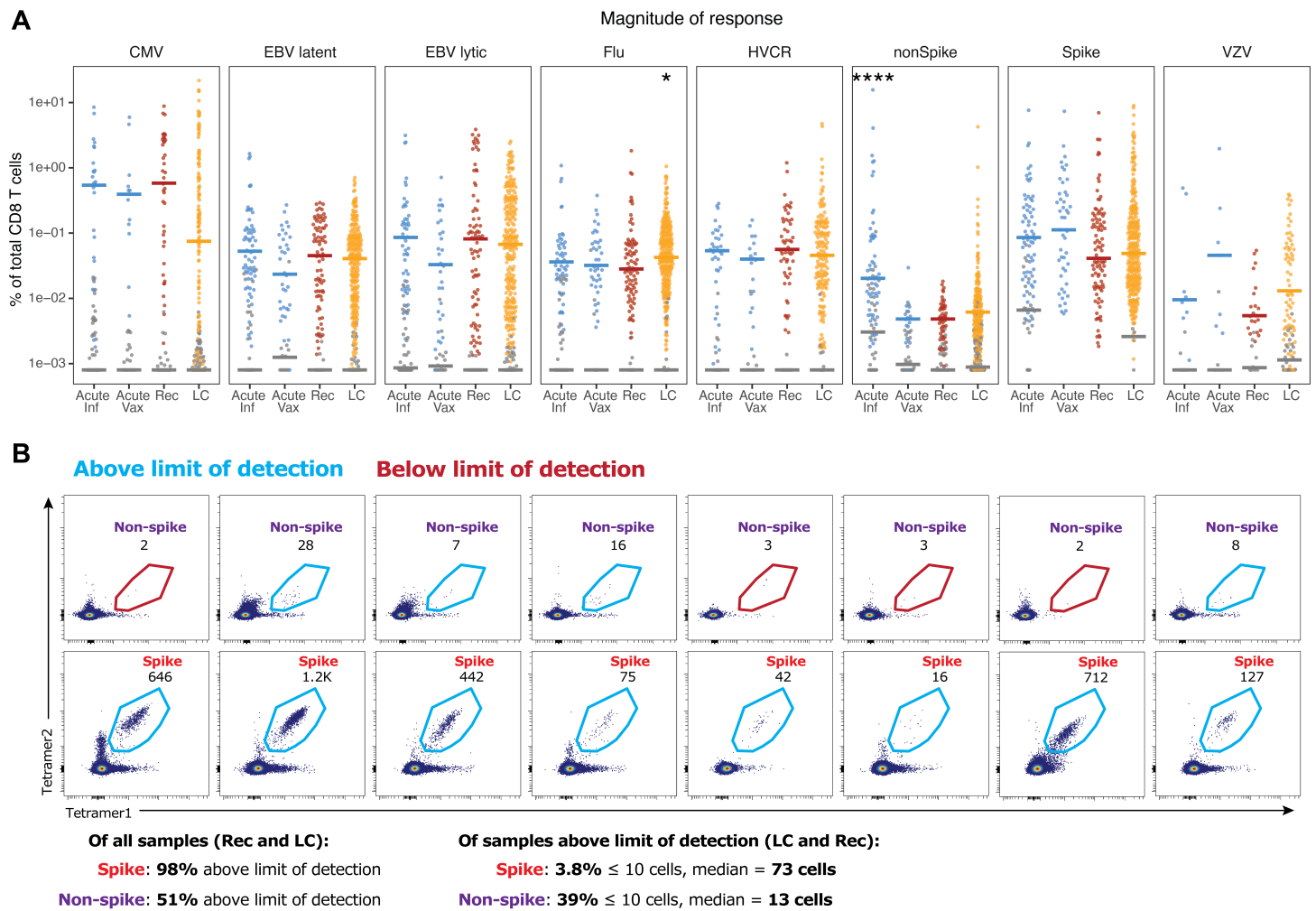

**Fig. S3: Detection of antigen-specific CD8 T cells with HLA-I tetramers**

(A) Summary data depicting the percent of total CD8 T cells staining with HLA-I tetramers loaded with viral peptides. Each point represents an individual sample. Grey points were below the limit of detection, colored points were included in the analyses throughout the manuscript. Statistical significance was determined using Wilcoxon tests with Benjamini-Hochberg correction for multiple comparisons. All statistical tests are comparisons to recovered controls. \*$p < 0.05$, \*\*$p < 0.01$, \*\*\*$p < 0.001$, \*\*\*\*$p < 0.0001$. (B) Paired representative flow cytometry plots from 8 samples depicting CD8 T cells specific for non-spike (top) or spike (bottom) epitopes from SARS-CoV-2 as detected using dual-fluorophore tetramer staining. Numbers indicate the count of tetramer+ cells. CD8 T cells targeting the non-spike epitopes restricted to HLA-A\*02:01 and HLA-A\*03:01 used in this study were rare and frequently below the limit of detection (minimum 5 tetramer+ cells), precluding robust analysis of these responses.

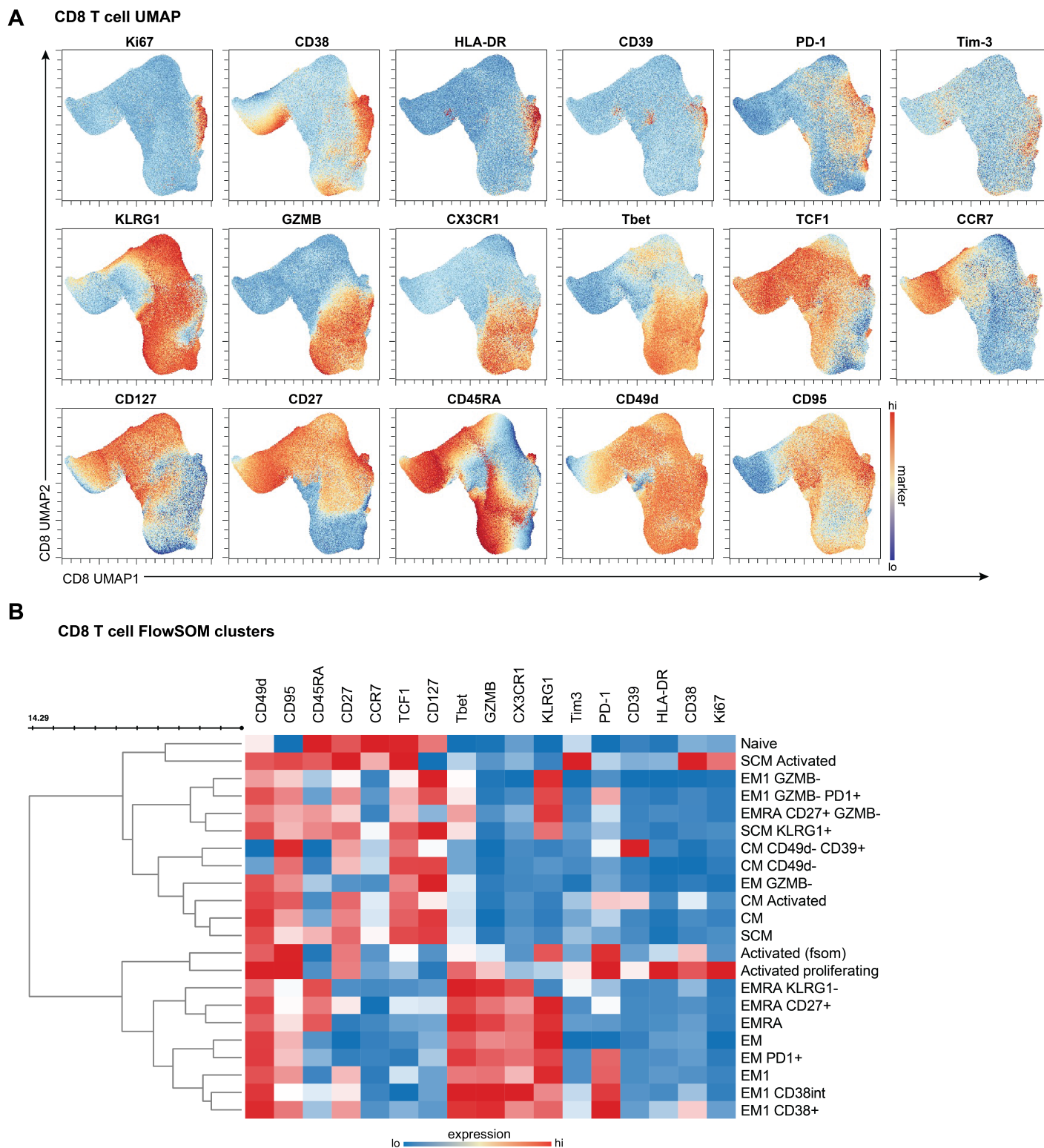

**Fig. S4: Dimensionality reduction and clustering of CD8 T cells**

(A) UMAP representations of all CD8 T cells from all samples, colored by the intensity of expression of the indicated markers across UMAP space. (B) Clustered heatmap depicting the average expression of each marker for each of the indicated FlowSOM metaclusters across bulk and virus-specific CD8 T cells.

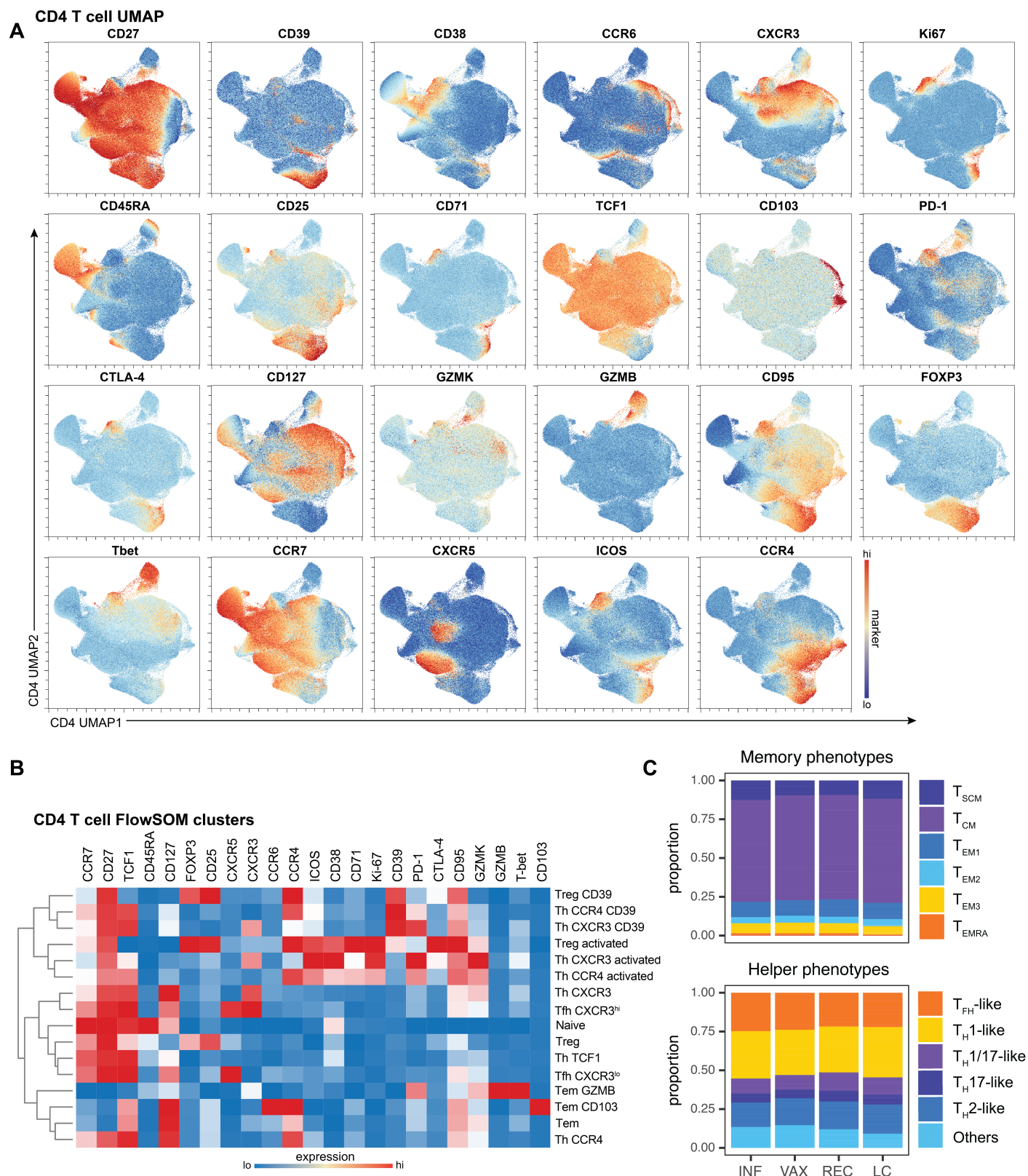

**Fig. S5: Dimensionality reduction and clustering of CD4 T cells**

(A) UMAP representations of all CD4 T cells from all samples, colored by the intensity of expression of the indicated markers across UMAP space. (B) Clustered heatmap depicting the average expression of each marker for each of the indicated FlowSOM metaclusters across bulk and spike-specific CD4 T cells. (C) Summary data comparing the distribution of bulk CD4 T cells across manually-gated memory (top) and helper (bottom) CD4 T cell subsets, showing minimal differences across groups.

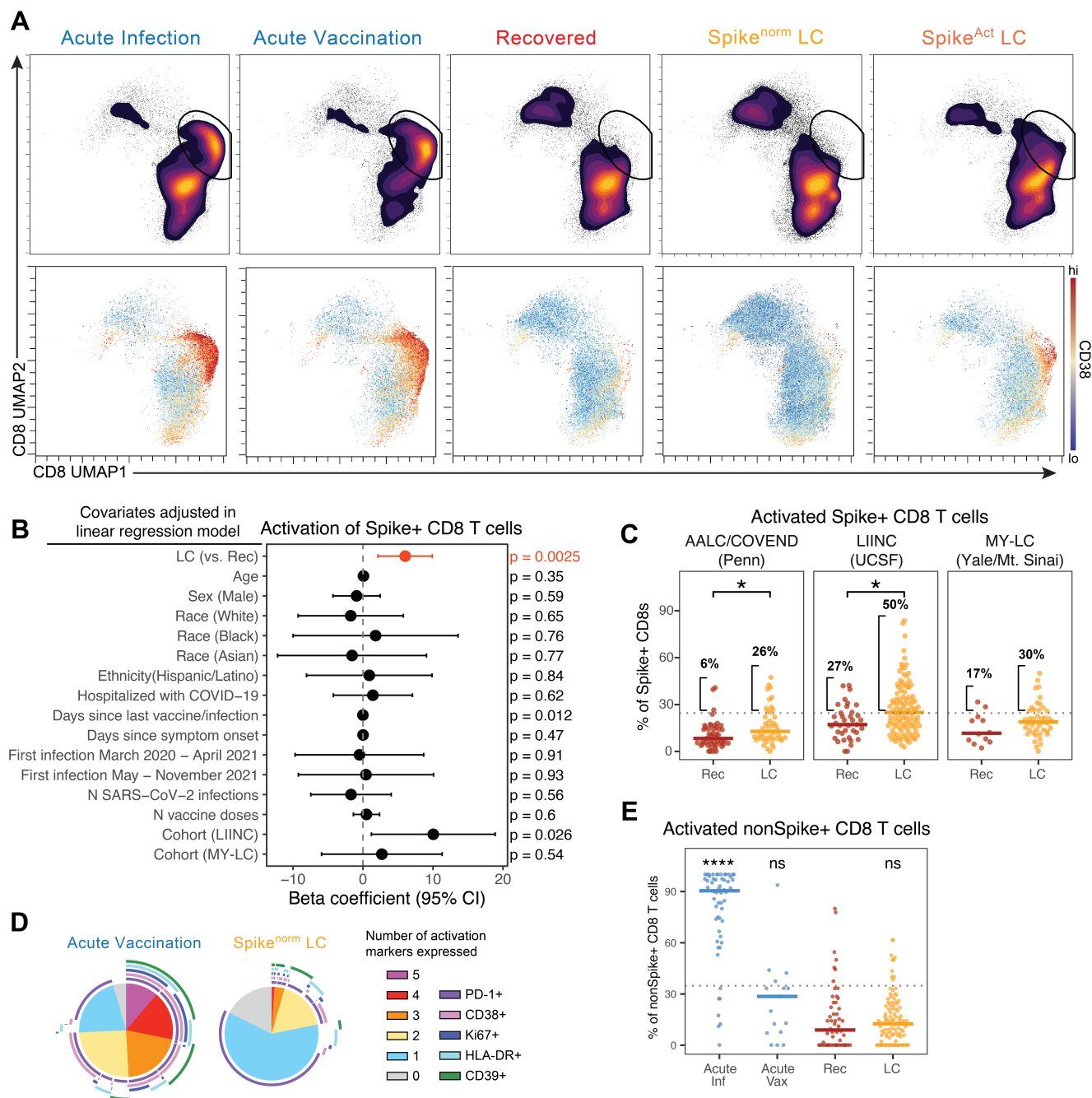

**Fig. S6: Spike-specific CD8 T cells are activated in a subset of people with LC.**

(A) UMAP representations of all spike-specific CD8 T cells from the indicated groups depicting the density (top) and CD38 expression (bottom) of spike-specific CD8 T cells in UMAP space. (B) Forest plot depicting the beta coefficient and 95% confidence interval from a linear regression model testing the association between LC and spike-specific CD8 T cell activation after simultaneously adjusting for each of the indicated covariates. Orange represents the effect of LC compared to recovered controls, and black represents the effect of the indicated covariate on spike-specific CD8 T cell activation. Values to the right of the dashed line indicate a positive association with elevated activation of spike-specific CD8 T cells. Reference levels for categorical covariates were: Sex – Female; Race – Other; Cohort – Penn/AALC. (C) Summary data of the activation of spike-specific CD8 T cells comparing recovered controls to LC samples from each of the 3 cohorts analyzed in this study. Horizontal dotted line represents the mean plus one standard deviation of recovered controls across all cohorts, and the percent of samples falling above this threshold is indicated. (D) SPICE analysis of the co-expression of five activation markers by spike-specific CD8 T cells from the indicated groups. (E) Summary data of the activation of non-spike SARS-CoV-2-specific CD8 T cells by group. Horizontal dotted line represents the mean plus one standard deviation of recovered controls. Statistical significance was determined using Wilcoxon tests with Benjamini-Hochberg correction for multiple comparisons. \*$p < 0.05$, \*\*$p < 0.01$, \*\*\*$p < 0.001$, \*\*\*\*$p < 0.0001$. In the absence of brackets, all statistical tests are comparisons to recovered controls.

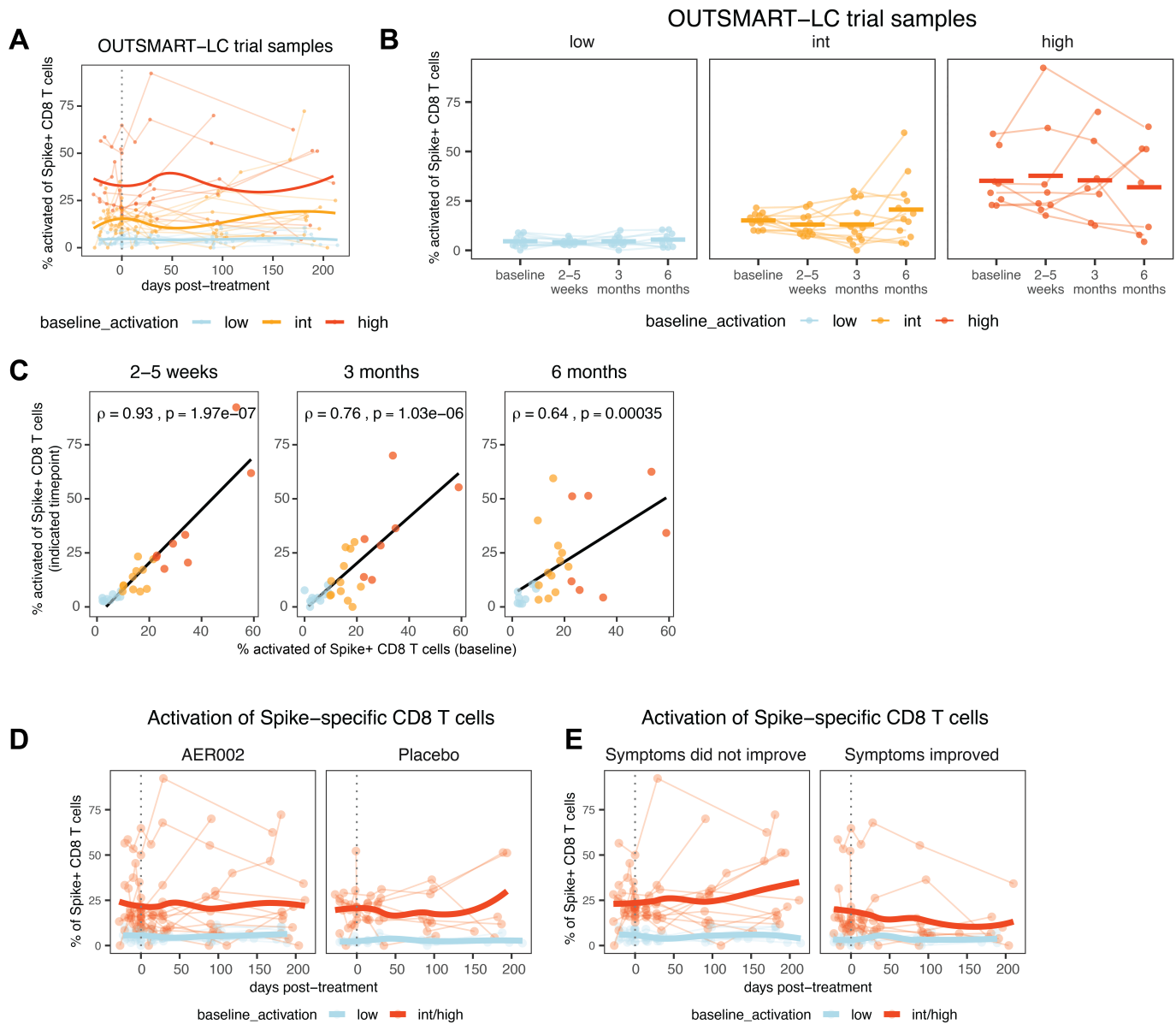

**Fig. S7: Spike-specific CD8 T cell activation was maintained over time in the OUTSMART-LC clinical trial and was not associated with treatment or symptom improvement**

(A) Longitudinal analysis of spike-specific CD8 T cell activation of all individual samples from the OUTSMART-LC clinical trial. (B) Longitudinal analysis of spike-specific CD8 T cell activation of individuals from the OUTSMART-LC clinical trial, separated by baseline activation levels and averaging activation of all samples falling within the indicated timepoint windows relative to treatment initiation. (C) Spearman rank correlation of average activation in baseline samples (x-axis) with average activation at later visits after treatment initiation. (D-E) Longitudinal analysis of spike-specific CD8 T cell activation of all individual samples from the OUTSMART-LC clinical trial, separated by (D) treatment group or (E) whether participants reported symptom improvement. Samples were divided into low ($x < 9.5\%$), intermediate ($9.5\% < x < 22\%$), or high ($x > 22\%$) average activation at baseline timepoints. Lines connect individuals sampled longitudinally.

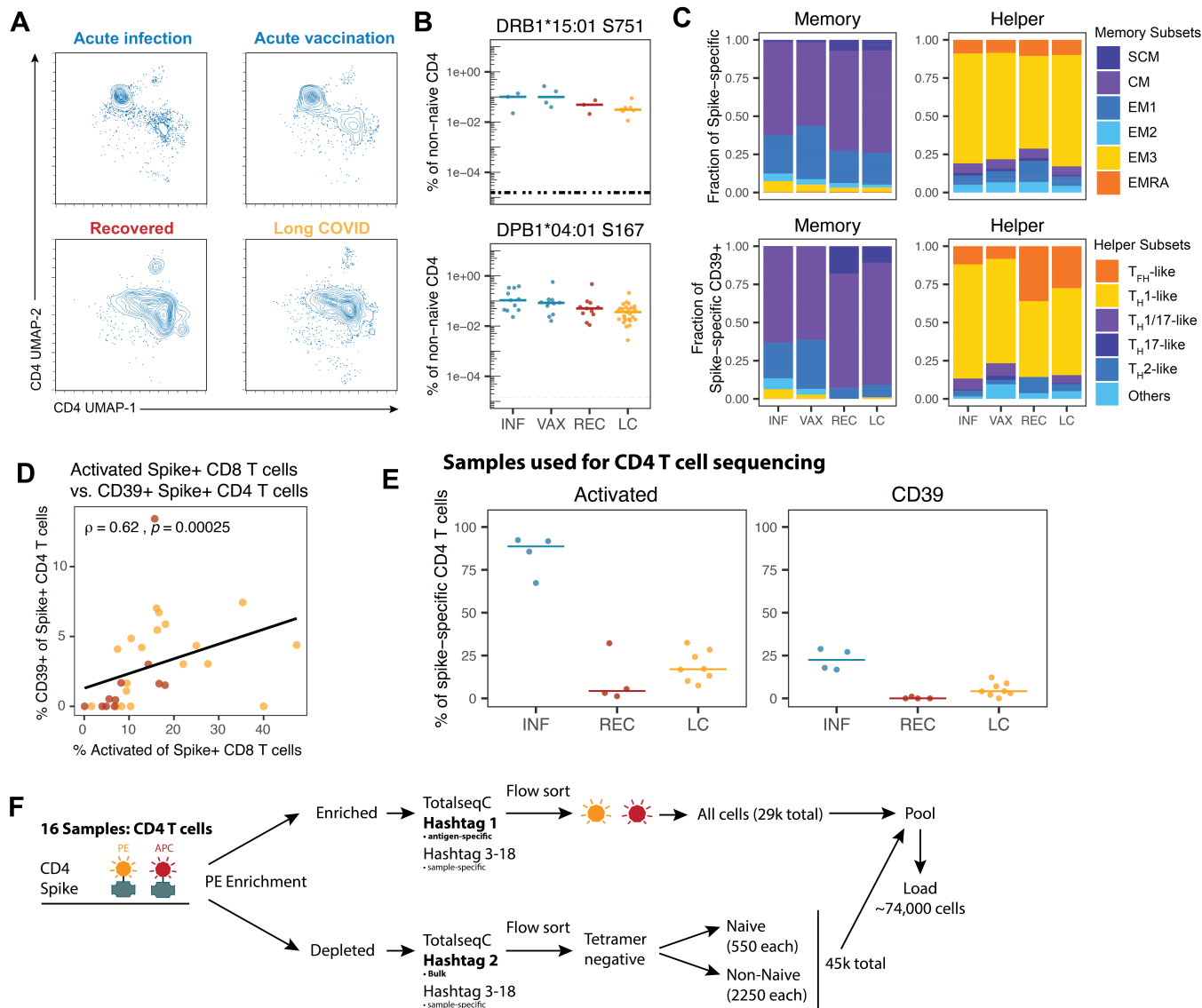

**Fig. S8: Spike-specific CD4 T cell supplementary data and single-cell sequencing workflow**

(A) Plots displaying the position of spike-specific CD4 T cells in the UMAP space introduced in Fig. 1 and Fig. S5. (B) Dot plot showing the frequency of CD4 T cells specific for the Spike S751 or S167 epitope across acute infection, acute vaccination, LC, and recovered controls. (C) Stacked bar plots displaying the memory and helper phenotype polarization of spike-specific CD4 T cells (upper panels) and spike-specific CD39+ CD4 T cells (lower panels) across acute infection, acute vaccination, LC, and recovered controls. (D) Correlation between spike-specific CD39+ CD4 T cells and activated spike-specific CD8 T cells in donors that were eligible for CD4 as well as CD8 T cell analysis. (E) Summary data showing activation status and CD39 expression, determined by flow cytometry, of spike-specific CD4 T cells in samples used for 10x single-cell RNA sequencing. (F) Schematic of the workflow for the preparation of bulk and spike-specific CD4 T cells for single-cell RNA sequencing.

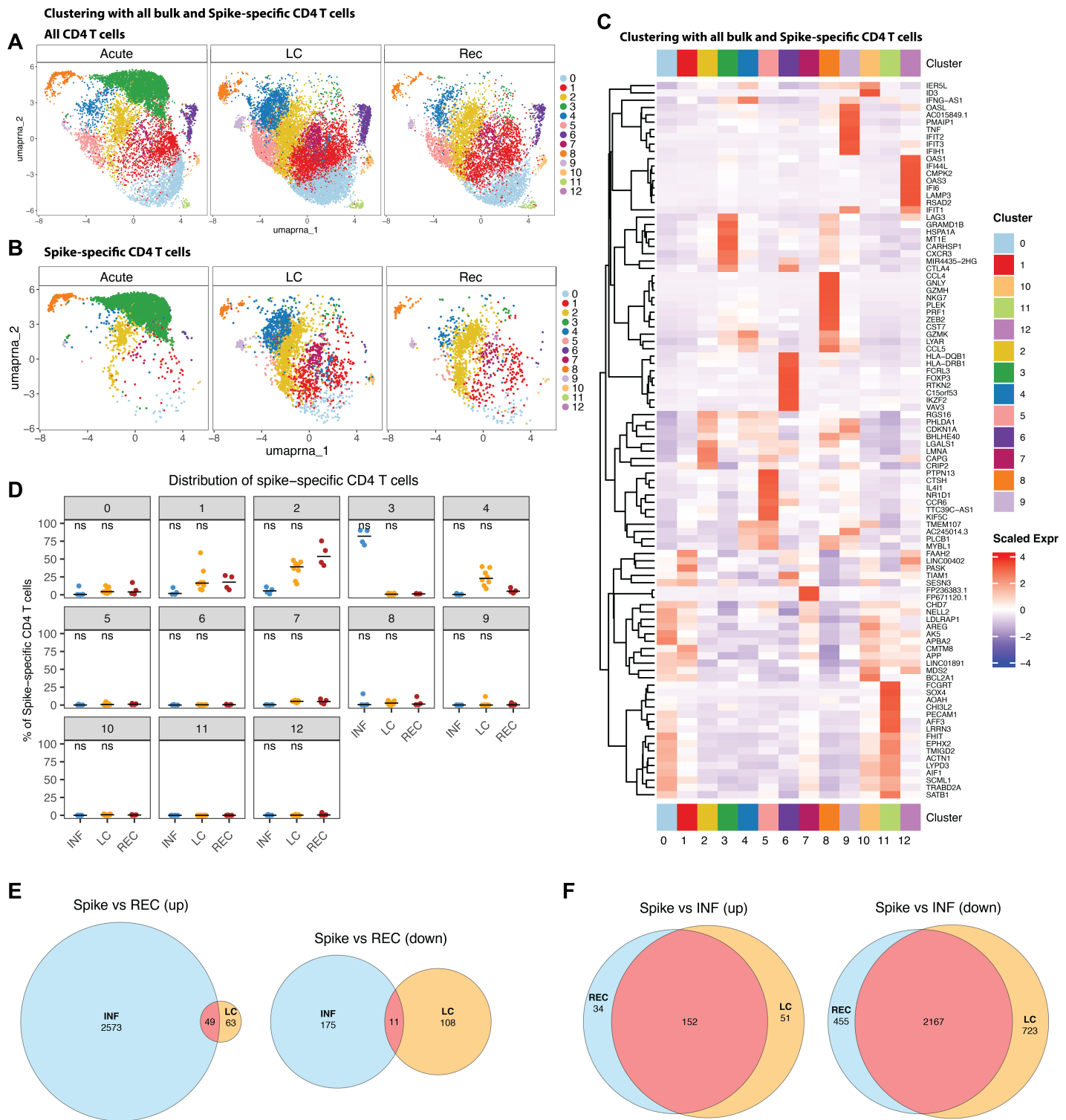

**Fig. S9: Single-cell RNA-seq of bulk and spike-specific CD4 T cells**

(A-B) UMAP representation of the clustering of (A) all or (B) spike-specific CD4 T cells from donors with acute infection, LC, and recovered controls. (C) Heatmap displaying the top differentially expressed genes of the CD4 T cell clusters introduced in Fig. 2K and Fig. S9. (D) Percentage of spike-specific CD4 T cells from each sample in clusters 0-12. (E-F) Venn diagrams showing the overlap of upregulated and downregulated DEGs in spike-specific CD4 T cells from (E) acute infection or LC, each compared to recovered controls (Spike vs REC), and (F) recovered controls or LC, each compared to acute infection (Spike vs INF). Significant DEGs were identified via the Seurat FindMarkers function by applying the model-based analysis of single-cell transcriptomics (MAST)

framework and considering sex as a latent variable. Statistical significance was determined using Wilcoxon tests with Benjamini-Hochberg correction for multiple comparisons. \*$p < 0.05$, \*\*$p < 0.01$, \*\*\*$p < 0.001$, \*\*\*\*$p < 0.0001$.

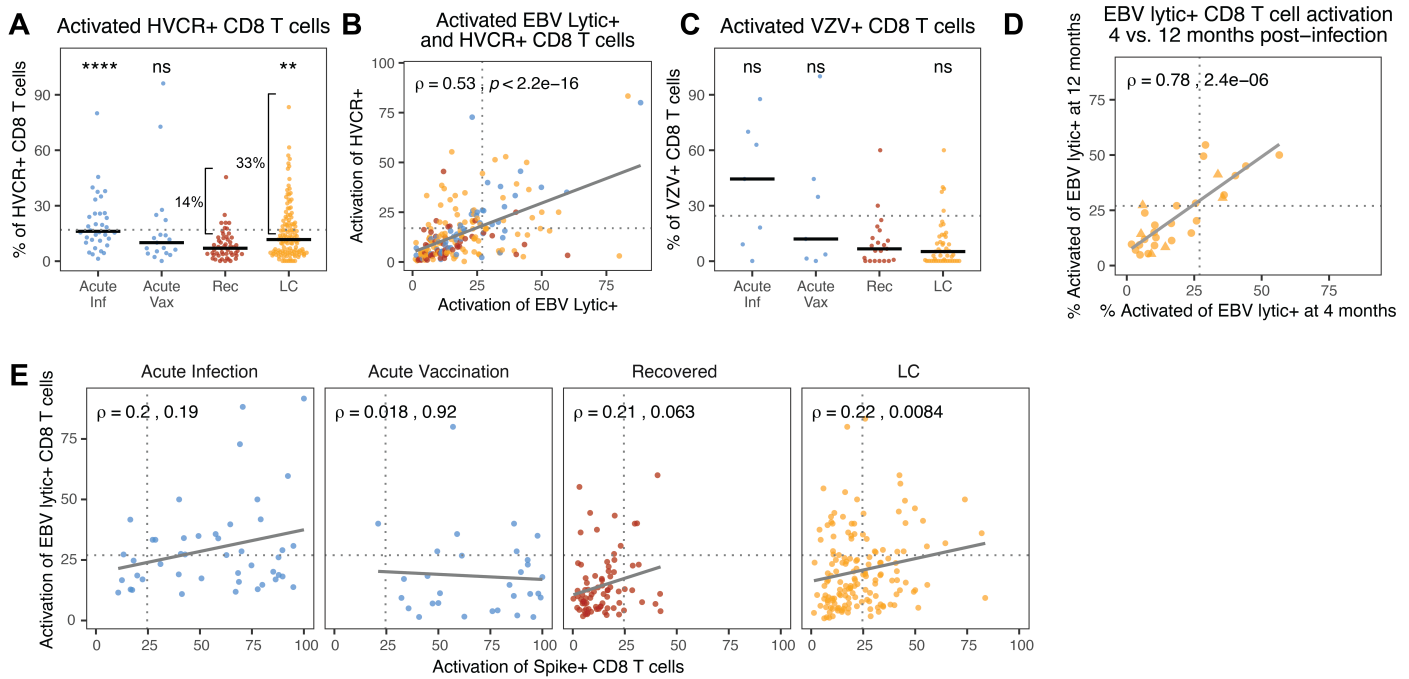

**Fig. S10: Activation of herpesvirus-specific CD8 T cells in LC.**

(A) Summary data of the activation (defined in Fig. 1B and Fig. S1) of herpesvirus cross-reactive (HVCR)-specific CD8 T cells by group. Horizontal dotted line represents the mean plus one standard deviation of recovered controls. (B) Correlation of activation of EBV lytic- and HVCR-specific CD8 T cells from all samples. Points represent individual samples where both responses were detectable and are colored by group (blue = acute infection or vaccination, red = recovered, orange = LC). (C) Summary data of the activation of VZV-specific CD8 T cells by group. Horizontal dotted line represents the mean plus one standard deviation of recovered controls. (D) Correlation of the percent activation of EBV lytic-specific CD8 T cells from paired longitudinal LC samples from the LIINC cohort collected at 4 and 12 months post-infection. (E) Correlation of the percent activation of EBV lytic-specific and spike-specific CD8 T cells from samples split by group. All correlations were calculated using nonparametric Spearman rank correlation. Statistical significance was determined using Wilcoxon tests with Benjamini-Hochberg correction for multiple comparisons. \*$p < 0.05$, \*\*$p < 0.01$, \*\*\*$p < 0.001$, \*\*\*\*$p < 0.0001$. In the absence of brackets, all statistical tests are comparisons to recovered controls.

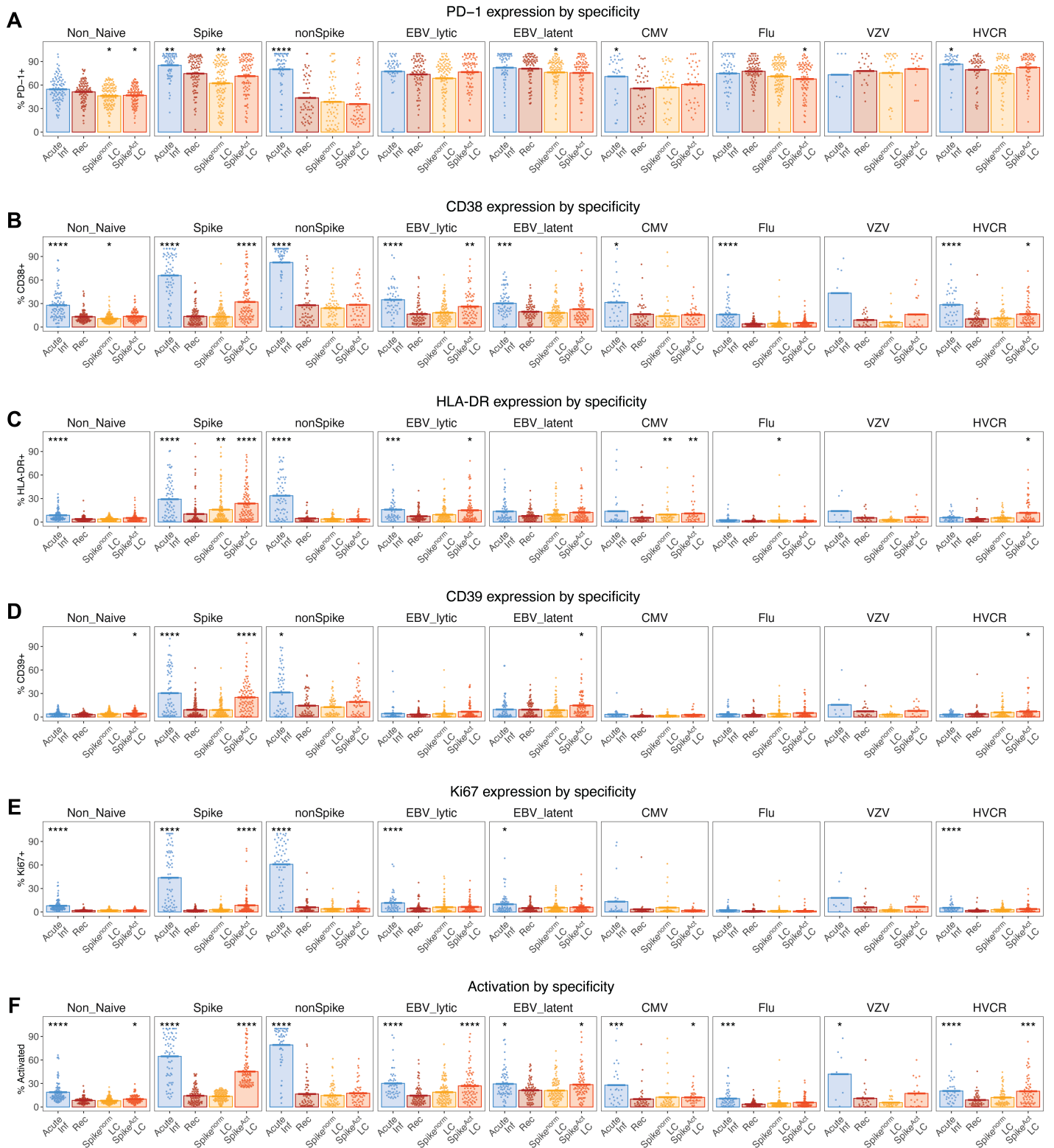

**Fig. S11: Expression of individual activation markers by all CD8 T cell specificities.**

(A-F) Summary data of the expression of each individual activation marker by the indicated antigen-specific CD8 T cell populations, including (A) PD-1, (B) CD38, (C) HLA-DR, (D) CD39, (E) Ki67, and (F) overall activation as measured by expressing at least 2 of 5 markers. LC samples were split into Spike<sup>Act</sup> LC and Spike<sup>norm</sup> LC subsets using the cutoff in Fig. 2B. Statistical significance was determined using Wilcoxon tests with Benjamini-Hochberg correction for multiple comparisons. All statistical tests are comparisons to recovered controls. \*$p < 0.05$, \*\*$p < 0.01$, \*\*\*$p < 0.001$, \*\*\*\*$p < 0.0001$.

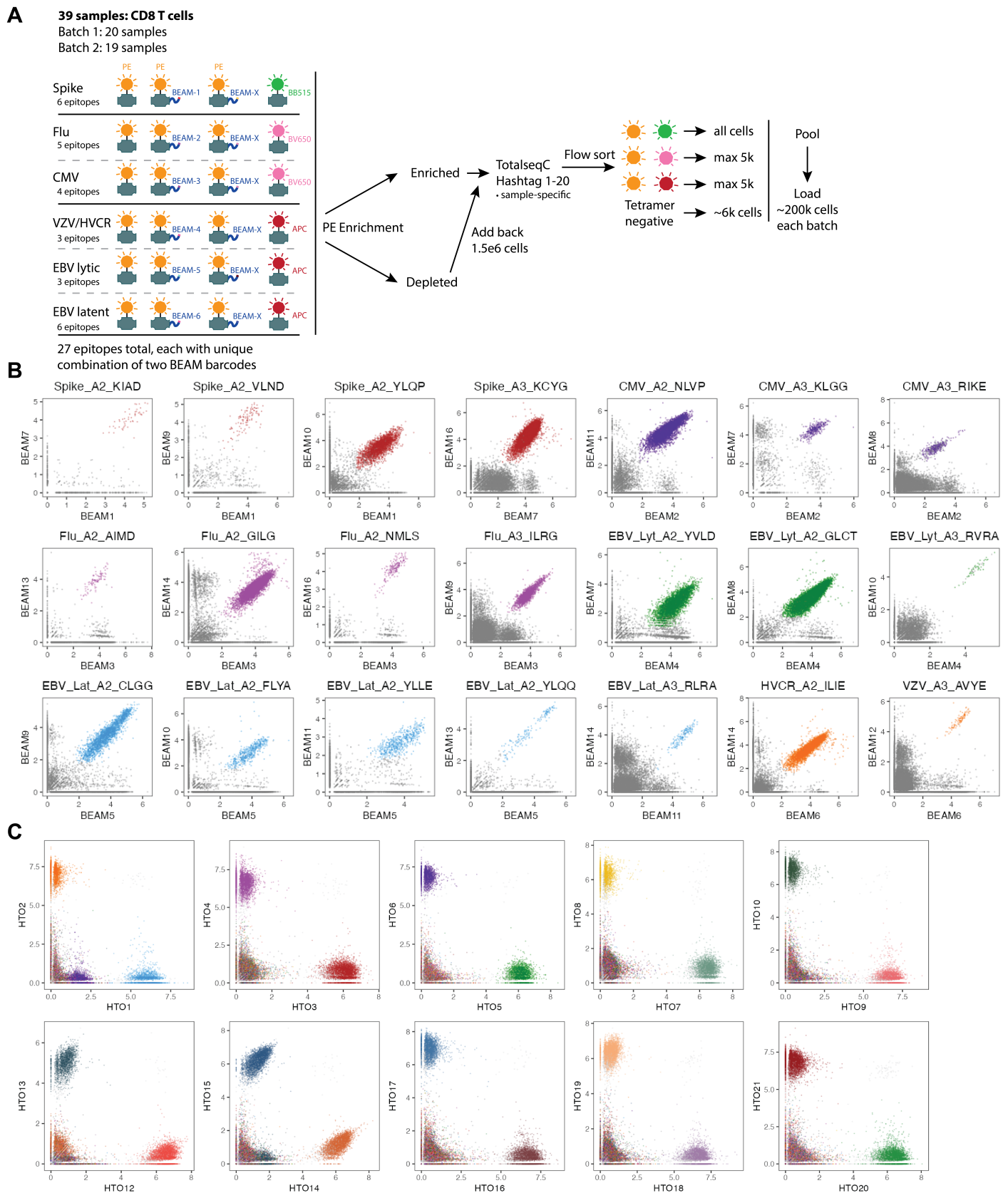

**Fig. S12: Overview of barcoded tetramer staining for antigen-specific CD8 T cell single-cell sequencing.**

(A) Schematic representation of the workflow for tetramer staining and sorting CD8 T cells for single-cell RNA-, TCR-, and tetramer barcode sequencing. (B) Representative plots showing the specificity of the dual-code approach to identify tetramer-binding CD8 T cells of the indicated antigen specificities. Each point is a cell, colored by antigen specificity assignment. **(C)** Representative plots showing the specificity of sample hashtags used to distinguish cells from each of the 20 samples pooled in batch 1, with similar results for the 19 samples in batch 2. Each point is a cell, colored by sample hashtag assignment.

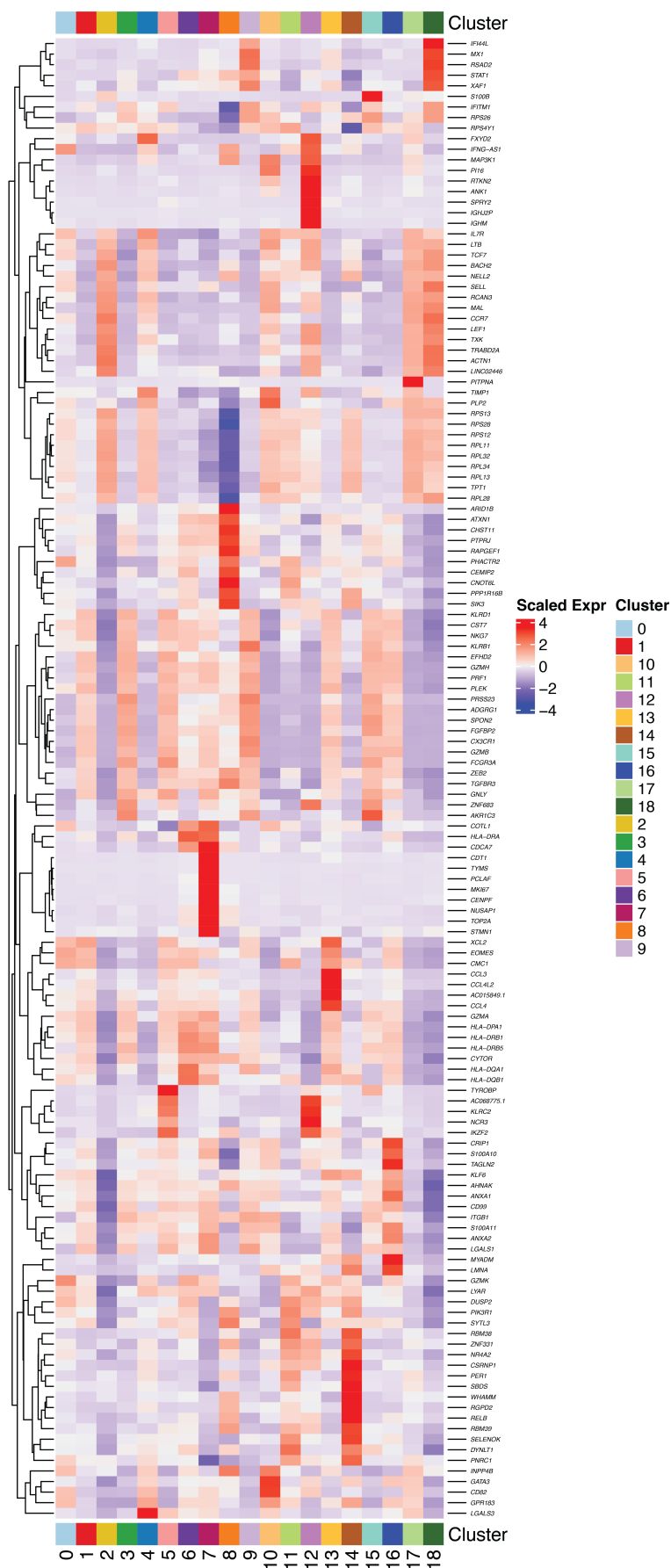

**Fig. S13: Differentially expressed genes defining clusters of CD8 T cells in gene expression data.**

Heatmap displaying the top differentially expressed genes of the CD8 T cell clusters introduced in Fig. 4B.

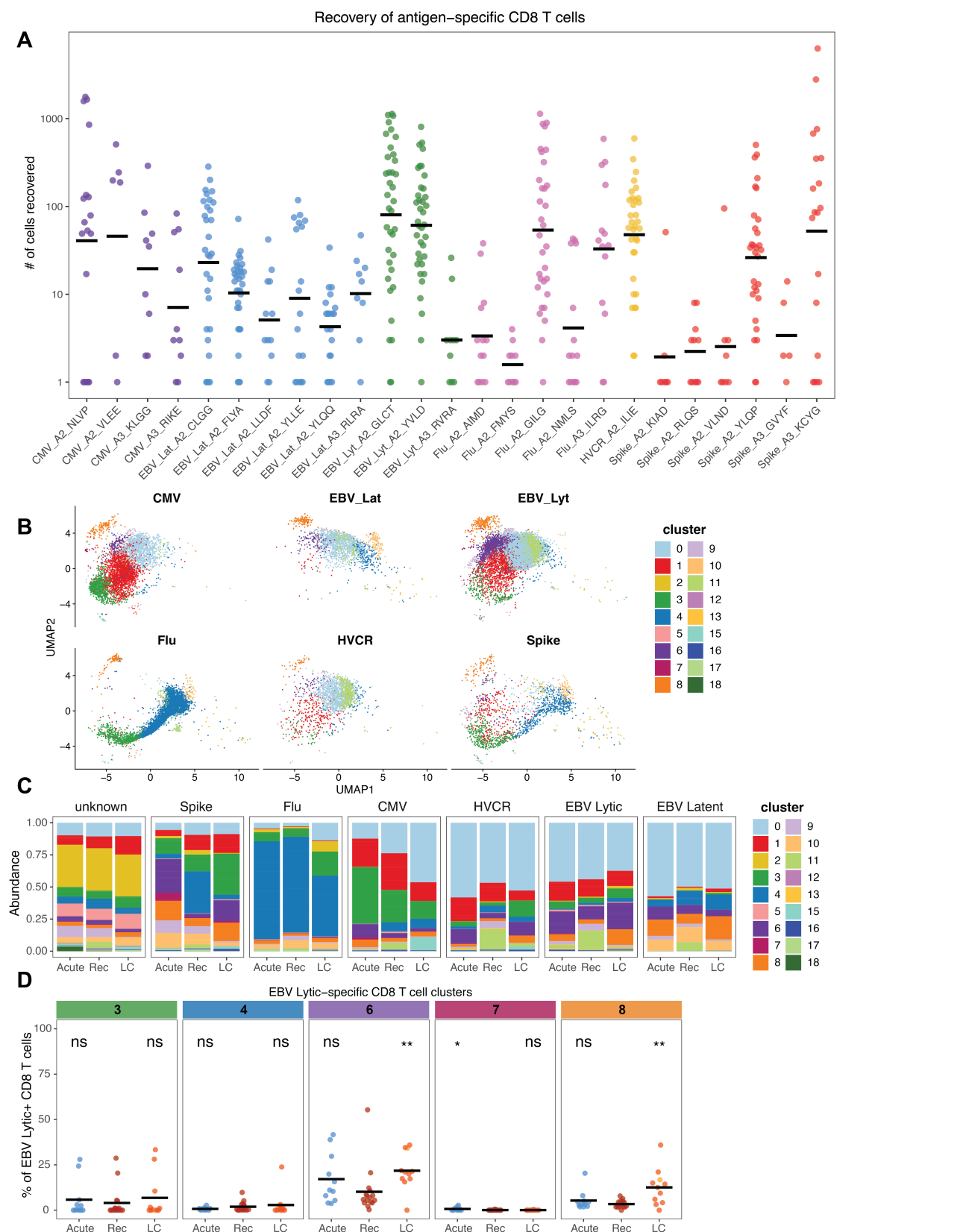

**Fig. S14: Recovery and clustering of antigen-specific CD8 T cells using barcoded HLA-I/peptide tetramers.**

**(A)** Summary data of the number of cells detected for each antigen specificity using tetramer barcodes. Each point is a sample, and samples are only plotted if at least one cell of the specificity in question was detected. **(B)** UMAP coordinates of antigen-specific CD8 T cell populations in recovered control samples. **(C)** Summary data of the percent of the indicated tetramer+ or bulk CD8 T cell populations found in each scRNA-seq cluster, split by group. **(D)** Quantification of the frequency of EBV lytic-specific CD8 T cells from each sample found in clusters of interest, split by group. Statistical significance was determined using Wilcoxon tests with Benjamini-Hochberg correction for multiple comparisons. All statistical tests are comparisons to recovered controls. \*$p < 0.05$, \*\*$p < 0.01$, \*\*\*$p < 0.001$, \*\*\*\*$p < 0.0001$.

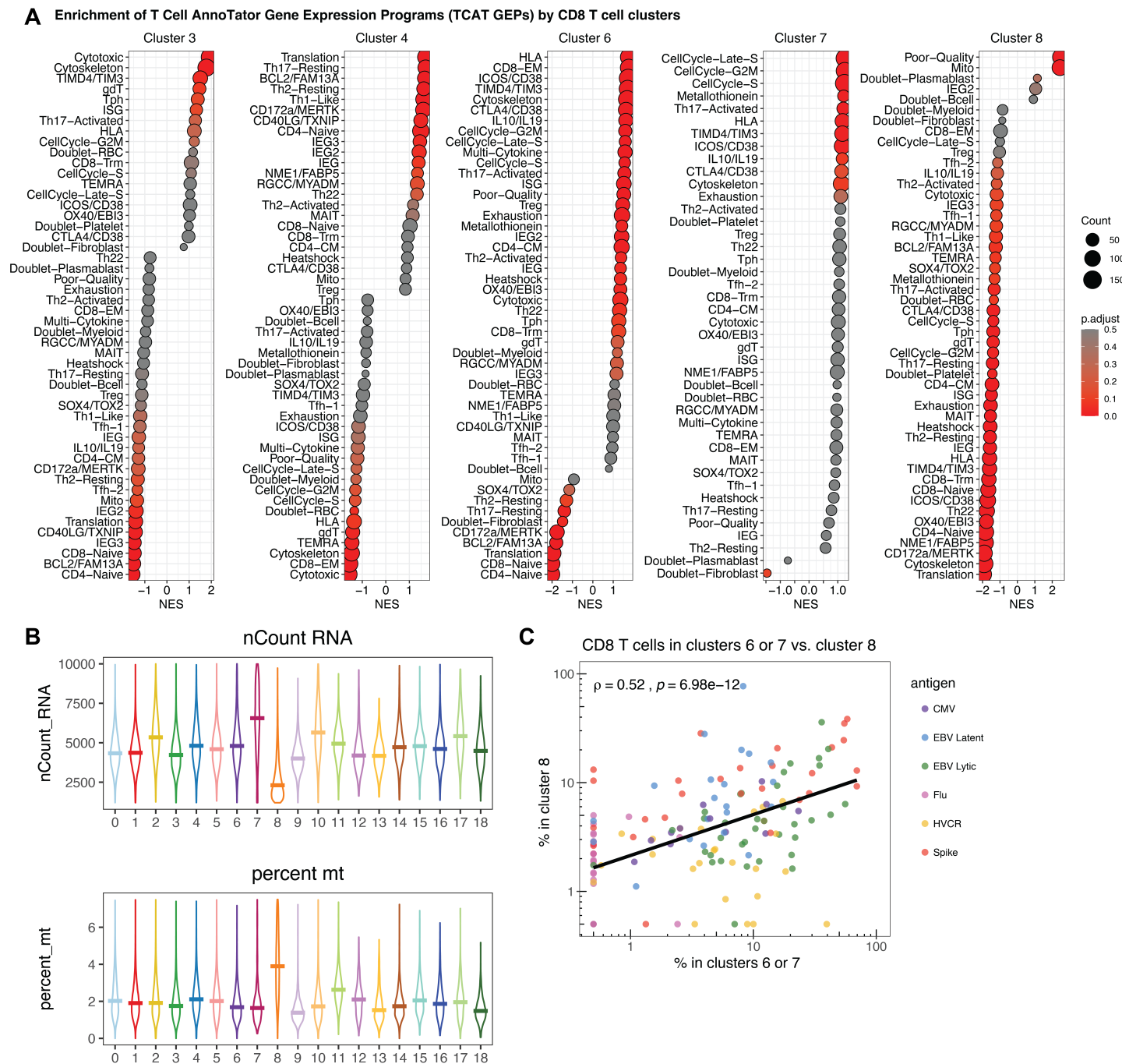

**Fig. S15: Gene expression programs that define clusters of interest and the identification of cluster 8 as an activation-associated cluster with low RNA content.**

(A) Gene set enrichment analyses using the gene lists for each gene expression program (GEP) from T-CellAnnoTator (TCAT)(64) to test for enrichment within ranked gene expression lists for clusters 3, 4, 6, 7, and 8 compared to all other cells in the dataset. Significantly enriched gene sets are colored red and have high NES values, significantly depleted gene sets are colored red and have low NES values. Gene sets in grey are not significantly enriched in that cluster. (B) Number of RNA features (top) and percent mitochondrial reads (bottom) detected for cells in each cluster. (C) Spearman rank correlation between the percent of cells in high-quality activated clusters 6 and 7 versus “low-quality” activated cluster 8 for each antigen specificity across all samples, colored by specificity.

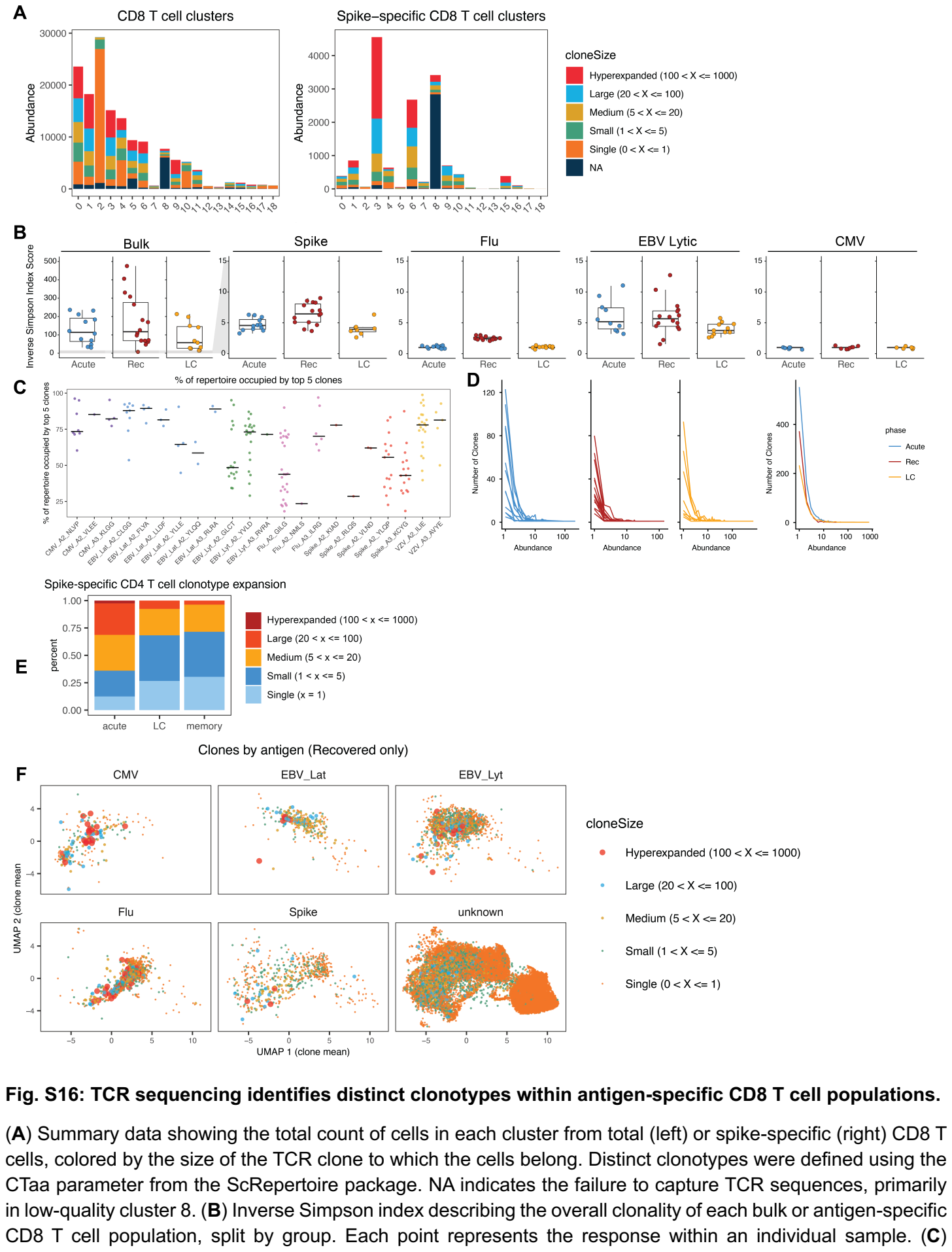

Summary data of the percent of the repertoire for each CD8 T cell response that was occupied by the top 5 clones, showing all responses where at least 5 clones were detected. Each point represents the response within an individual sample. **(D)** The number of spike-specific CD8 T cell clones detected at each clone size (abundance) for each sample (left) or summarized by group (right). **(E)** Summary data depicting the percent of the spike-specific CD4 T cell population belonging to different sizes of clonal expansions in samples from acute infection, LC, or recovered controls. **(F)** Mean UMAP coordinates for all cells within each CD8 T cell clonotype in recovered controls. Clones were defined using the CTaa parameter from the scRepertoire package, and the size and color of the points represent the number of cells recovered within each clonotype.

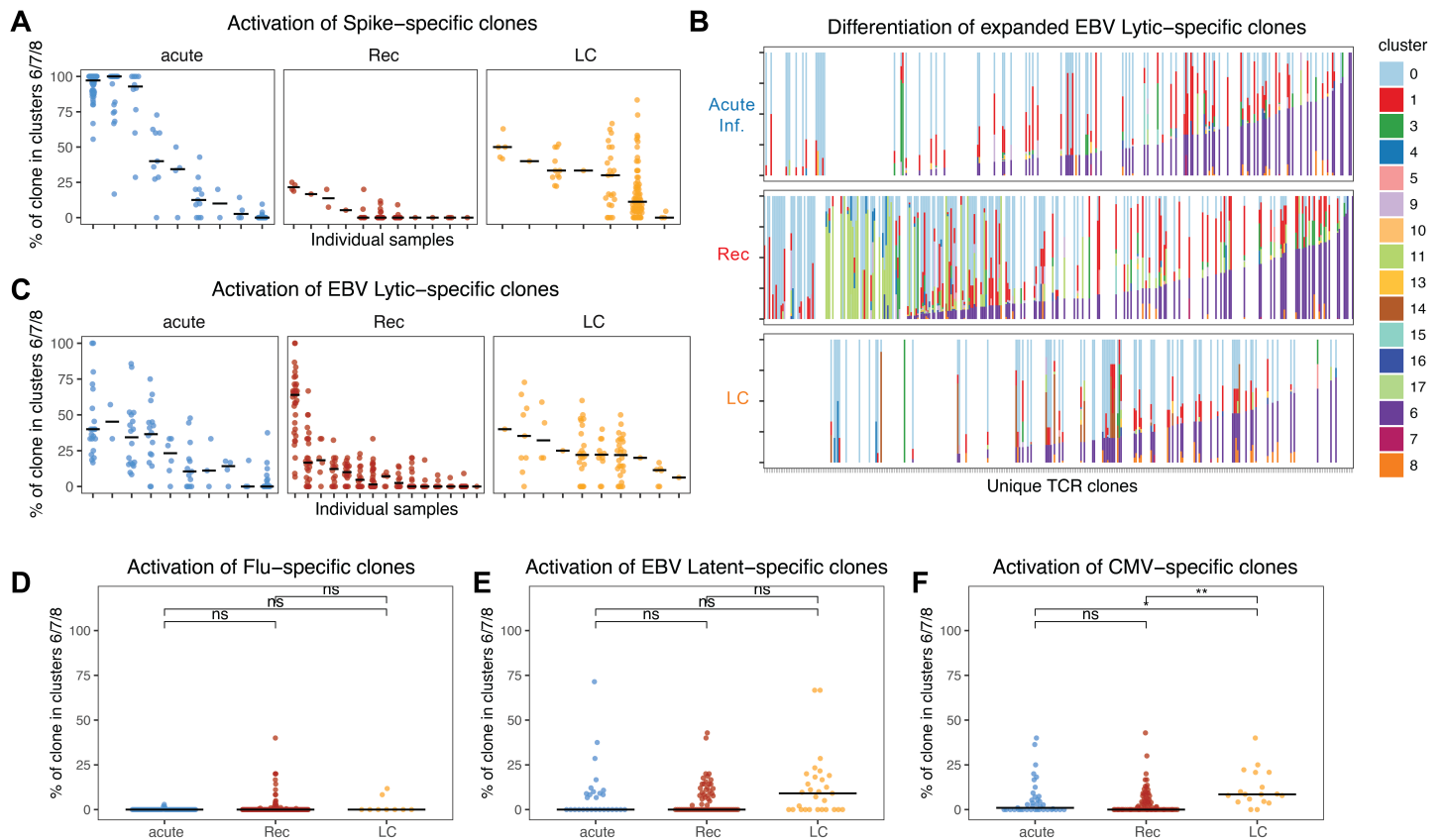

**Fig. S17: Activation of antigen-specific CD8 T cell clones.**

(A) Summary data for the percent of each spike-specific CD8 T cell clone in activated clusters 6, 7, or 8, split by group and individual samples. Each point represents a unique clone, and only clones with at least 5 cells were included. (B) Ranked ordering by percent of cells falling in activated clusters 6, 7, or 8 for all EBV lytic-specific CD8 T cell clones with at least 5 cells. Each column represents a unique clone, ranked along the x-axis and colored by the proportion of cells assigned to each cluster. (C) Summary data for the percent of each EBV lytic-specific CD8 T cell clone in activated clusters 6, 7, or 8, split by group and individual samples. Each point represents a unique clone, and only clones with at least 5 cells were included. (D-F) Summary data for the percent of each (D) Flu-, (E) EBV latent-, and (F) CMV-specific CD8 T cell clone in activated clusters 6, 7, or 8. Each point represents a unique clone, and only clones with at least 5 cells were included.

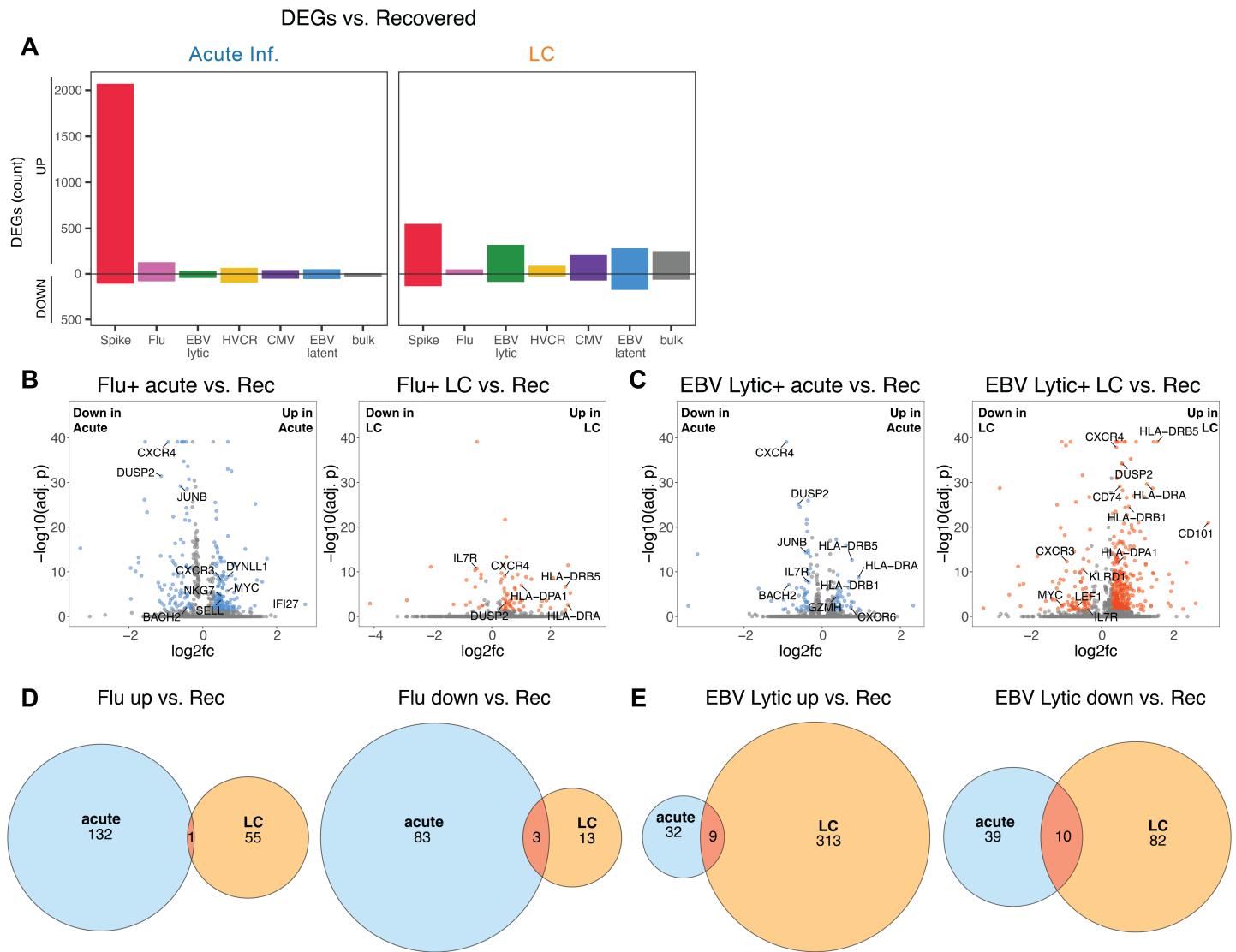

**Fig. S18: Differential gene expression analyses comparing CD8 T cells of different viral specificities from acute infection or LC to cells from recovered controls.**

(A) Summary data displaying the count of significantly differentially expressed genes comparing cells from acute infection (left) or LC (right) to cells from recovered controls for the indicated specificities. (B-C) Volcano plots displaying significantly upregulated and downregulated DEGs in (B) Flu-specific and (C) EBV lytic-specific CD8 T cells from acute infection or LC compared to recovered controls. Significant DEGs were identified via the Seurat FindMarkers function. (D-E) Venn diagrams displaying the shared and distinct significantly upregulated (left) and downregulated (right) genes from (D) Flu-specific and (E) EBV lytic-specific CD8 T cells from acute infection and LC versus recovered controls. All analyses were performed after downsampling each antigen-specific population to a maximum of 150 cells per sample to avoid bias associated with highly expanded responses. Significant DEGs were defined as those with an adjusted p value below 0.05 and log2fc above 0.3 or below -0.3.

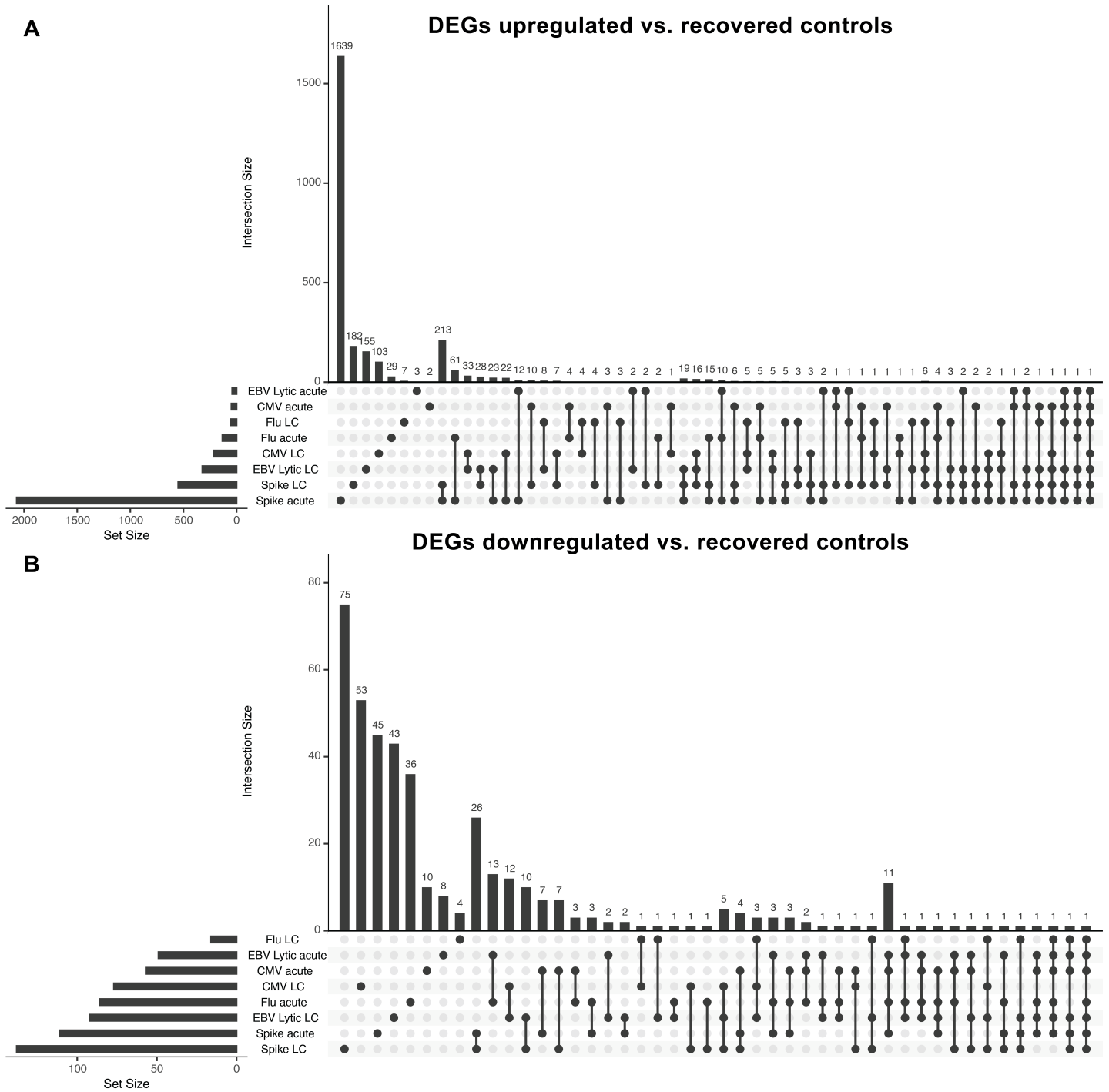

**Fig. S19: Shared and distinct DEGs in virus-specific CD8 T cell populations from acute infection or LC compared to recovered controls.**

(A-B) UpSet plots comparing shared and distinct differentially (A) upregulated and (B) downregulated genes from spike-, EBV lytic-, CMV-, and Flu-specific CD8 T cell populations from acute infection or LC compared to recovered controls. Significant DEGs were defined as those with an adjusted p value below 0.05 and log2fc above 0.3 or below -0.3.

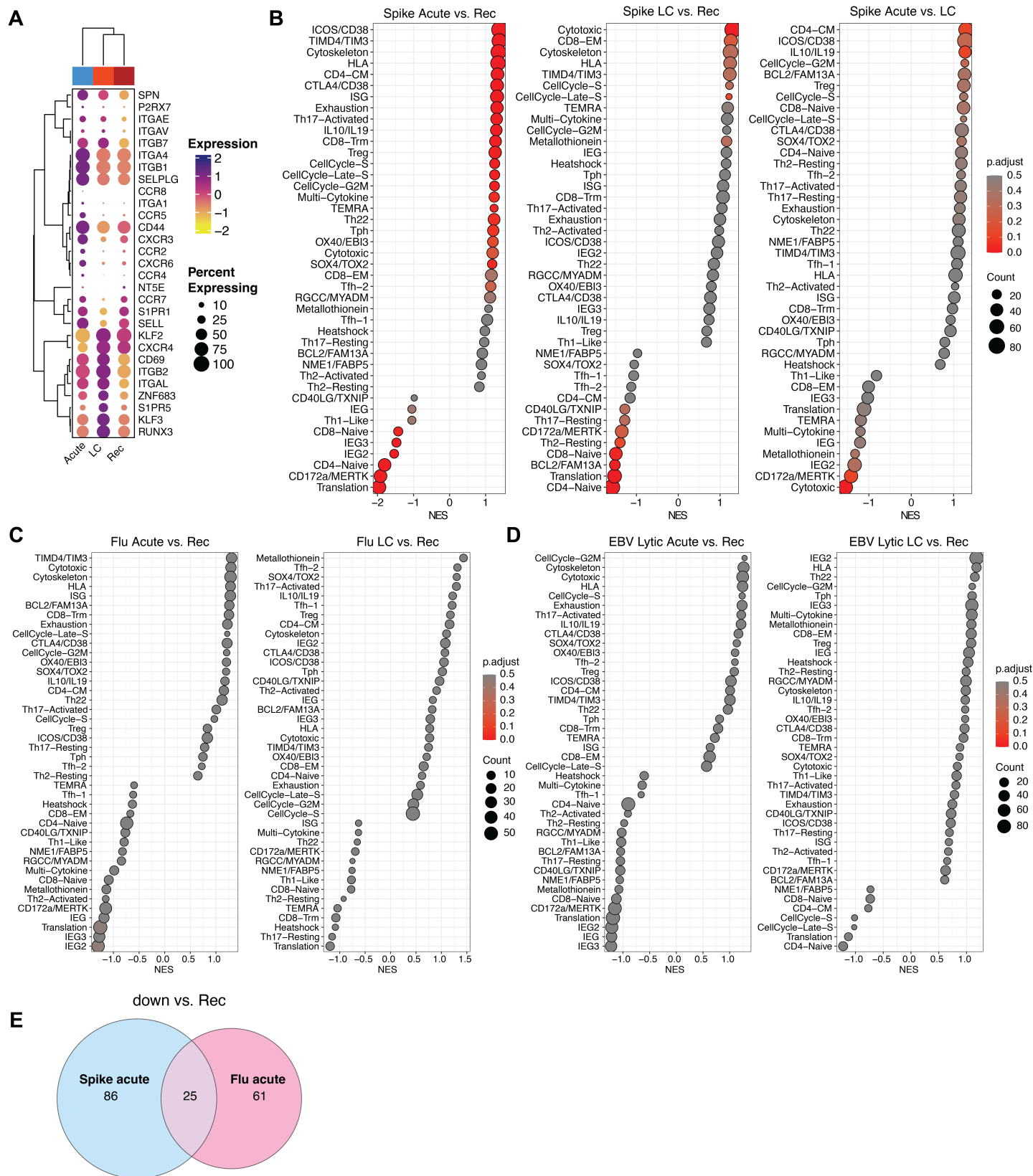

**Fig. S20: Gene expression programs enriched in spike-, Flu-, and EBV lytic-specific CD8 T cells from acute infection and LC.**

**(A)** Clustered dot plot displaying the expression of selected genes associated with CD8 T cell trafficking. **(B)** Gene set enrichment analyses using the gene lists for each gene expression program (GEP) from T-CellAnnoTator (TCAT)(64) to test for enrichment within ranked gene expression lists for spike-specific CD8 T

cells from acute infection or LC compared to recovered controls, or from acute infection compared to LC (right). Significantly enriched gene sets are colored red and have high NES values, significantly depleted gene sets are colored red and have low NES values. Gene sets in grey are not significantly enriched in that cluster. **(C-D)** Gene set enrichment analyses using the gene lists for each TCAT GEP to test for enrichment within ranked gene expression lists for **(C)** Flu-specific and **(D)** EBV lytic-specific CD8 T cells from acute infection or LC compared to recovered controls. **(E)** Venn diagram displaying the shared and distinct significantly downregulated genes for spike- and Flu-specific CD8 T cells from acute infection compared to recovered controls. Along with upregulated genes (Fig. 5E), these genes were used to derive signatures of acute TCR stimulation and bystander stimulation as indicated in Fig. 5E. All analyses were performed after downsampling each antigen-specific population to a maximum of 150 cells per sample to avoid bias associated with highly expanded responses.

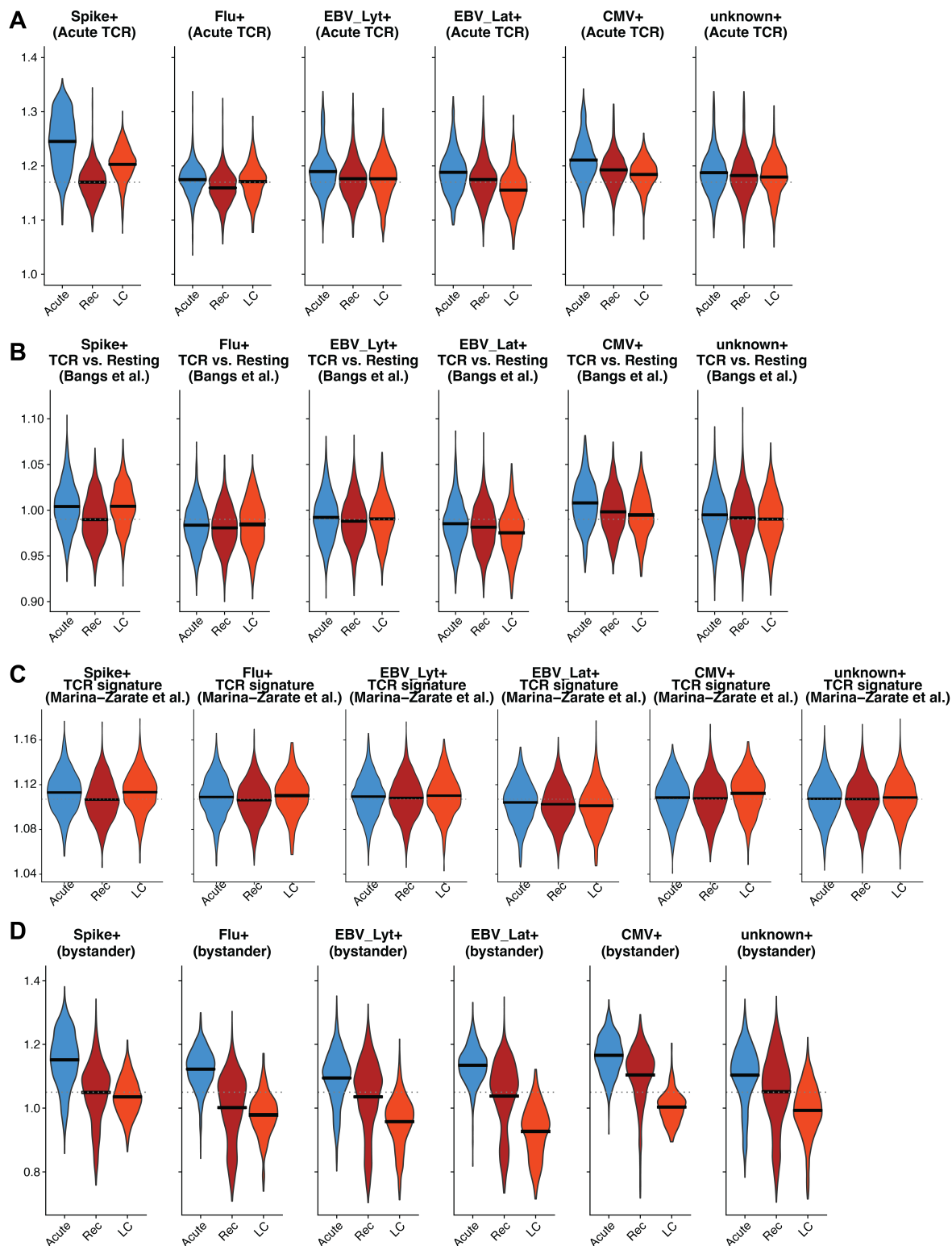

**Fig. S21: Module score enrichment for acute TCR stimulation or bystander activation signatures for all specificities by group.**

(A-D) Summary data depicting module scores for virus-specific CD8 T cells from the indicated groups. Module scores were calculated using the UCell package. For (A) and (D), upregulated and downregulated modules were derived separately using the DEGs comparing Spike- and Flu-specific CD8 T cells during acute infection (as described in Fig. 5E). These were then combined into a single module by subtracting the downregulated score from the upregulated score after normalizing by the number of genes in each module. For (B-C), genes that are upregulated after TCR stimulation in two distinct published signatures were used to derive the score(66, 67).

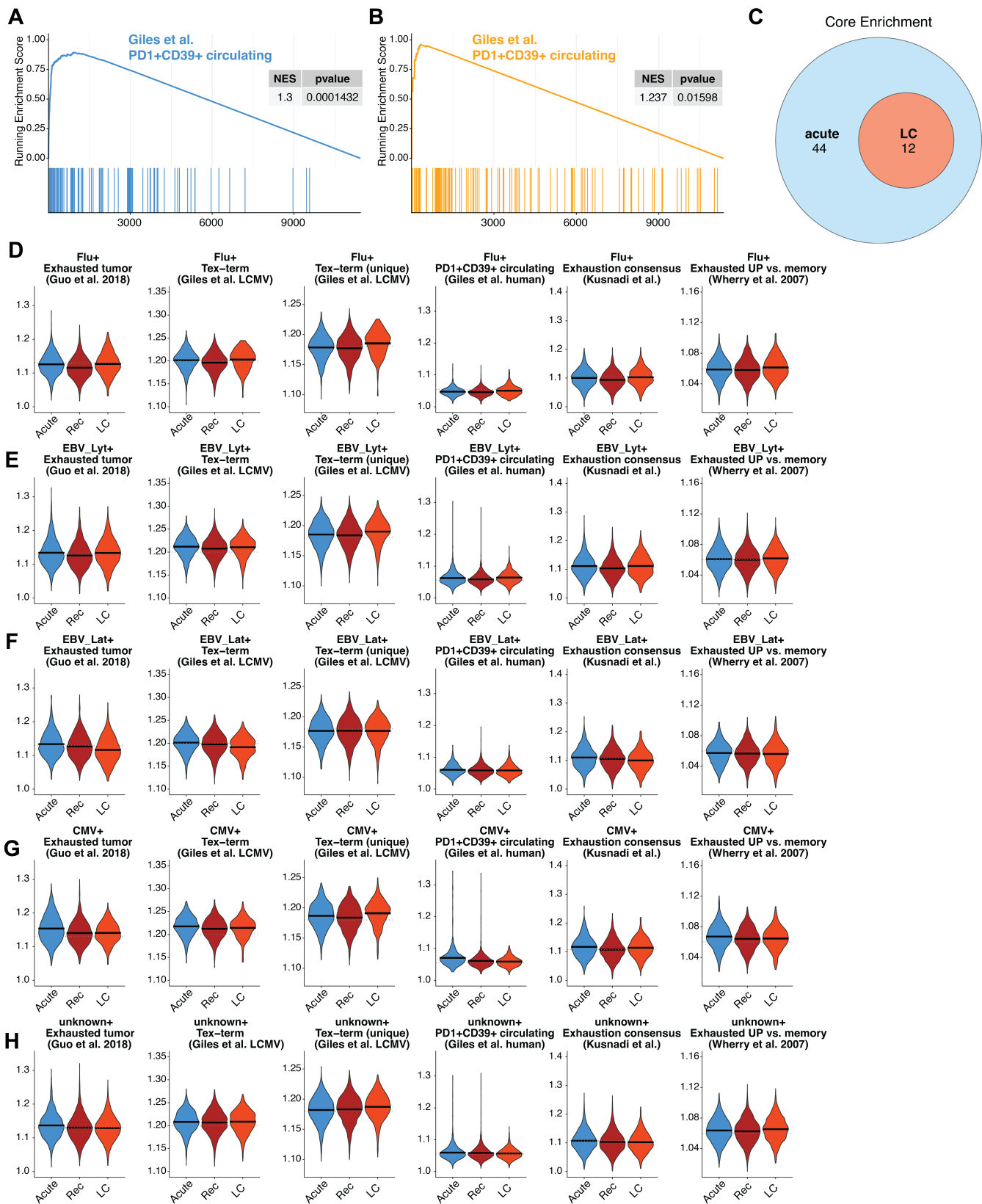

**Fig. S22: Module score enrichment for T cell exhaustion signatures for all specificities by group.**

(A-B) Gene set enrichment analysis of spike-specific CD8 T cells from (A) acute infection or (B) LC compared to recovered controls, using a published gene set of human circulating PD-1+ CD39+ CD8 T cells(69). (C) Venn diagram depicting the overlap in genes driving the core enrichment for the gene set in (A-B) for spike-specific CD8 T cells from acute infection or LC compared to recovered controls. (D-H) Summary data depicting module scores for **(D)** Flu-specific, **(E)** EBV lytic-specific, **(F)** EBV latent-specific, **(G)** CMV-specific, and **(H)** bulk CD8 T cells from the indicated groups. Module scores were calculated using the UCell package using six distinct published gene sets of T cell exhaustion, including genes upregulated in exhausted T cells from non-small cell lung cancer tumors(68), terminally exhausted CD8 T cells from LCMV clone 13 infection compared to all other cells(70), terminally exhausted CD8 T cells from LCMV clone 13 infection and not upregulated in effector CD8 T cells(70), circulating PD-1+ CD39+ CD8 T cells in humans(69), a consensus signature of T cell exhaustion(71), and a signature of exhausted T cells compared to memory T cells in LCMV(72). All analyses were performed after downsampling each antigen-specific population to a maximum of 150 cells per sample to avoid bias associated with highly expanded responses.

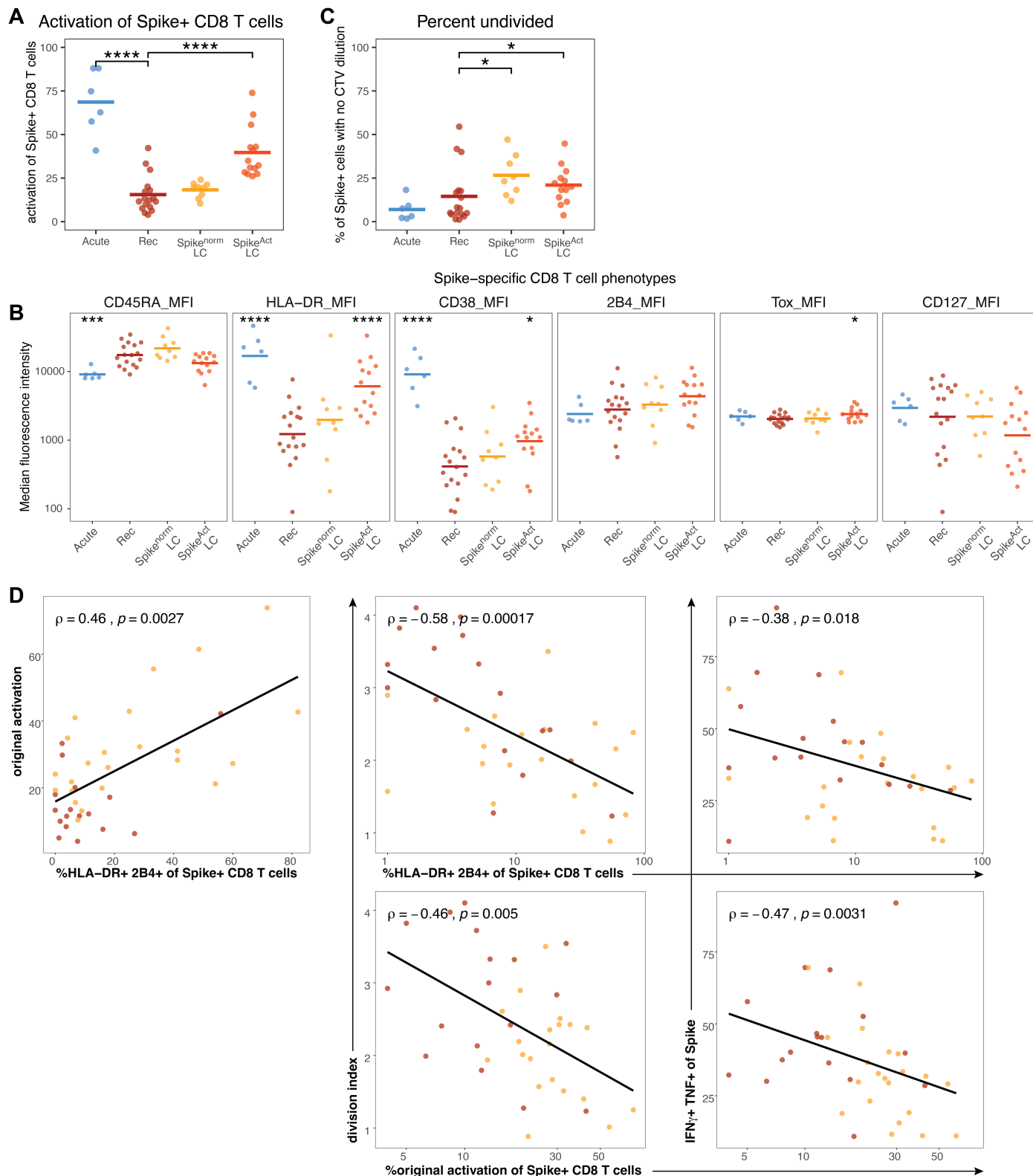

**Fig. S23: Supplementary information relating to spike-specific CD8 T cell functional assays.**

(A) Summary data describing the activation of spike-specific CD8 T cells in the samples selected for additional profiling and spike-specific CD8 T cell functional assays, with LC samples separated based on the level of T cell activation detected in Fig. 2B. (B) Summary data depicting the median fluorescence intensity (MFI) of the indicated markers on spike-specific CD8 T cells from each group. (C) Summary data depicting the percent of spike-specific CD8 T cells that were undivided (no dilution of CTV) after 96 hours of stimulation with spike peptides. **(D)** Representative Spearman rank correlations of baseline spike-specific CD8 T cell differentiation states (as a percentage of the spike-specific CD8 T cell population) and functional assay measurements, as included in the matrix in Fig. 6N. All recovered and LC samples were included in correlation analyses. Baseline expression of HLA-DR and 2B4 was correlated with original activation measurements in Fig. 2B, and both measures were independently correlated with division index and the percent of spike-specific CD8 T cells producing IFNγ and TNF. Statistical tests were Wilcoxon tests with Benjamini-Hochberg correction for multiple comparisons. \*$p < 0.05$, \*\*$p < 0.01$, \*\*\*$p < 0.001$, \*\*\*\*$p < 0.0001$. In the absence of brackets, all statistical tests are comparisons to recovered controls.

**Table S1: LIINC cohort samples (UCSF) for CD8 T cell analyses in Fig. 1-3**

| **Characteristic** | **LC**<br>**N = 107<sup>1</sup>** | **Recovered**<br>**N = 37<sup>1</sup>** |
| --- | --- | --- |
| **age (years)** | 47.2 (25 - 71) | 45.0 (27 - 85) |
| **sex** |  |  |
| Female | 69 (64%) | 11 (30%) |
| Male | 38 (36%) | 26 (70%) |
| **ethnicity** |  |  |
| Hispanic/Latino | 21 (20%) | 3 (8.1%) |
| Not Hispanic/Latino | 86 (80%) | 34 (92%) |
| **race** |  |  |
| Asian | 6 (6.5%) | 0 (0%) |
| Native American | 2 (2.2%) | 0 (0%) |
| White | 84 (91%) | 34 (100%) |
| Unknown/Other | 15 | 3 |
| **first infection date** |  |  |
| March 2020 - April 2021 | 90 (84%) | 31 (84%) |
| May 2021 - November 2021 | 13 (12%) | 6 (16%) |
| >=December 2021 | 4 (3.7%) | 0 (0%) |
| **n vaccine doses** | 1.0 (0 - 4) | 1.1 (0 - 3) |
| **n SARS-CoV-2 infections** | 1.0 (1 - 2) | 1.1 (1 - 2) |
| **days since last vaccine/infection** | 160.4 (46 - 400) | 146.1 (48 - 366) |
| **hospitalized with COVID-19** | 22 (21%) | 1 (2.7%) |
| **days since symptom onset** | 243.8 (92 - 419) | 251.9 (108 - 481) |
| **BMI** | 29.9 (20 - 51) | 25.2 (18 - 33) |
| **BMI category** |  |  |
| 25 to 30kg/m2 | 26 (24%) | 13 (35%) |
| <25kg/m2 | 30 (28%) | 17 (46%) |
| > 30kg/m2 | 51 (48%) | 7 (19%) |
| **autoimmune disease** | 7 (6.5%) | 0 (0%) |
| **cancer** | 7 (6.5%) | 0 (0%) |

| Characteristic | LC<br>N = 107<sup>1</sup> | Recovered<br>N = 37<sup>1</sup> |
| --- | --- | --- |
| diabetes | 10 (9.3%) | 1 (2.7%) |
| hypertension | 15 (14%) | 1 (2.7%) |
| lung disease | 27 (25%) | 7 (19%) |
| kidney disease | 3 (2.8%) | 0 (0%) |

<sup>1</sup>Mean (Min - Max); n (%)

N = number of samples

**Table S2: COVEND and AALC cohort samples (University of Pennsylvania) for CD8 T cell analyses in Fig. 1-3**

| Characteristic | LC<br>N = 44<sup>1</sup> | Recovered<br>N = 47<sup>1</sup> |
| --- | --- | --- |
| **age (years)** | 51.0 (22 - 80) | 30.4 (22 - 54) |
| **sex** |  |  |
| Female | 32 (73%) | 28 (60%) |
| Male | 12 (27%) | 19 (40%) |
| **ethnicity** |  |  |
| Hispanic/Latino | 2 (4.5%) | 1 (2.1%) |
| Not Hispanic/Latino | 42 (96%) | 46 (98%) |
| **race** |  |  |
| Asian | 0 (0%) | 5 (11%) |
| Black | 3 (7.3%) | 1 (2.1%) |
| Native American | 0 (0%) | 1 (2.1%) |
| White | 38 (93%) | 40 (85%) |
| Unknown/Other | 3 | 0 |
| **first infection date** |  |  |
| March 2020 - April 2021 | 18 (44%) | 4 (8.5%) |
| May 2021 - November 2021 | 1 (2.4%) | 1 (2.1%) |
| >=December 2021 | 22 (54%) | 42 (89%) |
| Unknown | 3 | 0 |
| **n vaccine doses** | 3.3 (0 - 6) | 4.0 (2 - 5) |
| Unknown | 3 | 0 |
| **n SARS-CoV-2 infections** | 1.5 (1 - 3) | 1.5 (1 - 3) |
| Unknown | 3 | 0 |
| **days since last vaccine/infection** | 247.9 (79 - 627) | 226.9 (45 - 641) |
| **hospitalized with COVID-19** |  |  |
| No | 43 (100%) | 47 (100%) |
| **days since symptom onset** | 605.3 (201 - 1,196) | 369.1 (106 - 1,021) |

<sup>1</sup>Mean (Min - Max); n (%)

N = number of samples

**Table S3: MY-LC cohort samples (Yale/Mt. Sinai) for CD8 T cell analyses in Fig. 1-3**

| Characteristic | Study Cohorts |  |
| --- | --- | --- |
|  | LC<br>N = 45<sup>1</sup> | Recovered<br>N = 12<sup>1</sup> |
| **age (years)** | 45.4 (26 - 79) | 45.3 (30 - 68) |
| **sex** |  |  |
| Female | 33 (73%) | 7 (58%) |
| Male | 12 (27%) | 5 (42%) |
| **ethnicity** |  |  |
| Hispanic/Latino | 1 (2.2%) | 3 (25%) |
| Not Hispanic/Latino | 44 (98%) | 9 (75%) |
| **race** |  |  |
| Asian | 1 (2.5%) | 0 (0%) |
| Black | 3 (7.5%) | 2 (20%) |
| White | 36 (90%) | 8 (80%) |
| Unknown/Other | 5 | 2 |
| **first infection date** |  |  |
| March 2020 - April 2021 | 45 (100%) | 10 (83%) |
| May 2021 - November 2021 | 0 (0%) | 1 (8.3%) |
| >=December 2021 | 0 (0%) | 1 (8.3%) |
| **n vaccine doses** | 1.5 (0 - 2) | 2.4 (2 - 3) |
| **n SARS-CoV-2 infections** | 1.0 (1 - 1) | 1.0 (1 - 1) |
| **days since last vaccine/infection** | 171.5 (49 - 578) | 187.1 (92 - 323) |
| **hospitalized with COVID-19** | 7 (16%) | 0 (0%) |
| **days since symptom onset** | 399.2 (104 - 578) | 427.0 (88 - 705) |
| **BMI** | 26.4 (16 - 55) | 24.6 (19 - 31) |

<sup>1</sup>Mean (Min - Max); n (%)

N = number of participants (1 sample each)

**Table S4: List of HLA-I and HLA-II tetramers**

| HLA | Virus/antigen category | Peptide | Fluorophore1 | Fluorophore2 | BEAM1 | BEAM2 | Used in Fig. 6 |
| --- | --- | --- | --- | --- | --- | --- | --- |
| HLA-A*02:01 | Non-spike SARS-CoV-2 (Replicase) | ALWEIQQVV | BV786 | PE-Fire700 |  |  |  |
| HLA-A*02:01 | Non-spike SARS-CoV-2 (Replicase) | FLAHIQWMV | BV786 | PE-Fire700 |  |  |  |
| HLA-A*02:01 | Non-spike SARS-CoV-2 (Replicase) | FLLNKEMYL | BV786 | PE-Fire700 |  |  |  |
| HLA-A*02:01 | Non-spike SARS-CoV-2 (Replicase) | KLWAQCVQL | BV786 | PE-Fire700 |  |  |  |
| HLA-A*02:01 | Non-spike SARS-CoV-2 (Replicase) | SMWALIISV | BV786 | PE-Fire700 |  |  |  |
| HLA-A*02:01 | Non-spike SARS-CoV-2 (Replicase) | YLATALLTL | BV786 | PE-Fire700 |  |  |  |
| HLA-A*02:01 | Non-spike SARS-CoV-2 (Nucleoprotein) | LALLLLDRL | BV786 | PE-Fire700 |  |  |  |
| HLA-A*02:01 | Non-spike SARS-CoV-2 (Nucleoprotein) | GMSRIGMEV | BV786 | PE-Fire700 |  |  |  |
| HLA-A*02:01 | Non-spike SARS-CoV-2 (Nucleoprotein) | LLLDRLNQL | BV786 | PE-Fire700 |  |  |  |
| HLA-A*02:01 | Non-spike SARS-CoV-2 (ORF3a) | ALSKGVHFV | BV786 | PE-Fire700 |  |  |  |
| HLA-A*02:01 | Non-spike SARS-CoV-2 (ORF3a) | LLYDANYFL | BV786 | PE-Fire700 |  |  |  |
| HLA-A*02:01 | Spike (SARS-CoV-2) | ALNTLVKQL | BB515 | PE |  |  |  |
| HLA-A*02:01 | Spike (SARS-CoV-2) | KIADYNYKL | BB515 | PE | 1 | 7 | yes |
| HLA-A*02:01 | Spike (SARS-CoV-2) | RLQSLQTYV | BB515 | PE | 1 | 8 | yes |
| HLA-A*02:01 | Spike (SARS-CoV-2) | SIIAYTMSL | BB515 | PE |  |  |  |
| HLA-A*02:01 | Spike (SARS-CoV-2) | VVFLHVTYV | BB515 | PE |  |  | yes |
| HLA-A*02:01 | Spike (SARS-CoV-2) | FIAGLIAIV | BB515 | PE |  |  |  |
| HLA-A*02:01 | Spike (SARS-CoV-2) | NLNESLIDL | BB515 | PE |  |  |  |
| HLA-A*02:01 | Spike (SARS-CoV-2) | RLNEVAKNL | BB515 | PE |  |  |  |
| HLA-A*02:01 | Spike (SARS-CoV-2) | TLDSKTQSL | BB515 | PE |  |  |  |
| HLA-A*02:01 | Spike (SARS-CoV-2) | VLNDILSRL | BB515 | PE | 1 | 9 | yes |
| HLA-A*02:01 | Spike (SARS-CoV-2) | YLQPRTFLL | BB515 | PE | 1 | 10 | yes |
| HLA-A*02:01 | CMV | NLVPMVATV | BV650 | BV786 | 2 | 11 |  |
| HLA-A*02:01 | CMV | VLEETSVML | BV650 | BV786 | 2 | 12 |  |
| HLA-A*02:01 | Flu | AIMDKNIIL | BV650 | PE | 3 | 13 |  |
| HLA-A*02:01 | Flu | GILGFVFTL | BV650 | PE | 3 | 14 |  |
| HLA-A*02:01 | Flu | FMYSDFHFI | BV650 | PE | 3 | 15 |  |
| HLA-A*02:01 | Flu | NMLSTVLGV | BV650 | PE | 3 | 16 |  |
| HLA-A*02:01 | EBV Lytic | YVLDHLIVV | BV650 | BB515 | 4 | 7 |  |
| HLA-A*02:01 | EBV Lytic | GLCTLVAML | BV650 | BB515 | 4 | 8 |  |
| HLA-A*02:01 | EBV Latent | CLGGLLTMV | BV786 | BB515 | 5 | 9 |  |
| HLA-A*02:01 | EBV Latent | FLYALALLL | BV786 | BB515 | 5 | 10 |  |
| HLA-A*02:01 | EBV Latent | YLLEMLWRL | BV786 | BB515 | 5 | 11 |  |
| HLA-A*02:01 | EBV Latent | LLDFVRFMGV | BV786 | BB515 | 5 | 12 |  |
| HLA-A*02:01 | EBV Latent | YLQQNWWTL | BV786 | BB515 | 5 | 13 |  |
| HLA-A*02:01 | HVCR | ILIEGIFFV | BV786 | PE | 6 | 14 |  |
| HLA-A*03:01 | Non-spike SARS-CoV-2 (Replicase) | KLFAAETLK | BV786 | PE-Fire700 |  |  |  |
| HLA-A*03:01 | Non-spike SARS-CoV-2 (Replicase) | KLFDRYFKY | BV786 | PE-Fire700 |  |  |  |
| HLA-A*03:01 | Non-spike SARS-CoV-2 (Replicase) | KTIQPRVEK | BV786 | PE-Fire700 |  |  |  |
| HLA-A*03:01 | Non-spike SARS-CoV-2 (Replicase) | VTNNTFTLK | BV786 | PE-Fire700 |  |  |  |
| HLA-A*03:01 | Non-spike SARS-CoV-2 (Nucleoprotein) | KTFPPTEPK | BV786 | PE-Fire700 |  |  |  |
| HLA-A*03:01 | Spike (SARS-CoV-2) | GVYFASTEK | BB515 | PE | 7 | 15 | yes |
| HLA-A*03:01 | Spike (SARS-CoV-2) | KCYGVSPTK | BB515 | PE | 7 | 16 | yes |
| HLA-A*03:01 | CMV | KLGGALQAK | BV650 | BV786 | 2 | 7 |  |
| HLA-A*03:01 | CMV | RIKEHMLKK | BV650 | BV786 | 2 | 8 |  |
| HLA-A*03:01 | Flu | ILRGSVAHK | BV650 | PE | 3 | 9 |  |
| HLA-A*03:01 | EBV Lytic | RVRAYTYSK | BV650 | BB515 | 4 | 10 |  |
| HLA-A*03:01 | EBV Latent | RLRAEAQVK | BV786 | BB515 | 11 | 14 |  |
| HLA-A*03:01 | VZV | SVYFWWLNR | BV786 | PE | 6 | 11 |  |
| HLA-A*03:01 | VZV | AVYENPLSVEK | BV786 | PE | 6 | 12 |  |
| DPB1*04:01 | Spike (SARS-CoV-2) | TFEYVSQPFLMDLE | PE | APC |  |  |  |
| DRB1*15:01 | Spike (SARS-CoV-2) | NLLLQYGSFCTQLNRAL | PE | APC |  |  |  |

**Table S5: OUTSMART-LC clinical trial samples**

| **Characteristic** | **Study Cohorts** |
| --- | --- |
|  | **LC**<br>**N = 32 participants<sup>1</sup>** |
| **age (years)** | 43.4 (23 - 73) |
| Unknown | 1 |
| **sex** |  |
| Female | 17 (55%) |
| Male | 14 (45%) |
| Unknown | 1 |
| **ethnicity** |  |
| Hispanic/Latino | 3 (9.7%) |
| Not Hispanic/Latino | 28 (90%) |
| Unknown | 1 |
| **race** |  |
| Asian | 4 (13%) |
| Black | 1 (3.3%) |
| Native American | 1 (3.3%) |
| White | 24 (80%) |
| Unknown/Other | 2 |
| **first infection date** |  |
| March 2020 - April 2021 | 11 (35%) |
| May 2021 - November 2021 | 2 (6.5%) |
| >=December 2021 | 18 (58%) |
| Unknown | 1 |
| **hospitalized with COVID-19** | 2 (6.5%) |
| Unknown | 1 |

<sup>1</sup>Mean (Min - Max); n (%)

**Table S6: Key Resources Table**

| REAGENT or RESOURCE | SOURCE | IDENTIFIER |
| --- | --- | --- |
| **Antibodies** |  |  |
| BUV395 CD27 (anti-human) | BD Biosciences | Cat#563815 |
| BUV496 CD4 (anti-human) | BD Biosciences | Cat#612936 |
| BUV615 CD8 (anti-human) | BD Biosciences | Cat#751518 |
| BUV661 CD38 (anti-human) | BD Biosciences | Cat#612969 |
| BUV805 HLA-DR (anti-human) | BD Biosciences | Cat#748338 |
| BUV737 CCR6 (anti-human) | BD Biosciences | Cat#612780 |
| BV421 CXCR3 (anti-human) | Biolegend | Cat#353716 |
| BV570 CD3 (anti-human) | Biolegend | Cat#300436 |
| BV605 CD45RA (anti-human) | Biolegend | Cat#304134 |
| BV650 CD25 (anti-human) | BD Biosciences | Cat#563719 |
| BV786 CD71 (anti-human) | BD Biosciences | Cat#563768 |
| RB545 CD103 (anti-human) | BD Biosciences | Cat#756639 |
| RB613 PD-1 (anti-human) | BD Biosciences | Cat#571094 |
| BV480 Ki67 (anti-human) | BD Biosciences | Cat#566172 |
| AF488 TCF1 (anti-human) | Cell Signaling | Cat#6444 |
| BB700 CTLA4 (anti-human) | BD Biosciences | Cat#566902 |
| RB780 Granzyme K (anti-human) | BD Biosciences | Cat#569225 |
| PE-Cy5.5 FoxP3 (anti-human) | Invitrogen | Cat#35-4776-42 |
| RB744 CD127 (anti-human) | BD Biosciences | Cat#570607 |
| Spark NIR 685 CCR7 (anti-human) | Biolegend | Cat#353258 |
| APC/Fire 750 ICOS (anti-human) | Biolegend | Cat#313536 |
| APC/Fire 810 CCR4 (anti-human) | Biolegend | Cat#359440 |
| APC-R700 CXCR5 (anti-human) | BD Biosciences | Cat#565191 |
| PE/Fire 810 Tim-3 (anti-human) | Biolegend | Cat#345059 |
| RB780 CD127 (anti-human) | BD | Cat#568750 |
| AF647 TCF1 (anti-human) | CellSignaling | Cat#6709S |
| AF700 CCR7 (anti-human) | Biolegend | Cat#353244 |
| APC/Fire750 KLRG1 (anti-human) | Biolegend | Cat#367718 |
| BUV395 CD45RA (anti-human) | BD | Cat#740298 |
| BUV496 CD8 (anti-human) | BD | Cat#612942 |
| BUV661 CX3CR1 (anti-human) | BD | Cat#750690 |
| BUV737 CD27 (anti-human) | BD | Cat#612830 |
| BUV805 CD3 (anti-human) | BD | Cat#612896 |
| BV421 PD-1 (anti-human) | Biolegend | Cat#329920 |
| BV480 CD49d (anti-human) | BD | Cat#566134 |
| V500 CD14 (anti-human) | BD | Cat#561391 |
| V500 CD16 (anti-human) | BD | Cat#561394 |
| V500 CD19 (anti-human) | BD | Cat#561121 |
| V500 CD41a (anti-human) | BD | Cat#563250 |
| BV605 HLA-DR (anti-human) | Biolegend | Cat#307640 |
| BV711 Ki67 (anti-human) | Biolegend | Cat#350516 |
| BV750 CD38 (anti-human) | BD | Cat#746909 |
| PE-Cy5 CD95 (anti-human) | BD | Cat#561977 |
| PE-Cy5.5 CD4 (anti-human) | Invitrogen | Cat#35-0047-42 |
| PE-Cy7 T-bet (anti-human) | Biolegend | Cat#644824 |
| PE-TexasRed Granzyme B (anti-human) | Invitrogen | Cat#GRB17 |
| BUV563 CD39 (anti-human) | BD | Cat#748473 |
| Pacific Blue Perforin | Biolegend | Cat#353305 |
| BV480 HLA-DR | BD | Cat#566154 |
| BV570 CD4 | Biolegend | Cat# 317445 |
| BV605 Tbet | Biolegend | Cat# 644817 |
| BV828 Granzyme B | BD | Cat# 572658 |
| RB670 CD95 | BD | Cat# 572261 |
| RB705 LAG3 | BD | Cat# 570636 |
| PE-Cy5 CD244 (2B4) | Biolegend | Cat#329539 |
| APC TOX | Miltenyi | Cat# 130-118-335 |
| R718 TCF1 | BD | Cat# 567587 |
| APC/Fire 750 CCR7 | Biolegend | Cat# 353246 |
| APC/Fire 810 KLRG1 | Biolegend | Cat# 367732 |
| PE-CF594 Blimp-1 | BD | Cat#565274 |
| RY655 CD101 | BD | Cat# 773706 |
| PE-Cy7 CD160 | Biolegend | Cat# 341212 |
| Spark NIR 685 CD69 | Biolegend | Cat# 310957 |
| BV480 CD69 | BD | Cat# 747519 |
| IL-2 BV750 | BD | Cat# 566361 |
| RY775 TNF | BD | Cat# 571664 |
| Spark NIR 685 IFNγ | Biolegend | Cat# 502552 |
| BV750 CD4 | BD | Cat# 750987 |
| PE/Fire 640 CD25 | Biolegend | Cat# 356148 |
| TotalSeq-C anti-human Hashtag antibodies | Biolegend | e.g. Cat#394661 |

#### Chemicals, peptides, and recombinant proteins

|  |  |  |
| --- | --- | --- |
| BV421 Streptavidin | Biolegend | Cat#405226 |
| BV650 Streptavidin | BD Biosciences | Cat#563262 |
| BV786 Streptavidin | BD Biosciences | Cat#563858 |
| BUV615 Streptavidin | BD Biosciences | Cat#613013 |
| BB515 Streptavidin | BD Biosciences | Cat#564453 |
| PE Streptavidin | Biolegend | Cat#405203 |
| PE/Fire 700 Streptavidin | Biolegend | Cat#405206 |
| APC Streptavidin | Biolegend | Cat#405207 |
| CellTrace Violet | Thermo Fisher | Cat#C34571 |
| Zombie NIR Viability Dye | Biolegend | Cat#423106 |
| Ghost Viability Dye Violet 510 | Tonbo | Cat#13-0870-T100 |
| Human TruStain FcX (Fc Receptor Blocking Solution) | Biolegend | Cat#422302 |
| GolgiStop (Containing Monensin) | BD Biosciences | Cat#51-2092K7 |
| Anti-Human CD28/CD49d Purified | BD Biosciences | Cat#347690 |
| Foxp3 / Transcription Factor Fixation/Permeabilization Concentrate and Diluent | eBioscience | Cat#00-5521-00 |
| Peptides used for tetramers or stimulation assay (Table 3) | Genscript | Custom synthesis |
| Dasatinib | Cell Signaling Technology | 9052S |
| Flex-T™ HLA-A*02:01 Monomer UVX | BioLegend | Cat#280004 |
| Flex-T™ HLA-A*03:01 Monomer UVX | BioLegend | Cat#280005 |
| Chromium, Single Cell 5' BEAM Core Kit, PE, Set A 128 rxns | 10X Genomics | Cat# 1000539 |
| Chromium Next GEM Single Cell 5' HT Kit v2, 8 rxns | 10X Genomics | Cat# 1000374 |
| HLA-II tetramer TFEYVSQPFLMDLE; Spike 167 - 180; DPB1*04:01 (PE and APC) | NIH Tetramer Core Facility |  |
| HLA-II tetramer NLLLQYGSFCTQLNRAL; Spike 751 - 767; DRB1*15:01 (PE and APC) | NIH Tetramer Core Facility |  |
| Brilliant Stain Buffer | BD | Cat# 566349 |
